# Limits and Convergence properties of the Sequentially Markovian Coalescent

**DOI:** 10.1101/2020.07.23.217091

**Authors:** Thibaut Sellinger, Diala Abu Awad, Aurélien Tellier

**Affiliations:** Professorship for Population Genetics, Department of Life Science Systems – Technical University of Munich, Germany

**Keywords:** Hidden Markov Model, Ancestral Recombination Graph, Population Genetics

## Abstract

Many methods based on the Sequentially Markovian Coalescent (SMC) have been and are being developed. These methods make use of genome sequence data to uncover population demographic history. More recently, new methods have extended the original theoretical framework, allowing the simultaneous estimation of the demographic history and other biological variables. These methods can be applied to many different species, under different model assumptions, in hopes of unlocking the population/species evolutionary history. Although convergence proofs in particular cases have been given using simulated data, a clear outline of the performance limits of these methods is lacking. We here explore the limits of this methodology, as well as present a tool that can be used to help users quantify what information can be confidently retrieved from given datasets. In addition, we study the consequences for inference accuracy violating the hypotheses and the assumptions of SMC approaches, such as the presence of transposable elements, variable recombination and mutation rates along the sequence and SNP call errors. We also provide a new interpretation of the SMC through the use of the estimated transition matrix and offer recommendations for the most efficient use of these methods under budget constraints, notably through the building of data sets that would be better adapted for the biological question at hand.

## Introduction

Recovering the demographic history of a population has become a central theme in evolutionary biology. The demographic history (the variation of effective population size over time) is linked to environmental and demographic changes that existing and/or extinct species have experienced (population expansion, colonization of new habitats, past bottlenecks) (Bergstrom et al., 2020; Gaut et al., 2018; Palkopoulou, Lipson, et al., 2018). Current statistical tools to estimate the demographic history rely on genomic data (Schraiber and Akey, 2015) and these inferences are often linked to archaeological or climatic data, providing novel insights on the evolutionary history (Barroso et al., 2019; Fulgione et al., 2018; Lau et al., 2020; H Li and Durbin, 2011; Mattle-Greminger et al., 2018; Palkopoulou, Mallick, et al., 2015; Yew et al., 2018). From these analyses, evidence for migration events have been uncovered (Browning et al., 2018; H Li and Durbin, 2011), as have genomic consequences of human activities on other species (Choo et al., 2016). Linking demographic history to climate and environmental data greatly supports the field of conservation genetics (Ekblom et al., 2018; Hendricks et al., 2018; Oh et al., 2019). Indeed, using such approaches can help ecologists in detecting effective population size decrease (Williams et al., 2020), and thus serve as a guide in maintaining or avoiding the erosion of genetic diversity in endangered populations, and potentially predicting the consequences of climate change on genetic diversity (S Li et al., 2014). In addition, studying the demographic histories of different species in relation to one another can unveil latent biological or environmental evolutionary forces (Hecht et al., 2018), unveiling links and changes within entire ecosystems. With the increased accuracy of current methods, the availability of very large and diverse data sets and the development of new theoretical frameworks, the demographic history has become an information that is essential in the field of evolution (Cao et al., 2011; Prado-Martinez et al., 2013). However, obtaining unbiased estimations/interpretations of the demographic history remain challenging (Beichman et al., 2017; Chikhi et al., 2018).

The most sophisticated methods to infer demographic history make use of whole genome polymorphism data. Among the state-of-the-art methods, are those based on the theory of the Sequentially Markovian Coalescent (SMC) developed by McVean and Cardin (2005) after the work of Wiuf and Hein (1999), corrected by Marjoram and Wall (2006) and first applied to whole genome sequences by H Li and Durbin (2011), who introduced the now well-known, Pairwise Sequentially Markovian Coalescent (PSMC) method. PSMC allows demographic inference of populations with unprecedented accuracy, while requiring only one sequenced diploid individual. This method uses the distribution of SNPs along the genome between the two sequences to account for and infer recombination and demographic history of a given population, assuming neutrality and panmixia. Although PSMC was a breakthrough in demographic inference, it has limited power in inferring more recent events. In order to address this issue, PSMC has been extended to account for multiple sequences (*i.e*. more than two) into the method known as the Multiple Sequentially Markovian Coalescent (MSMC) (Schiffels and Durbin, 2014). By using more sequences, MSMC better infers recent events and also provides the possibility of inferring population splits using the cross-coalescent rate. MSMC, unlike PSMC, is not based on SMC theory (McVean and Cardin, 2005) but on SMC’ theory (Marjoram and Wall, 2006), therefore MSMC applied to only two sequences has been defined as PSMC’. Methods developed after MSMC followed suit, with MSMC2 (Malaspinas et al., 2016) extending PSMC by incorporating pairwise analysis, increasing efficiency and the number of sequences that can be inputted (up to a hundred), resulting in more accurate results. SMC++ (Terhorst, Kamm, et al., 2017) brings the SMC theory to another level by allowing the use of hundreds of unphased sequences (MSMC requires phased input data) and breaking the piece-wise constant population size hypothesis, while accounting for the sample frequency spectrum (SFS). Because SMC++ incorporates the SFS in the estimation of demographic history, it increases accuracy in recent time (Terhorst, Kamm, et al., 2017). SMC++ is currently the state of the art SMC-based method for big data sets (>20 sequences), but seems to be outperformed by PSMC when using smaller data sets (Patton et al., 2019). In a similar vein, the Ascertained Sequentially Markovian Coalescent (ASMC) (Palamara et al., 2018) extends the SMC theory to estimate coalescence times at the locus scale from ascertained SNP array data, something that was made possible by the theory presented by Hobolth and Jensen (2014).

More recently, a second generation of SMC-based methods have been developed. New features have been added to the initial SMC theory, extending its application beyond simply inferring past demography (Barroso et al., 2019; Sellinger et al., 2020; K Wang et al., 2020). The development of C-PSMC (Hecht et al., 2018) allows the interpretation of estimated demographic history in the light of coevolution between species, making the first link between demographic history estimated by PSMC and evolutionary forces (although biological interpretation remains limited). iSMC (Barroso et al., 2019) extends the PSMC theory to account and infer the variation of the recombination rate along sequences, unlocking recombination map estimations. An impressive advancement is the development of MSMC-IM, which to some extent solves the population structure problem, allowing the accurate and simultaneous inference of the demographic history and population admixture (K Wang et al., 2020). eSMC (Sellinger et al., 2020) incorporates common biological traits (such as self-fertilization and dormancy) and demonstrated the strong effect life-history traits can have on demographic history estimations. Results which could not be explained under the initial SMC hypotheses can now be explained by the potential presence of measurable phenomena.

New methods have been developed since PSMC, that have been either strongly inspired by the SMC (Sheehan, Harris, et al., 2013; Steinrucken et al., 2019) or that are completely dissociated from it (Beeravolu et al., 2018; Johndrow and Palacios, 2019; Kardos et al., 2017; Lynch, Haubold, et al., 2020; Rodriguez et al., 2018; Smith and Flaxman, 2020; Speidel et al., 2019; Waltoft and Hobolth, 2018). Though there are alternative approaches, methods based on the SMC are still considered state of the art, and remain widely used (Beichman et al., 2017; Mather et al., 2020; Spence et al., 2018), notably in human evolution studies (Patton et al., 2019; Spence et al., 2018). However, each described method has its specificity, being based on different hypothesis in order to solve a particular problem or shortcomings of existing methodology. Although all these methods allow a new and different interpretation of genomic data, none of these methods guarantees unbiased inference, and their limitations have underlined how crucial and challenging demographic inference is, highlighting the complementarity and usefulness of applying several inference methods on a given dataset.

SMC-based methods display very good fits when using simulated data, especially when using simple single population models based on typical human data parameters (Schiffels and Durbin, 2014; Sellinger et al., 2020; Terhorst, Kamm, et al., 2017; K Wang et al., 2020). However, the SMC makes a large number of hypotheses (H Li and Durbin, 2011; Schiffels and Durbin, 2014) that are often violated in data obtained from natural populations. When inputting data from natural populations, extracting information or correctly interpreting the results can become troublesome (Beichman et al., 2017; Chikhi et al., 2018; Terhorst and Song, 2015) and several studies address the consequences of hypothesis violation (Chikhi et al., 2018; Hawks, 2017; Mazet et al., 2016; Rodriguez et al., 2018; Schrider et al., 2016). They bring to light how strongly population structure or introgression influence demographic history estimation if not correctly accounted for (Chikhi et al., 2018; Hawks, 2017). Furthermore, some SMC-based methods require phased data (such as MSMC (Schiffels and Durbin, 2014) and MSMC-IM (K Wang et al., 2020)), and phasing errors can lead to a strong overestimation of population size in recent time (Terhorst, Kamm, et al., 2017). The effect of sequencing coverage has also been tested in Nadachowska-Brzyska et al. (2016), showing the importance of high coverage in order to obtain trustworthy results, and yet, SMC methods seem robust to genome quality (Patton et al., 2019). Selection, if not accounted for, can result in a bottleneck signature (Schrider et al., 2016), and there is currently no solution to this issue within the SMC theory, though it could be addressed using different theoretical frameworks that are being developed (Nakagome et al., 2019; Sheehan and Song, 2016). More problematic, is the ratio of effective recombination over effective mutation rates 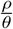 which, if it is greater than one, biases estimations (Barroso et al., 2019; Sellinger et al., 2020; Terhorst, Kamm, et al., 2017). It is also important to keep in mind that there can be deviations between 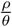 and the ratio of recombination rate over mutation rate measured experimentally 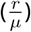 as the former can be greatly influenced by life-history, such as in organisms displaying self-fertilization, parthenogenesis or dormancy, and this can lead to issues when interpreting results (*e.g*. Sellinger et al., 2020). It is thus necessary to keep in mind that the accuracy of SMC-based methods depends on which of the many underlying hypothesis are prone to being violated by the data sets being used.

In an attempt to complement previous works, we here study the limits and convergence properties of methods based on the Sequentially Markovian Coalescent. We first define the limits of SMC-based methods (*i.e*. how well they perform theoretically), which we will call the best-case convergence. In order to do this, we use a similar approach to Gattepaille et al. (2016), Johndrow and Palacios (2019), and Palacios et al. (2015), and compare simulation results obtained with the simulated Ancestral Recombination Graph (ARG) as input to results obtained from sequences simulated under the same ARG, so as to study the convergence properties linked to data sets in the absence of hypothesis violation. We test several scenarios to check whether there are instances, where even without violating the underlying hypotheses of the methodology, the demographic scenarios cannot be retrieved because of theoretical limits (and not issues linked with data). We also study the effect of the optimization function (or composite likelihood) and the time window of the analysis on the estimations of different variables. Lastly, we test the effect of commonly violated hypotheses, such as the effect of the variation of recombination and mutation rates along the sequence and between scaffolds, errors in SNP calls and the presence of transposable elements and link abnormal results to specific hypothesis violations. Through this work, our aim is to provide guidelines concerning the interpretation of results when applying this methodology on data sets that may violate the underlying hypotheses of the SMC framework.

## Methods

In this study we use four different SMC-based methods: MSMC, MSMC2, SMC++ and eSMC. All methods are Hidden Markov Models and use whole genome sequence polymorphism data. The hidden states of these methods are the coalescence times (or genealogies) of the sample. In order to have a finite number of hidden states, they are grouped into *x* bins (*x* being the number of hidden states). The reasons for our model choices are as follows: *i*) MSMC, unlike any other method, focuses on the first coalescence event of a sample of size *n*, and thus exhibits different convergence properties (Schiffels and Durbin, 2014), *ii*) MSMC2 computes coalescence times of all pairwise analysis from a sample of size *n*, and can deal with a large range of data sets (Speidel et al., 2019), *iii*) SMC++ (Terhorst, Kamm, et al., 2017) is the most advanced and efficient SMC method and lastly, *iv*) eSMC (Sellinger et al., 2020) is a re-implementation of PSMC’ (similar to MSMC2), which will contribute to highlighting the importance of algorithmic translations as it is very flexible in its use and outputs intermediate results necessary for this study.

## SMC methods

### PSMC’, MSMC2 and eSMC

PSMC’ and methods that stem from it (such as MSMC2 (Malaspinas et al., 2016) and eSMC (Sellinger et al., 2020)) focus on the coalescence events between only two individuals (or sequences in practice), and, as a result, do not require phased data. The algorithm goes along the sequence and estimates the coalescence time at each position. In order to do this, it checks whether the two sequences are similar or different at each position. The presence or absence of a segregating site along the sequence (determined by the population mutation rate *θ*) is used to infer the hidden state (*i.e*. coalescence time). However, the hidden state is only allowed to change in the event of a recombination, which leads to a break in the current genealogy. Thus, the population recombination rate *ρ* constrains the inferred changes of hidden states along the sequence (for a detailed description of the algorithm see Schiffels and Durbin (2014), Sellinger et al. (2020), and K Wang et al. (2020)).

### MSMC

MSMC is mathematically and conceptually very similar to the PSMC’ method. Unlike other SMC methods, it simultaneously analyses multiple sequences and because of this, MSMC requires the data to be phased. In combination with a second HMM, to estimate the external branch length of the genealogy, it can follow the distribution of the first coalescence event in the sample along the sequences. However, due to computational load, MSMC cannot analyze more than 10 sequences simultaneously (for a detailed description see Schiffels and Durbin (2014)).

### SMC++

SMC++ is slightly more complex than MSMC or PSMC. Though it is conceptually very similar to PSMC’, mathematically it is quite different. SMC++ has a different emission matrix compared to previous methods because it calculates the sample frequency spectrum of sample size *n* + 2, conditioned on the coalescence time of two “distinguished” haploids and *n* “undistinguished” haploids. In addition SMC++ offers features such as a cubic spline to estimate demographic history (*i.e*. not a piece-wise constant population size). The SMC++ algorithm is fully described in Terhorst, Kamm, et al. (2017).

### Best-case convergence

Using sequence simulators such as msprime (Kelleher, Etheridge, et al., 2016) or scrm (Staab et al., 2015), one can simulate the Ancestral Recombination Graph (ARG) of a sample. Usually the ARG is given through a sequence of genealogies (*e.g*. a sequence of trees in Newick format). From this ARG, one can find what state of the HMM the sample is in at each position. Hence, one can build the series of states along the genomes, and build the transition matrix. The transition matrix, is a square matrix of dimension *x* (where *x* is the number of hidden states) counting all the possible pairwise transitions between the *x* states (including from a given state to itself). Using the transition matrix built directly from the exact ARG, one can estimate parameters using eSMC or MSMC as if they could correctly infer the hidden states. Hence estimations using the exact transition matrix represents the upper bound of performance for these methods. We choose to call this upper bound the best-case convergence (since it can never be reached in practice). For this study’s purpose, a second version of the R package eSMC (Sellinger et al., 2020) was developed. This package enables the building of the transition matrix (for eSMC or MSMC), and can then use it to infer the demographic history. The package is mathematically identical to the previous version, but includes extra functions, features and new outputs necessary for this study. The package and its description can be found at https://github.com/TPPSellinger/eSMC2.

### Baum-Welch algorithm

SMC-based methods can use different optimization functions to infer the demographic parameters (*i.e*. likelihood or composite likelihood). The four studied methods use the Baum-Welch algorithm to maximize the likelihood. MSMC2 and SMC++ implement the original Baum-Welch algorithm (which we call the complete Baum-Welch algorithm), whereas eSMC and MSMC compute the expected composite likelihood *Q*(*θ*|*θ*^*t*^) based only on the transition matrix (which we call the incomplete Baum-Welch algorithm). The use of the complete Baum-Welch algorithm or the incomplete one can be specified in the eSMC package. The composite likelihood for SMC++ and MSMC2 is given by equations 1 and the composite likelihood for eSMC and MSMC by equation 2:

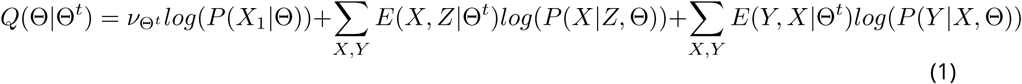

and:

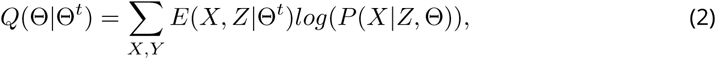

with:

- *ν*Θ: The equilibrium probability conditional to the set of parameters Θ.
- *P* (*X*_1_|Θ): The probability of the first hidden state conditional to the set of parameters Θ.
- *E*(*X, Z*|Θ^*t*^): The expected number of transitions of X from Z conditional to the observation and set of parameters Θ^*t*^.
- *P* (*X*|*Z*, Θ): The transition probability from state Z to state X, conditional to the set of parameters Θ.
- *E*(*Y, X*|Θ^*t*^) The expected number of observations of type Y that occurred during state X conditional to observation and set of parameters Θ^*t*^.
- *P* (*Y* |*X*, Θ): The emission probability conditional to the set of parameters Θ.

### Time window

Each tested SMC-based method has its own specific time window for which estimations are made. Note that hidden states are defined as discretized intervals, as a consequences of which the boundaries, length and number of states of the time window do implicitly affect inferences. For example, the original PSMC has a time window wider than PSMC’, so that estimations cannot be compared one to one. To measure the effect of choosing different time window parameters, we analyze the same data with four different settings. The first time window is the one used for PSMC’ defined in Schiffels and Durbin (2014). The second time window is that of MSMC2 (K Wang et al., 2020) (similar to the one of the original PSMC (H Li and Durbin, 2011)), which we call “big” since it goes further in the past and in more recent time than that of PSMC’. We then define a time window equivalent to the first one (i.e. PSMC’) shifted by a factor five in the past (first time window, *i.e*. hidden states, multiplied by five). The last one is a time window equivalent to the first one (i.e. PSMC’) shifted by a factor five in recent time (i.e. first time window divided by five).

### Regularization penalty

To avoid inferring unrealistic demographic histories with variations of population size that are too strong and/or too rapid, SMC++ introduced a regularization penalty (https://github.com/popgenmethods/smcpp). This parameter penalizes population size variation. In SMC++, the lower value of the penalty the more the estimated size history is a line (*i.e*. constant population size). Regularization penalty was also implemented in eSMC. Setting the regularization penalty parameter to 0 is equivalent to no penalization, and the higher the parameter value, the more population size variations are penalized (https://github.com/TPPSellinger/eSMC2 for more details). We tested the effect of regularization on inferences with both methods using simulated sequence data. The sequence data was simulated under sawtooth demographic histories with different amplitudes of population size variation. All the command lines to analyze the data generated can be found in the Appendix 2.

### Simulated sequence data

Throughout this paper we simulate different demographic scenarios using either the coalescence simulation program scrm (Staab et al., 2015) or msprime (Kelleher, Etheridge, et al., 2016). We use scrm for the best-case convergence as it can output the genealogies in a Newick format (which we use as input). We use scrm, which outputs simulated sequences in the ms format, to simulate data for eSMC, MSMC, MSMC2. We use msprime to simulate data for SMC++ since msprime is more efficient than scrm for big sample sizes (Kelleher, Etheridge, et al., 2016) and can directly output .vcf files (which is the input format of SMC++).

### Absence of hypothesis violation

We simulate five different demographic scenarios: saw-tooth (successions of population size exponential expansion and decrease), bottleneck, exponential expansion, exponential decrease and constant population size. Each of the scenarios with varying population size is tested under four amplitude parameters (*i.e*. by how many fold the population size varies: 2, 5, 10, 50). We infer the best-case convergence under four different sequence lengths (10^7^, 10^8^, 10^9^ and 10^10^ bp) and choose the per site mutation and recombination rates recommended for humans in MSMC’s manual, respectively 1.25*×*10^*−*8^ and 1*×*10^*−*8^ (https://github.com/stschiff/msmc/blob/master/guide.md). When analyzing simulated sequence data, we simulate sequences of 100 Mb: two sequences for eSMC and MSMC2, four sequences for MSMC and twenty sequences for SMC++.

### Calculation of the mean square error (MSE)

To measure the accuracy of inferences we calculate the Mean Square Error (MSE). We first divide the time window (in log10 scale) of each analysis into ten thousand points. We then calculate the MSE by comparing the actual population size and the one estimated by the method at each of the ten thousand points. We thus have the following formula:

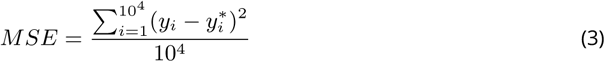

Where:

- *y*_*i*_ is the population size at the time point *i*.
- 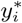 is the estimated population size at the time point *i*.

All the command lines to simulate data can be found in the Appendix 1.

### Presence of hypothesis violation

#### SNP calling

In practice, SNP calling from next generation sequencing can yield different numbers and frequencies of SNPs depending on the chosen parameters for the different steps of analysis (read trimming, quality check, read mapping, and SNP calling) as well as the quality of the reference genome, data coverage and depth of sequencing, species ploidy (Pfeifer, 2017). Therefore, based on raw sequence data, the stringency of filters can lead to excluding SNPs (false negatives) or including spurious ones (false positives). When dealing with complex genomes or ancient DNA (Chang and Shapiro, 2016; Slatkin, 2016), SNPs can be simultaneously missed and added. We thus simulate four sequences of 100 Mb under a “saw-tooth” scenario and then a certain percentage (5,10 and 25 %) of SNPs is randomly added to and/or deleted from the simulated sequences. We then analyze the variation and bias in SNP calling on the accuracy of demographic parameter estimations. As an additional analysis we test the effect of ascertainment bias on inferences (a prominent issue in microarray SNP studies) by simulating 100 sequences with msprime where only SNPs above a certain (Minor Allele Frequency) MAF threshold (1%,5% and 10%) are kept, then run SMC methods on a subset of the obtained data.

### Changes in mutation and recombination rates along the sequence

Because the recombination rate and the mutation rate can change along the sequence (Barroso et al., 2019), and chromosomes are not always fully assembled in the reference genome (which consists of possibly many scaffolds), we simulate short sequences where the recombination and/or mutation rate randomly change between the different scaffolds around an average value of 1.25 *×* 10*−*8 per generation per base pair (between 2.5*×* 10*−*9and 6.25 *×* 10*−*8). We simulate 20 scaffolds of size 2 Mb, as this seems representative of the best available assembly for non-model organisms (Lynch, Gutenkunst, et al., 2017; Stam et al., 2019). We then analyze the simulated sequences to study the effect of assuming scaffolds share the same mutation and recombination rates. In addition, we simulate sequences of 40 Mb (assuming genomes are fully assembled) where the recombination rate along the sequence randomly changes every 2 Mbp (up to five-fold) around an average value of 1.25 *×* 10*−*8 (the mutation rate being fixed at 1.25 *×* 10*−*8 per generation per bp) to study the effect of the assumption of a constant recombination rate along the sequence.

### Transposable elements (TEs)

Genomes can contain transposable elements whose dynamics violate the classic infinite site mutational model for SNPs, and thus potentially affect the estimation of different parameters. Although methods have been developed to detect (Nelson et al., 2017) and simulate them (Kofler, 2018), understanding how their presence/absence influences demographic inferences remains unclear. TEs are usually masked when detected in the reference genome and thus not taken into account in the mapped individuals due to the redundancy of read mapping for TEs. Due to their repetitive nature, it can be difficult to correctly detect and assemble them if using short reads, as well as to assess the presence/absence polymorphism of individuals of a population (Ewing, 2015). In addition, the quality and completeness of the reference genome (*e.g*. using the reference genome of a sister species as the reference genome) can strongly affect the accuracy of detecting, assembling and masking TEs (Platt et al., 2016). To best capture and mimic the effect of TEs unaccounted for in the data, we altered four simulated sequences of length 20 Mb in four different ways. The first way to simulate the effect of unmapped and unaccounted TEs is to assume they exhibit presence/absence polymorphism, hence creating gaps in the sequence. For each individual, we remove small pieces of sequence of different length (1kb, 10 kb or 100kb), so that up to a certain percentage (5,10,25,50%) of the original simulated sequence is removed, so as to shorten and fragment the whole sequence to be analyzed. The second way, is to consider unmasked TEs, done by randomly selecting small pieces of the original simulated sequence (1kb, 10 kb or 100kb), making up to a certain percentage of it (5,10,25,50%), and removing all the SNPs found in those regions (*i.e*. removing mutations from TEs). The removed SNPs are hence structured in many small regions along the genome. Thirdly, we test the consequences of simultaneously having both removed and unmasked TEs in the data set. Lastly, to measure the importance of detecting and masking TEs, we assume all TEs to be present and masked when building the multihetsep file (*i.e*. considering TEs as missing data).

## Results

### Best-case convergence

Results of the best-case convergence of eSMC under the saw-tooth demographic history are displayed in Figure 1. Increasing the sequence length increases accuracy and reduces variability, leading to better convergence and reducing the mean square error (see Figures 1a-c and Supplementary Table 1). However, when the amplitude of population size variation is too great (here for 50 fold), the demographic history cannot be retrieved, even when using very large data sets (see Figure 1d). Similar results are obtained for the three other demographic scenarios (bottleneck, expansion and decrease, respectively displayed in Supplementary Figures 1, 2 and 3). The bottleneck scenario seems especially difficult to infer, requiring large amounts of data, and the stronger the bottleneck, the harder it is to detect it, even with sequence lengths equivalent to 10^10^bp. In Supplementary Figure 4, we show that even when changing the number of hidden states (*i.e*. number of inferred parameters), some scenarios with very strong variation of population size remain badly inferred.

**Table 1.** Average estimated values for the recombination over mutation ratio 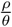by eSMC over ten repetitions for different sizes of the time window. The coefficient of variation is indicated in brackets. Four sequences of 50 Mb were simulated with a recombination rate set to 1 *×* 10^*−*8^ per generation per bp and a mutation rate to 1.25 *×* 10^*−*8^ per generation per bp.

| Optimization function | Scenario | real $\frac{\rho}{\theta}$ | normal window $\frac{\rho}{\theta}^*$ | Big Window $\frac{\rho}{\theta}^*$ | Old window $\frac{\rho}{\theta}^*$ | Recent window $\frac{\rho}{\theta}^*$ |
| --- | --- | --- | --- | --- | --- | --- |
| Incomplete Baum-Welch | Saw-tooth | 0.8 | 0.79 (0.036) | 0.72 (0.039) | 0.72 (0.042) | 0.94 (0.005) |
| Complete Baum-Welch | Saw-tooth | 0.8 | .79 (0.044) | 0.72 (0.039) | 0.72 (0.042) | 1.56 (0.087) |
| Incomplete Baum-Welch | Constant | 0.8 | 0.86 (0.019) | 0.85 (0.020) | 0.84 (0.019) | 0.98 (0.002) |
| Complete Baum-Welch | Constant | 0.8 | 0.86 (0.019) | 0.85 (0.020) | 0.84 (0.019) | 1.06 (0.02) |

**Figure 1.**
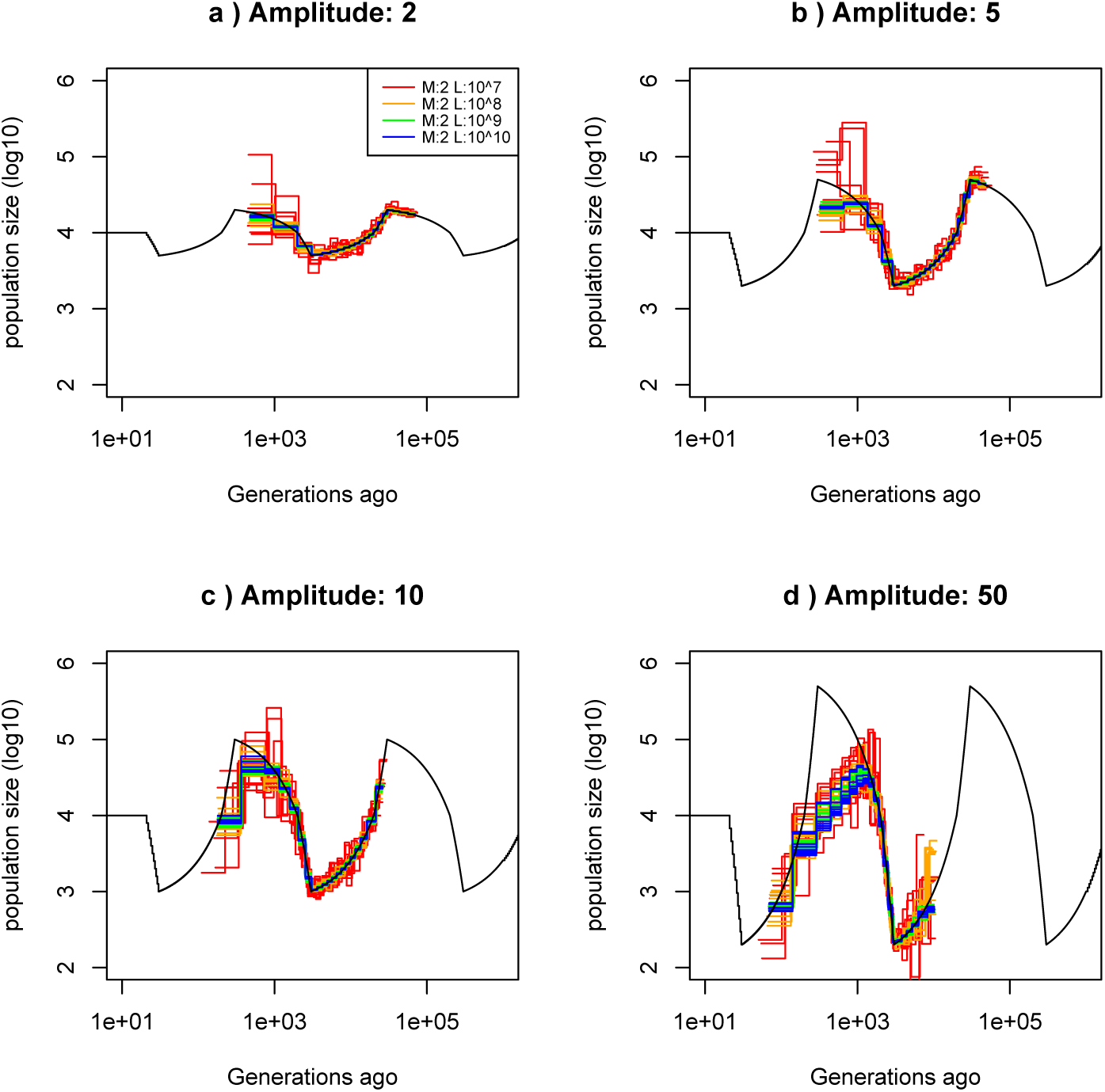
Best-case convergence of eSMC. Estimated demographic history using simulated genealogy over sequences of 10,100,1000,10000 Mb (respectively in red,orange, green and blue) under a saw-tooth scenario (original scenario in black) with 10 replicates for different amplitudes of size change: a) 2-fold, b) 5-fold, c) 10-fold, and d) 50-fold. The recombination rate is set to 1 *×* 10^*−*8^ per generation per bp and the mutation rate to 1.25 *×* 10^*−*8^ per generation per bp.

**Figure 2.**
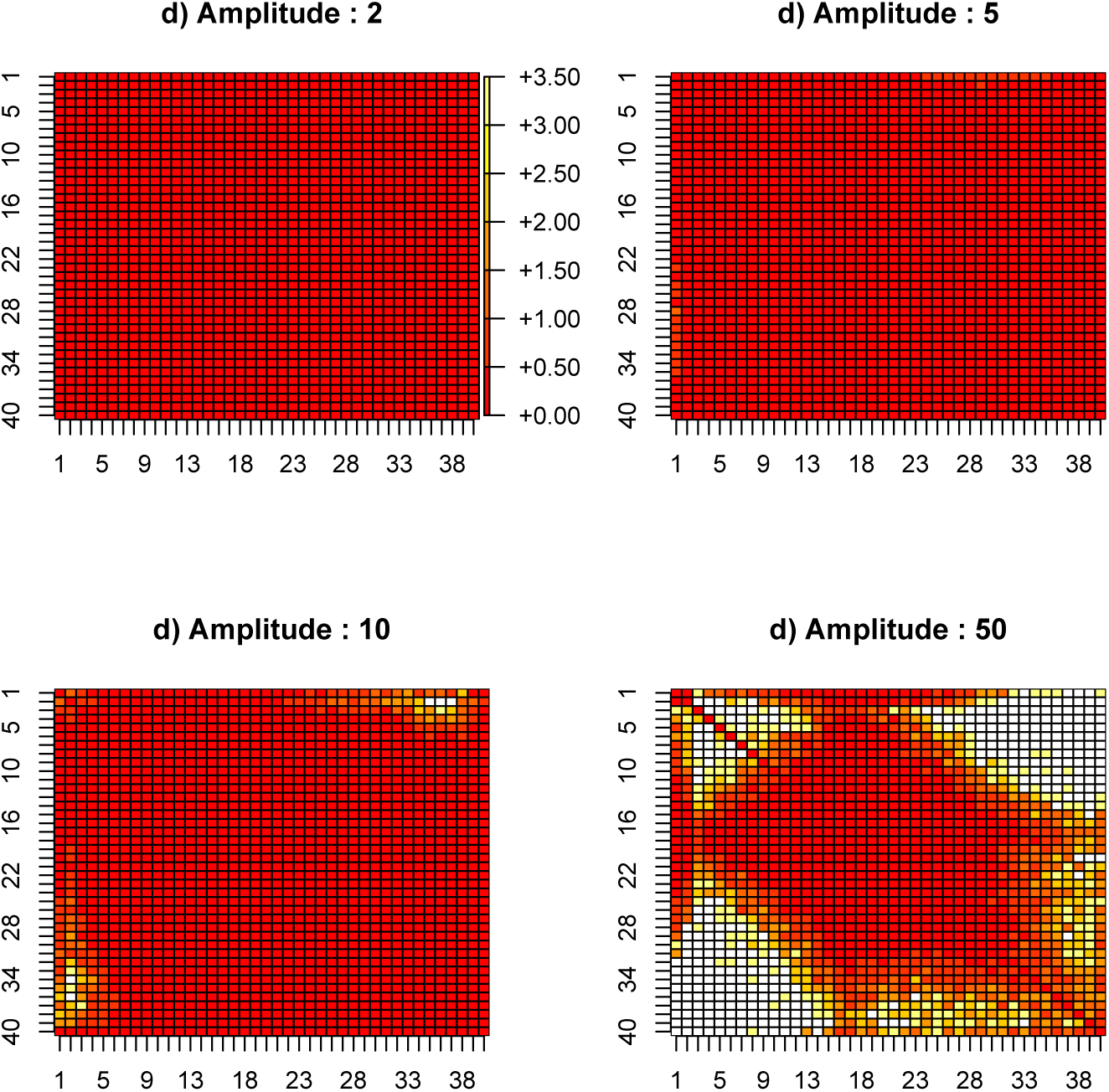
Estimated transition matrix in sharp saw-tooth scenario. Estimated coefficient of variation of the transition matrix using simulated genealogy over sequences of 10000 Mb under a saw-tooth scenario of amplitude 2, 5,10 and 50 (respectively in a, b, c and d) each with 10 replicates. Recombination and mutation rates are as in Figure 1. White squares indicate absence of observed transitions (*i.e. no data*).

**Figure 3.**
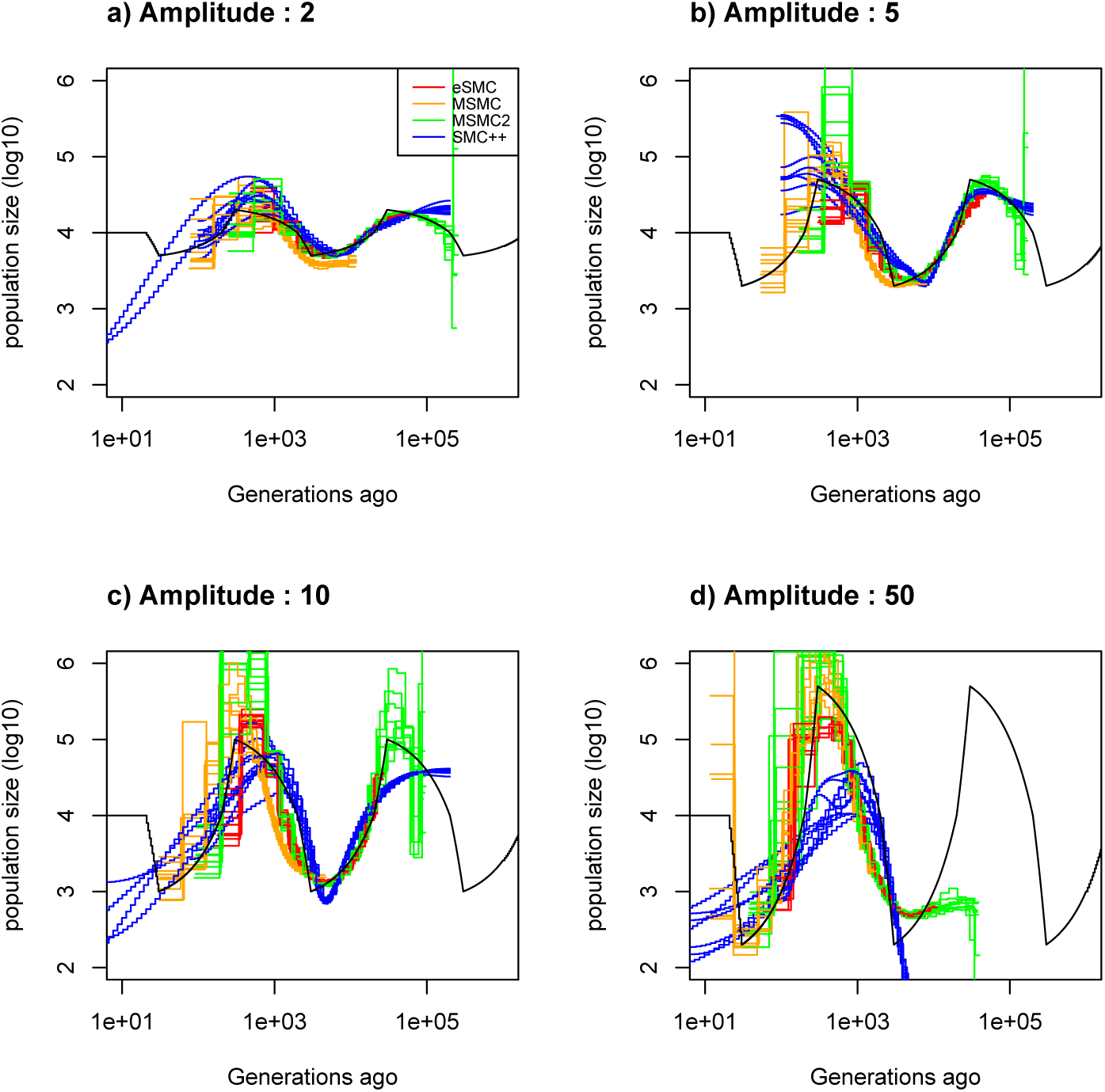
Estimated demography using simulated sequences as input. Estimated demographic history (black) under a saw-tooth scenario with 10 replicates using simulated sequences for different amplitudes of population size change: a) 2, b) 5, c) 10 and d) 50. Two sequences of 100 Mb for eSMC and MSMC2 (respectively in red and green), four sequences of 100 Mb for MSMC (orange) and 20 sequences of 100 Mb for SMC++ (blue) were simulated. Recombination and mutation rates are respectively set to 1 *×* 10^*−*^8 and 1.25 *×* 10^*−*^8.

**Figure 4.**
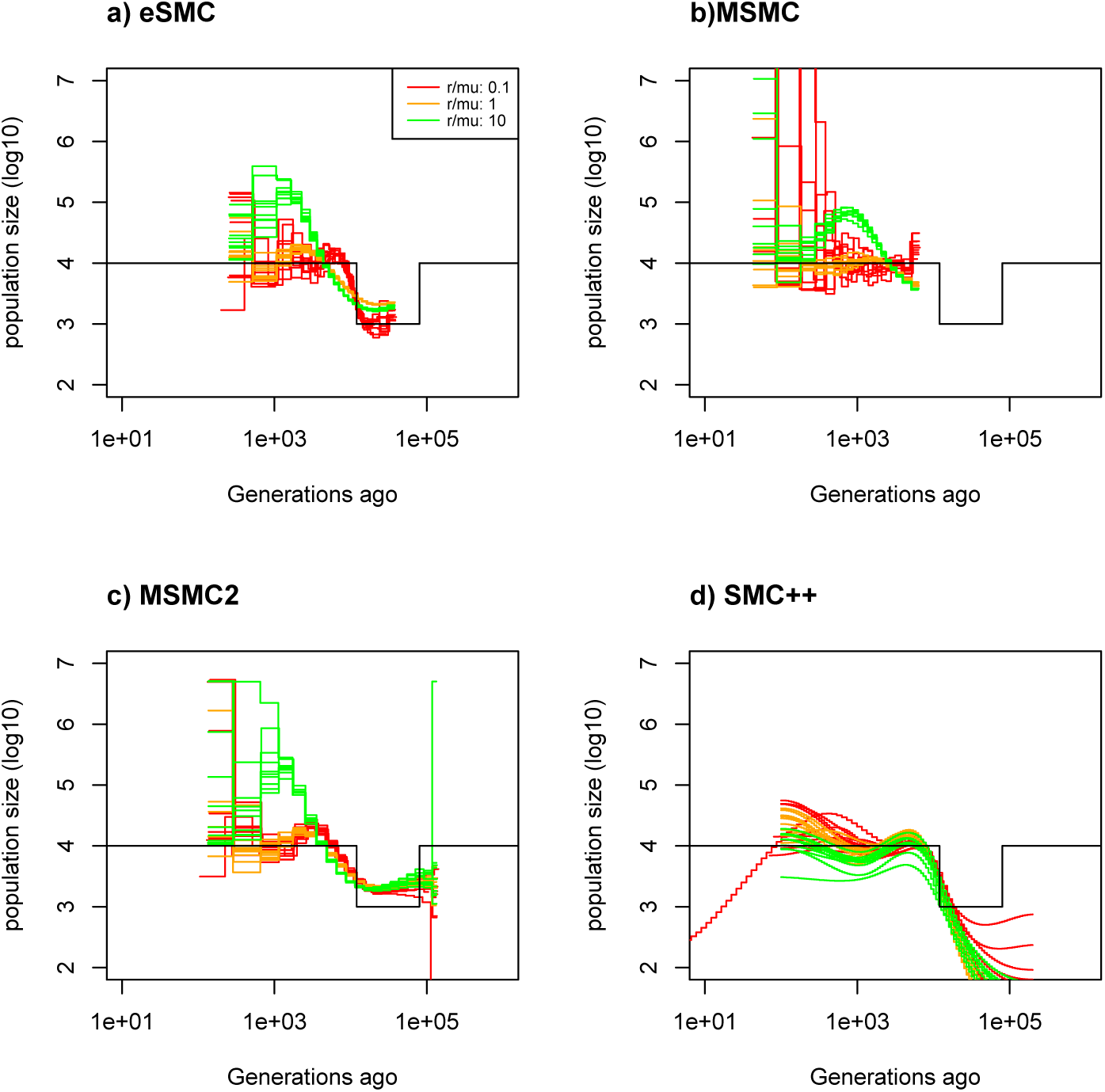
Effect of 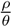 on inference of demographic history. Estimated demographic history under a bottleneck scenario with 10 replicates using simulated sequences. We simulate two sequences of 100 Mb for eSMC and MSMC2 (respectively in a and b), four sequences of 100 Mb for MSMC (c) and twenty sequences of 100 Mb for SMC++ (d). The mutation rate is set to 1.25 *×* 10^*−*8^ per generation per bp and the recombination rates are 1.25 *×* 10^*−*9^,1.25 *×* 10^*−*8^ and 1.25 *×* 10^*−*7^ per generation per bp, giving 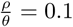, 1 and 2 and the inferred demographies are in red, orange and green respectively. The demographic history is simulated under a bottleneck scenario of amplitude 10 and is represented in black.

In Supplementary Figures 5, 6, 7 and 8, we show the best-case convergence of MSMC with four genome sequences and generally find that these analyses present a higher variance than eSMC. However, MSMC shows better fits in recent times and is better able to retrieve population size variation than eSMC (see Supplementary Figure 5d). Scenarios with strong variation of population size (*i.e*. with large amplitudes) still pose a problem (see Supplementary Figure 9), and no matter the number of estimated parameters, such scenarios cannot be correctly inferred using MSMC.

**Figure 5.**
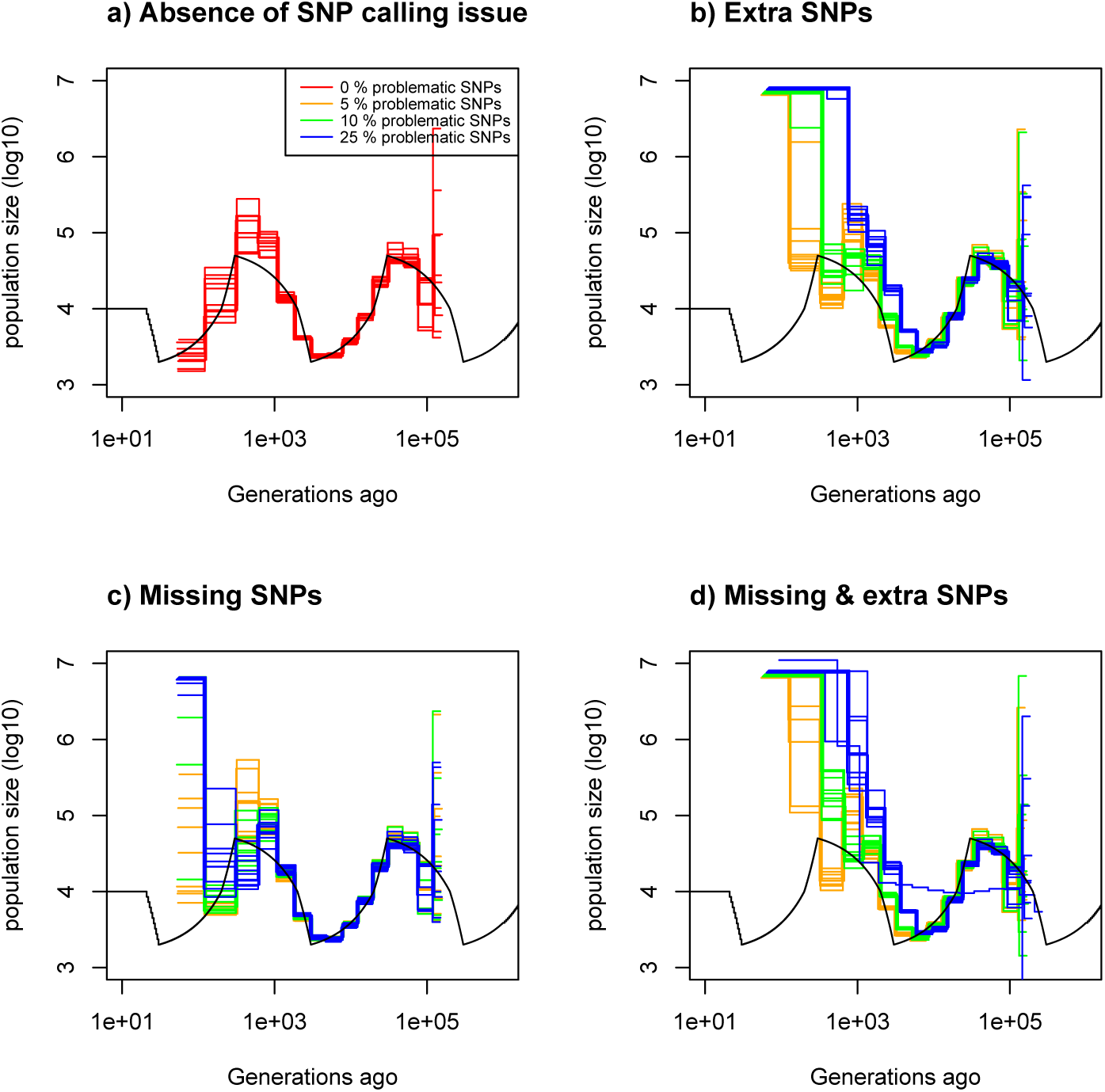
Consequences of SNP calling errors. Estimated demographic history using MSMC2 under a saw-tooth scenario with 10 replicates using four simulated sequences of 100 Mb. Recombination and mutation rates are as in Figure 1 and the simulated demographic history is represented in black. Demographic history simulated with ibsence of SNP calling issue (red). b) Demographic history simulated with 5% (orange),10% (green) and 25% (blue) missing SNPs. c) Demographic history simulated with 5% (orange), 10% (green) and 25% (blue) additional SNPs. d) Demographic history simulated with 5% (orange),10% (green) and 25% (blue) of additional and missing SNPs.

**Figure 6.**
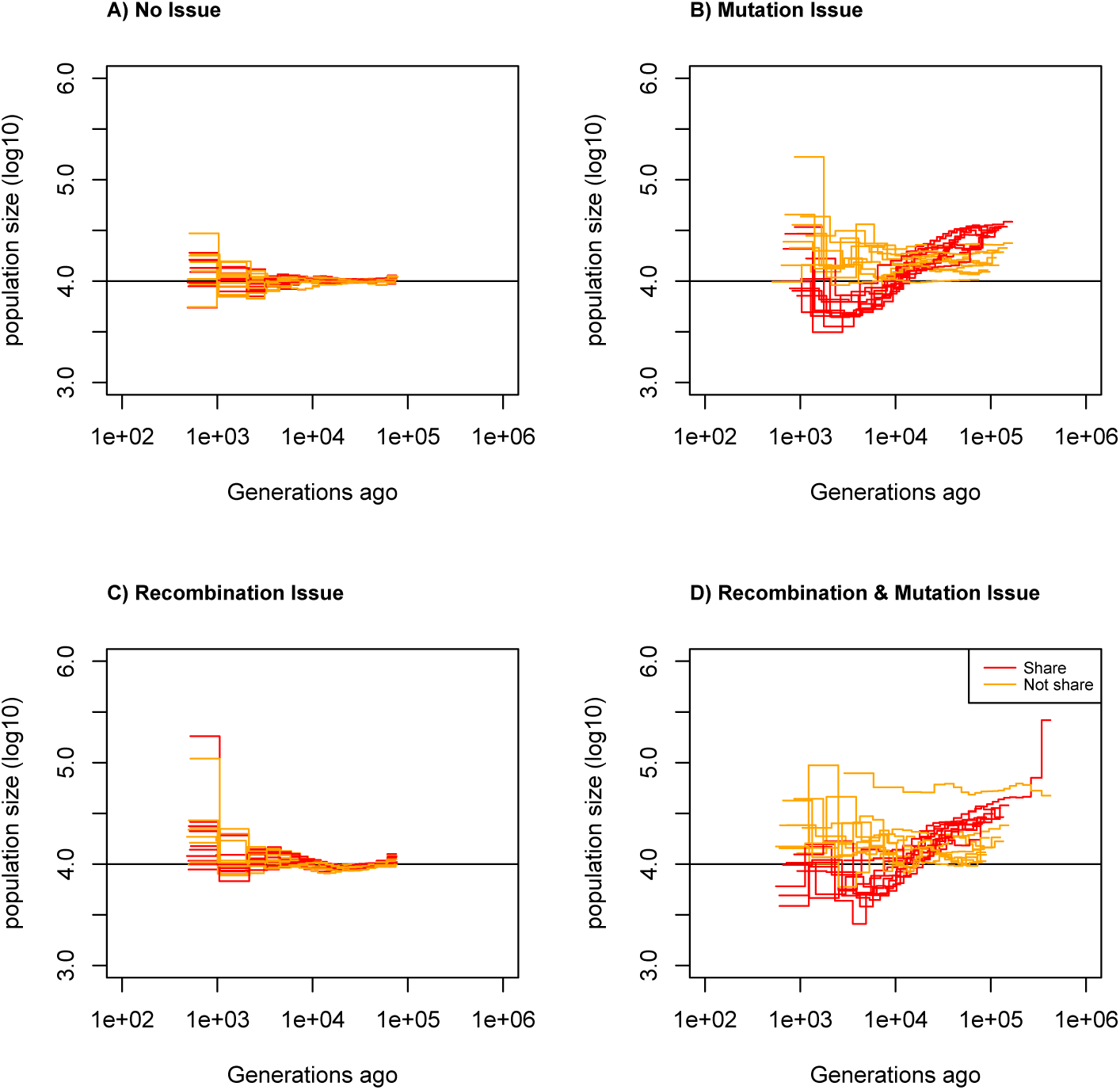
Estimating demographic history using scaffolds sharing or differing in mutation and recombination rates. Estimated demographic history using eSMC under a saw-tooth scenario with 10 replicates using twenty simulated scaffolds of two sequences of 2 Mb assuming scaffolds share (red) or do not share recombination and mutation rates (orange). The simulated demographic history is represented in black. a) Scaffolds share the same parameters, recombination and mutation rates are set at 1.25 *×* 10^*−*8^, b) Each scaffold is randomly assigned a recombination rate between 2.5 *×* 10^*−*9^ and 6.25 *×* 10^*−*8^ and the mutation rate is 1.25 *×* 10^*−*8^, c) Each scaffold is randomly assigned a mutation rate between 2.5 *×* 10^*−*9^ and 6.25 *×* 10^*−*8^ and the recombination rate is 1.25 *×* 10^*−*8^ and d) Each scaffold is assigned a random mutation and an independently random recombination rate, both being between 2.5 *×* 10^*−*9^ and 6.25 *×* 10^*−*8^.

**Figure 7.**
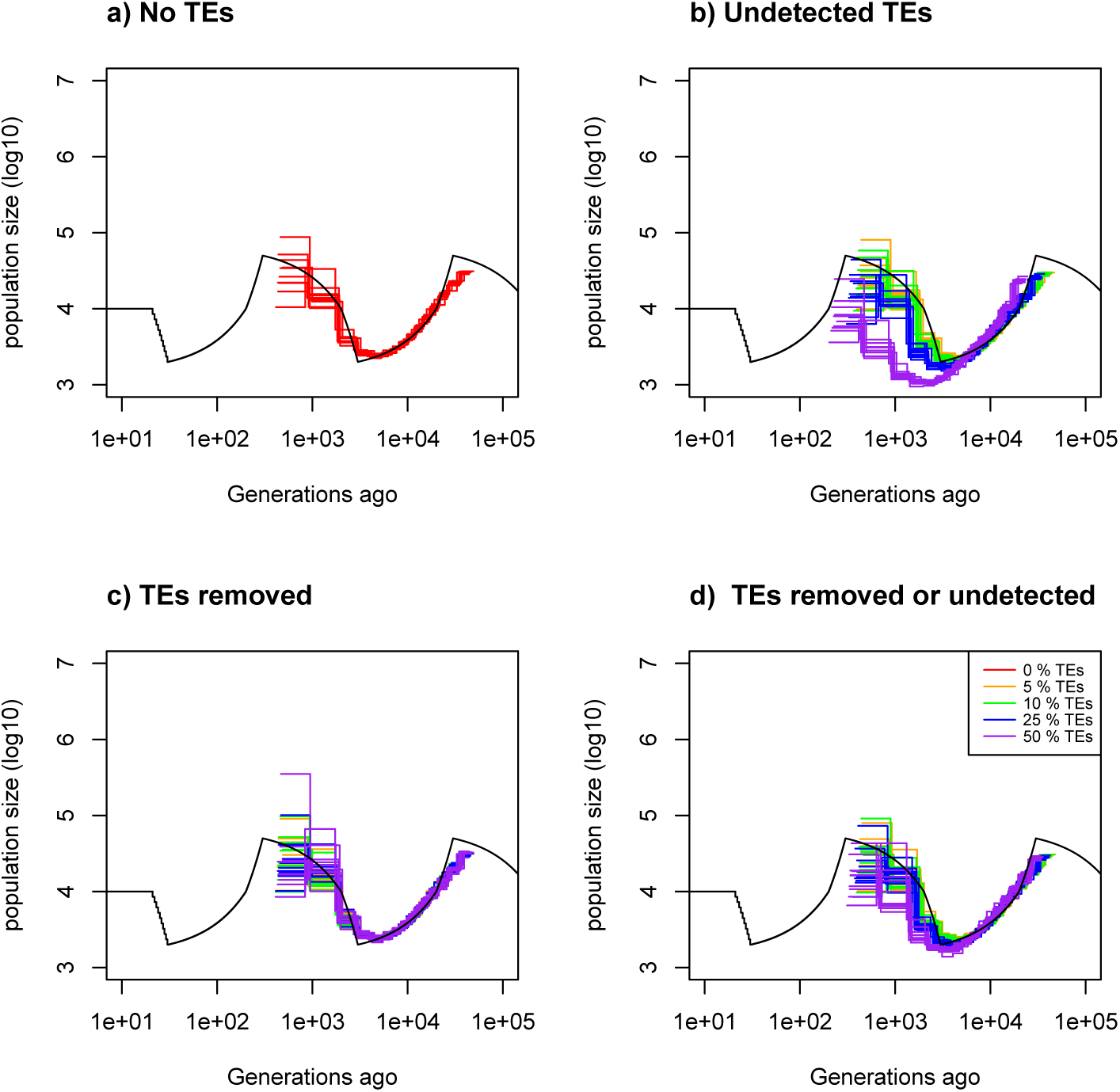
Consequences of masking or removing transposable elements (TEs) from data sets. Estimated demographic history by eSMC under a saw-tooth scenario with 10 replicates using four simulated sequences of 20 Mb. The recombination and mutation rates are as in Figure 1 and the simulated demographic history is represented in black. Here the TEs are of length 1kbp. a) Demographic history simulated with no TEs. b) Demographic history simulated where TEts are removed. c) Demographic history simulated where TEs are masked. d) Demographic history simulated where half of the TEs are removed and SNPs on the other half are removed. Proportion of the genome made up by TEs is set to 0% (red), 5% (orange), 10% (green), 25 % (blue) and 50 % (purple).

To better understand these results, we examine the coefficient of variation calculated from the replicates at each entry of the transition matrix. We can see that increasing the sequence length reduces the coefficient of variation (the ratio of the standard deviation to the mean, hence indicating convergence when equal to 0, see Supplementary Figure 10). Yet increasing the amplitude of population size variation decreases the number of some hidden state transitions leading to unobserved transitions. Unobserved transitions result from the reduced probability of coalescence events in specific time intervals (*i.e*. hidden states). In these cases matrices display higher coefficients of variation and can be partially empty (Figure 2). This explains the increase of variability of the inferred scenarios, as well as the incapacity of SMC methods to correctly infer the demographic history with strong population size variation in specific time intervals independently of the amount of data available.

### Simulated sequence results

#### Scenario effect

In the previous section, we explored the theoretical performance limitations of eSMC and MSMC using trees in Newick format as input. In this section, we evaluate how these methods perform when inputting simulated sequence data using the same recombination and mutation rates. We first perform two benchmark analyses, the constant population size scenario (Supplementary Figure 11) and the sawtooth demographic scenario from Schiffels and Durbin (2014) (Supplementary Figure 12). eSMC and MSMC2 retrieve the constant population size scenario although MSMC fails to retrieve it in the far past and SMC++ in recent time (Supplementary Figure 11). All methods can retrieve the sawtooth demographic scenario although SMC++ displays high variance in recent times (Supplementary Figure 12). Secondly, we investigate the effect of amplitude of population size variation as in Figure 1. Results for the saw-tooth scenario are displayed in Figure 3, where the different models display a good fit, but are not as good as expected from the best-case convergence given the same amount of data (Figure 1 (orange line) and Supplementary Table 1 vs Figure 3 (red line) and Supplementary Table 2). As predicted by Figures 1 and 2, the case with the greatest amplitude of population size variation (Figure 1d) is the least well fitted (see Supplementary Table 2 for the MSE). All estimations display low variance and a relatively good fit in the bottleneck and expansion scenarios for small population size variation (see Supplementary Figures 13a and 14a). However, the strengths of expansions and bottlenecks are not fully retrieved in scenarios with population size variation equals to or is higher than tenfold the current population size (Supplementary Figures 13c-d,and 14c-d). To study the origin of differences between simulation results and theoretical results, we measure the difference between the transition matrix estimated by eSMC and the one built from the actual genealogy. Results show that hidden states are harder to correctly infer in scenarios with strong population size variation, explaining the high variance (see Supplementary Figure 15). We demonstrate there that for the same amount of data, the simulation, and thus by extension the real data, shows additional stochastic behaviour than the best-case convergence (Figure 1).

**Table 2.** Average estimated values for the recombination over mutation ratio 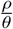 over ten repetitions. The coefficient of variation is indicated in brackets. For eSMC, MSMC and MSMC2 we have: set 1: 20 hidden states; set 2: 200 iterations; set3: 60 hidden states; set 4: 60 hidden states and 200 iterations and set 5: 20 hidden states and 200 iterations. For SMC++: set 1: 16 knots; set 2: 200 iterations; set 3: 4 knots in green; set 4: regularization penalty set to 3 and set 5: regularization-penalty set to 12.

| method | real $\frac{\rho}{\theta}$ | set 1, $\frac{\rho}{\theta}^*$ | set 2, $\frac{\rho}{\theta}^*$ | set 3, $\frac{\rho}{\theta}^*$ | set 4, $\frac{\rho}{\theta}^*$ | set 5, $\frac{\rho}{\theta}^*$ |
| --- | --- | --- | --- | --- | --- | --- |
| eSMC | 10 | 1.35 (0.026) | 1.76 (0.047) | 1.29 (0.027) | 1.74 (0.048) | 1.80 (0.041) |
| MSMC | 10 | 2.70 (0.011) | 6.58 (0.031) | 2.68 (0.011) | 6.57 (0.032) | 6.62 (0.030) |
| MSMC2 | 10 | 1.27 (0.055) | 1.65 (0.13) | 1.26 (0.060) | 1.75 (0.060) | 1.60 (0.29) |
| SMC++ | 10 | 0.56 (0.38) | 0.48 (0.38) | 1.32 (0.15) | 0.21 (0.62) | 0.98 (0.24) |

Increasing the time window in eSMC results in an increased variance of the inferences (Supplementary Figure 16). In addition, shifting the window towards more recent time leads to poor demographic estimations, but shifting it further in the past does not seem to bias it (there are however consequences on estimations of the recombination rates, see Table 1 for more details). Concerning the optimization function, we find that the complete Baum-Welch algorithm gives similar results to the incomplete one (Table 1).

Adding a regularization penalty to eSMC can drastically impact inferences (Supplementary Figure 17) and reduces performance quality. When regularization is added, eSMC fails to correctly capture the amplitude of population size variation and with extreme regularization penalty, eSMC infers a constant population size. Yet, adding regularization in SMC++ can increase performance and avoid spurious variation of population size (Supplementary Figure 18). However, strong regularization can lead to the inference of constant population size and thus poor estimations.

### Effect of the ratio of the recombination over the mutation rate

The ratio of the effective recombination over effective mutation rates 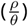 can influence the ability of SMC-based methods to retrieve the coalescence time between two points along the genome (Terhorst, Kamm, et al., 2017). Intuitively, if recombination occurs at a higher rate compared to mutation, then it renders it more difficult to detect any recombination events that may have taken place before the introduction of a new mutation, and thus bias the estimation of the coalescence time (Sellinger et al., 2020; Terhorst, Kamm, et al., 2017). Under the bottleneck scenario, we find that the lower 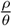 the better the fit of the inferred demography by eSMC and SMC++ in the past, but also the higher the variance of the inferences (see Figure 4). However each method displays the worse fit when 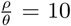 = 10 (Supplementary Table 3). SMC++ seems slightly less sensitive to 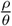 than other methods. When calculating the difference between the transition matrix estimated by eSMC and the one built from the actual genealogy (ARG), we find that, unsurprisingly, changes in hidden states are harder to detect when 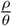 increases, leading to an overestimation of hidden states on the diagonal (see Supplementary Figures 19, 20 and 21).

**Table 3.** Average estimated values for the recombination over mutation ratio 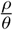over ten repetitions. The coefficient of variation is indicated in brackets. TEs are of length 1kb, 10kb or 100 kb and are completely removed and the proportion of the genome made up by TEs is 5%,10%, 25% and 50%.

| TE length | method | real $\frac{\rho}{\theta}$ | $\frac{\rho}{\theta}^*$ and 5% TEs | $\frac{\rho}{\theta}^*$ and 10% TEs | $\frac{\rho}{\theta}^*$ and 25% TEs | $\frac{\rho}{\theta}^*$ and 50% TEs |
| --- | --- | --- | --- | --- | --- | --- |
| 1 kb | eSMC | 1 | 0.95 (0.021) | 0.99 (0.022) | 1.16 (0.10) | 1.77 (0.36) |
|  | MSMC | 1 | 1.31 (0.098) | 1.35 (0.11) | 1.50 (0.088) | 1.91 (0.11) |
|  | MSMC2 | 1 | 0.87 (0.047) | 0.88 (0.049) | 1.0 (0.036) | 1.35 (0.035) |
| 10 kb | eSMC | 1 | 0.96 (0.053) | 0.98 (0.066) | 1.10 (0.18) | 1.36 (0.41) |
|  | MSMC | 1 | 1.38 (0.074) | 1.41 (0.090) | 1.54 (0.11) | 1.68 (0.13) |
|  | MSMC2 | 1 | 0.87 (0.064) | 0.89 (0.067) | .99 (0.15) | 1.13 (0.30) |
| 100 kb | eSMC | 1 | 0.95 (0.047) | 0.95 (0.051) | 0.98 (0.070) | 1.0 (0.12) |
|  | MSMC | 1 | 1.36 (0.048) | 1.36 (0.062) | 1.40 (0.093) | 1.49 (0.12) |
|  | MSMC2 | 1 | 0.87 (0.056) | 0.88 (0.050) | 0.91 (0.079) | 0.91 (0.073) |

It is, in some instances, possible to compensate for a 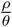 ratio that is not ideal by increasing the number of iterations. Indeed, for eSMC, the demographic history is better inferred (Supplementary Figure 22), although the correct recombination rate cannot be retrieved (Table 2). MSMC is able to better infer the correct recombination rate than other methods despite 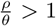 *>* 1, but poorly estimates the demographic history. The past demographic inferences obtained using MSMC2 and SMC++ are not improved when increasing the number of iterations (see Supplementary Figure 22 and Table 2).

### Simulation results under hypothesis violation

#### Imperfect SNP calling

We analyze simulated sequences that have been modified by removing and/or adding SNPs using the different SMC methods. We find that, when using MSMC2, eSMC and MSMC, having more than 10% of spurious SNPs (*e.g*. no quality filtering) can lead to a strong over-estimation of population size in recent time but that missing SNPs have no effects on inferences in the far past and only mild effects on inferences of recent time (see Figure 5 for MSMC2, Supplementary Figures 23 and 24 for eSMC and MSMC respectively). The mean square error is displayed in Supplementary Table 4.

As complementary analyses we analyze simulated sequences with a Minor Allele Frequency (MAF) threshold. Results are shown in Supplementary Figure 25. The more SNPs are removed, the poorer the estimations in recent time (Supplementary Figure 25), demonstrating the impact of severe ascertainment bias.

### Specific scaffold parameters

We simulate sequence data where scaffolds have either been simulated with the same recombination and mutation rates or with different recombination and mutation rates. Data sets are then analyzed assuming scaffolds share or do not share the same recombination and mutation rates. We can see in Figure 6 (and Supplementary Table 5) that when scaffolds all share the same parameter values, estimated demography is accurate both when the analysis assumed shared or differing mutation and recombination rates. However, when scaffolds are simulated with different parameter values, analyzing them under the assumption that they have the same mutation and recombination rates leads to poor estimations. Assuming scaffolds do not share recombination and mutation rates does improve the results somewhat, but the estimations remain less accurate than when scaffolds all share with same parameter values. If only the recombination rate changes from one scaffold to another, the demographic history is only slightly biased, whereas, if the mutation rate changes from one scaffold to the other, demographic history is poorly estimated.

Even if chromosomes are fully assembled, assuming we here have one scaffold of 40 Mb (chromosome fully assembled), there may be variations of the recombination rate along the sequence, however this seems of little consequence when applying eSMC. As can be seen in Supplementary Figure 26, the demographic scenario is well inferred, despite an increase in variance and a smooth “wave” shaped demographic history when sequences simulated with varying recombination rates are compared to those with a fixed recombination rate throughout the genome. Overall we see that when recombination rate is heterogeneous along the genome by a factor 5, it is not untypical to falsely estimate a two-fold variation of Ne even though the true Ne is constant in time.

### How transposable elements bias inference

Transposable elements (TEs) are present in most species, and are (if detected) taken into account as missing data by SMC methods (Schiffels and Durbin, 2014)). Depending on how TEs affect the data set, we find that methods are more or less sensitive to TEs. If TEs are unmapped/removed from the data set, there does not appear to be any bias in the estimated demographic history when using eSMC (see Figure 7 and Supplementary Table 6), but there is an overestimation of 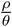 (see Table 3). We find that, the higher the proportion of sequences removed, the more 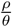 is over-estimated. For a fixed amount of missing/removed data, the smaller the sequences that are removed, the more 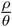 is over-estimated (Table 3). If TEs are present but unmasked in the data set (and thus not accounted for missing data by the model (Schiffels and Durbin, 2014)), we find that this is equivalent to a faulty calling of SNPs, in which SNPs are missing, hence resulting in demographic history estimations by eSMC similar to those observed in Figure 5a. However, if the size of unmasked TEs increases, different results are obtained (see Supplementary Figures 27 and 28). Indeed, in recent times there is a strong underestimation of population size and the model fails to capture the correct demographic history. The longer the TEs are, the stronger the effect on the estimated demographic history. Similar results are obtained with MSMC (Supplementary Figures 29, 30 and 31) and MSMC2 (Supplementary Figures 32, 33 and 34). However, when TEs are detected and correctly masked, there is no effect on demographic inferences (Supplementary Figures 35 and 36).

## Discussion

Throughout this work we have outlined the limits of PSMC’ and MSMC methodologies, which had, until now, not been clearly defined. We find that, in most cases, if enough genealogies (*i.e*. data) are inputted then the demographic history is accurately estimated, tending to results obtained previously (Chikhi et al., 2018; Gattepaille et al., 2016), however, we find that the amount of data required for an accurate fit depends on the underlying demographic scenario. The differences with previous works stems from estimations being made using the actual series of coalescence times (Chikhi et al., 2018; Gattepaille et al., 2016), whereas we use the series of hidden states built from the discretization of time summarized in a simple matrix. We also find that some scenarios are better retrieved when using either MSMC or methods based on PSMC’, indicating that there are complementary convergence properties between these methodologies.

We develop a method to indicate if the amount of data is enough to retrieve a specific scenario, notably by calculating the coefficient of variation of the transition matrix using either real or simulated data, and therefore offer guidelines to build appropriate data sets (see also Supplementary Figure 8). Our approach can also be used to infer demographic history given an ARG (using trees in Newick format or sequences of coalescence events), independently of how the ARG has been estimated. Our results suggest that whole genome polymorphism data can be summarized in a transition matrix based on the SMC theory to estimate demographic history of panmitic populations. As new methods can infer genealogies better and faster (Kelleher, Wong, et al., 2019; Mirzaei and Wu, 2017; Palamara et al., 2018; Speidel et al., 2019), the estimated transition matrix could become a powerful summary statistic in the future. HMM can be a computational burden depending on the model and model parameters, and estimating genealogy through more efficient methods would still allow the use of SMC theory for parameter estimation or hypothesis testing (as in Gattepaille et al. (2016), Johndrow and Palacios (2019), and P Wang et al. (2018)). In addition, using the work of K Wang et al. (2020), one could (to some extent Kim et al. (2020)) extend our approach to account for population structure and migration.

We have also demonstrated that the power of PSMC’, MSMC, and other SMC-based methods, rely on their ability to correctly infer the genealogies along the sequence (*i.e*. the Ancestral Recombination Graph or ARG). The accuracy of ARG inference by SMC methods, however, depends on the ratio of the recombination over the mutation rate 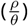 As this rate increases, estimations lose accuracy. Specifically, increasing 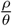 leads to an over-estimation of transitions on the diagonal, which explains the underestimation of the recombination rate and inaccurate demographic history estimations, as shown in Sellinger et al. (2020) and Terhorst, Kamm, et al. (2017). As a way around this issue, in some cases it is possible to obtain better results by increasing the number of iterations. MSMC’s demographic inference is more sensitive to 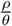 but the quality of the estimation of the ratio itself is less affected. This once again shows the complementarity of PSMC’ and MSMC. If the variable of interest is 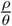 then MSMC should be used, but if the demographic history is of greater importance, PSMC’-based methods should be used. The amplitude of population size variation also influences the estimation of hidden states along the sequences, with high amplitudes leading to a poor estimation of the transition matrix, distorting the inferred demography. We find that increasing the size of the time window increases the variance of the estimations, despite using the same number of parameters, as this results in a small under-estimation of 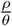 In addition the complete and incomplete Baum-Welch algorithms lead to identical results, demonstrating that all the information required for the inference is in the estimated transition matrix.

Finally, we explored how imperfect data sets (due to errors in SNP calling, the presence of transposable elements and existing variation in recombination and mutation rates) could affect the inferences obtained using SMC-based methods. We show that a data set with more than 10% of spurious SNPs will lead to poor estimations of the demographic history, whereas randomly removed SNPs (*i.e*. missing SNPs) have a lesser effect on inferences. It is thus better to be stringent during SNP calling, as SNPs is worse than missing SNPs. Note, however, that this consideration is valid for demographic inference under a neutral model of evolution, while biases in SNP calling also affect the inference of selection (especially for conserved genes under purifying selection). However, if missing SNPs are structured along the sequence (as would be the case with unmasked TEs), there is a strong effect on inference. If TEs are correctly detected and masked, there is no effect on demographic inferences. It is therefore recommended that checks should be run to detect regions with abnormal distributions of SNPs along the genome. Surprisingly, simulation results suggest that removing random pieces of sequences have no impact on the estimated demographic history. Taking this into account, when seeking to infer demographic history, it seems better to remove sections of sequences than to introduce sequences with SNP call errors or abnormal SNP distributions. However, removing sequences leads to an over-estimation of 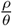 which seems to depend on the number and size of the removed sections. The removal of a few, albeit long sequences, will have almost no impact, whereas removing many short sections of the sequences will lead to a large overestimation of 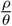 This consequence could provide an explanation for the frequent overestimation of 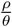 when compared to empirical measures of the ratio of recombination and mutation rates 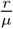 This implies, that in some cases, despite an inferred 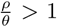 *>* 1, the inferred demographic history can surprisingly be trusted. Note also that as discussed in Sellinger et al. (2020), the discrepancy between 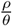 and 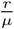 can be due to life history traits such as selfing or dormancy.

Simulation results suggest that any variation of the recombination rate along the sequence does not strongly bias demographic inference but slightly increases the variance of the results and leads to small waves in the demographic history (as a consequence of erroneously estimated hidden state transition events because of the non constant recombination rate along the sequence), as expected from previous works (H Li and Durbin, 2011). However, unlike Li and Durbin’s results (H Li and Durbin, 2011), if scaffolds do not share similar rates of mutation and recombination, but are analyzed together assuming that they do, estimations will be very poor. This could be due to the variation of mutation rate being within a scaffold in their study and the discrepancy between out and their results could suggest analyses based on longer scaffolds to be more robust. However, this problem can be avoided if each scaffold is assumed to have its own parameter values, although this would increase computation time, it could provide useful insight in unveiling any variation in molecular forces along the genome, albeit in a coarser way than in Barroso et al. (2019). As we found that non-accounted variation of the recombination rate along the sequence can lead to a spurious two-fold variation of population size, we here provide guidelines to test if small detected variations of population size are to be trusted. Since the consequecnes of a varying recombination rate might depend on the topology of the recombination map, one first needs estimate the recombination map (*e.g*. using iSMC (Barroso et al., 2019)). If problematic regions are found they can be removed with almost no negative impact on the estimated demography (Figure 7). Otherwise,the recombination map can be used to simulate sequences *e.g*. using scrm (Staab et al., 2015)), which can be compared to results obtained for a constant recombination rate. Analyses can be run on both data sets to quantify the effect of the recombination map.

### Guidelines when applying SMC-based methods

Our aim through this work is to provide guidelines to optimize the use of SMC-based methods for inference. First, if the data set is not yet built, but there is some intuition concerning the demographic history and knowledge of some genomic properties of a species (*e.g*. recombination and mutation rates), we recommend simulating a data set corresponding to the potential scenarios. From these simulations, the transition matrix for PSMC’ or MSMC-based methods can be built using the R package eSMC2. The results obtained can guide users when it comes to the amount and quality of data needed (sequence size and copy number) for a good inference. Beyond being used to guide the building of data sets, it is possible to assess trustworthiness of results obtained using SMC-based methods on existing data sets. If the estimated transition matrix is empty in some places (*i.e*. no observed transition event between two specific hidden states; white squares in Figure 2), it could suggest a lack of data and/or strong variation of the population size somewhere in time. In order to test the accuracy of the inferred demography, the estimated demographic history can be retrieved and used to simulate a data set with more sequences and/or simulate a demographic history with a higher amplitude than the estimated one. The SMC method can then be run on the simulated data in order to check whether using more data results in a matching scenario or if a higher amplitude of population size can indeed be inferred, in which cases the initial results are most probably trustworthy.

As mentioned above, it is better to sequence fewer individuals, but to have data of better quality. It is also important to note, that more data is not necessarily always better, especially if there is a risk of spurious SNPs (see Figure 5). In some cases, methods are limited by their own theoretical framework, hence no given data set will ever allow a correct demographic inference. In such cases, other methods based on a different theoretical frameworks (*e.g*. SFS and ABC) might perform better (Beichman et al., 2017; Schraiber and Akey, 2015).

### Concluding remarks

Here we present a simple method to help assess how accurate inferences obtained using PSMC’ and MSMC would be when applied to data sets with suspected flaws or limitations. We also provide new interpretations of results obtained when hypotheses are known to be violated, and thus offer an explanation as to why results sometimes deviate from expectations (*e.g*. when the estimated ratio of recombination over mutation is larger than the one measured experimentally). We propose guidelines for building/evaluating data sets when using SMC-based models, as well as a method which can be used to estimate the demographic history and recombination rate given a genealogy (in the same spirit as Popsicle (Gattepaille et al., 2016)). The estimated transition matrix is introduced as a summary statistic, which can be used to recover demographic history (more precisely the Inverse Instantaneous Coalescence Rate interpretation of population size variation, when assuming a panmictic population (Chikhi et al., 2018; Rodriguez et al., 2018)). This statistic could, in future, be used in scenarios with migration, without the computational load of Hidden Markov models. When faced with complex demographic histories, or 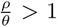, we show that there are strategies that would allow those wishing to use SMC methodology to make the best use of their data.

## Supplementary material

Scripts and codes are available online: https://www.biorxiv.org/content/10.1101/2020.07.23.217091v2.supplementary-material https://github.com/TPPSellinger/eSMC2

## Supporting information

command lines for simulation

command lines for analyses

Contains supplementary figures and tables

contains all mean square error values

## Acknowledgements

This work was funded by Deutsche Forschungsgemeinschaft, project number 317616126 (TE809/7-1) to AT. DAA was funded by the Alexander von Humboldt Stiftung. Version 3 of this preprint has been peer-reviewed and recommended by Peer Community In Evolutionary Biology (https://doi.org/10.24072/pci.evolbiol.100115)

## Appendix

Link to appendix 1:

https://www.biorxiv.org/content/biorxiv/early/2020/09/21/2020.07.23.217091/DC3/embed/media-3.zip?download=true

Link to appendix 2:

https://www.biorxiv.org/content/biorxiv/early/2020/09/21/2020.07.23.217091/DC4/embed/media-4.zip?download=true

Link to Supplementary Figures:

https://www.biorxiv.org/content/biorxiv/early/2020/09/21/2020.07.23.217091/DC1/embed/media-1.pdf?download=true

Link to Supplementary Tables:

https://www.biorxiv.org/content/biorxiv/early/2020/09/21/2020.07.23.217091/DC2/embed/media-2.zip?download=true

## References

Barroso GV, N Puzovic, and JY Dutheil (Nov. 2019). Inference of recombination maps from a single pair of genomes and its application to ancient samples. PLOS GENETICS 15. issn: 1553-7404. doi: 10.1371/journal.pgen.1008449;10.1371/journal.pgen.1008449.r001;10.1371/journal.pgen.1008449.r002;10.1371/journal.pgen.1008449.r003;10.1371/journal.pgen.1008449.r004.

Beeravolu CR, MJ Hickerson, LAF Frantz, and K Lohse (Sept. 2018). ABLE: blockwise site frequency spectra for inferring complex population histories and recombination. Genome Biology 19. issn: 1474-760X. doi: 10.1186/s13059-018-1517-y.

Beichman AC, TN Phung, and KE Lohmueller (Nov. 2017). Comparison of Single Genome and Allele Frequency Data Reveals Discordant Demographic Histories. G3-GENES GENOMES GENETICS 7, 3605–3620. issn: 2160-1836. doi: 10.1534/g3.117.300259.

Bergstrom A, SA McCarthy, R Hui, MA Almarri, Q Ayub, P Danecek, Y Chen, S Felkel, P Hallast, J Kamm, H Blanche, JF Deleuze, H Cann, S Mallick, D Reich, MS Sandhu, P Skoglund, A Scally, Y Xue, R Durbin, and C Tyler-Smith (Mar. 2020). Insights into human genetic variation and population history from 929 diverse genomes. SCIENCE 367, 1339+. issn: 0036-8075. doi: 10.1126/science.aay5012.

Browning SR, BL Browning, Y Zhou, S Tucci, and JM Akey (Mar. 2018). Analysis of Human Sequence Data Reveals Two Pulses of Archaic Denisovan Admixture. CELL 173, 53+. issn: 0092-8674. doi: 10.1016/j.cell.2018.02.031.

Cao J, K Schneeberger, S Ossowski, T Guenther, S Bender, J Fitz, D Koenig, C Lanz, O Stegle, C Lippert, X Wang, F Ott, J Mueller, C Alonso-Blanco, K Borgwardt, KJ Schmid, and D Weigel (Oct. 2011). Whole-genome sequencing of multiple Arabidopsis thaliana populations. Nature Genetics 43, 956–U60. issn: 1061-4036. doi: 10.1038/ng.911.

Chang D and B Shapiro (Feb. 2016). Using ancient DNA and coalescent-based methods to infer extinction. Biology Letters 12. issn: 1744-9561. doi: 10.1098/rsbl.2015.0822.

Chikhi L, W Rodriguez, S Grusea, P Santos, S Boitard, and O Mazet (Jan. 2018). The IICR (inverse instantaneous coalescence rate) as a summary of genomic diversity: insights into demographic inference and model choice. Heredity 120, 13–24. issn: 0018-067X. doi: 10.1038/s41437-017-0005-6.

Choo SW, M Rayko, TK Tan, R Hari, A Komissarov, WY Wee, AA Yurchenko, S Kliver, G Tamazian, A Antunes, RK Wilson, WC Warren, KP Koepfli, P Minx, K Krasheninnikova, A Kotze, DL Dalton, E Vermaak, IC Paterson, P Dobrynin, FT Sitam, JJ Rovie-Ryan, WE Johnson, AM Yusoff, SJ Luo, KV Karuppannan, G Fang, D Zheng, MB Gerstein, L Lipovich, S. O’Brien, and GJ Wong (Oct. 2016). Pangolin genomes and the evolution of mammalian scales and immunity. GENOME RESEARCH 26, 1312–1322. issn: 1088-9051. doi: 10.1101/gr.203521.115.

Ekblom R, B Brechlin, J Persson, L Smeds, M Johansson, J Magnusson, O Flagstad, and H Ellegren (Dec. 2018). Genome sequencing and conservation genomics in the Scandinavian wolverine population. Conservation Biology 32, 1301–1312. issn: 0888-8892. doi: 10.1111/cobi.13157.

Ewing AD (Dec. 2015). Transposable element detection from whole genome sequence data. MOBILE DNA 6. issn: 1759-8753. doi: 10.1186/s13100-015-0055-3.

Fulgione A, M Koornneef, F Roux, J Hermisson, and AM Hancock (Mar. 2018). Madeiran Arabidopsis thaliana Reveals Ancient Long-Range Colonization and Clarifies Demography in Eurasia. Molecular Biology and Evolution 35, 564–574. issn: 0737-4038. doi: 10.1093/molbev/msx300.

Gattepaille L, T Guenther, and M Jakobsson (Nov. 2016). Inferring Past Effective Population Size from Distributions of Coalescent Times. Molecular Biology and Evolution 204, 1191+. issn: 0016-6731. doi: 10.1534/genetics.115.185058.

Gaut BS, DK Seymour, Q Liu, and Y Zhou (Aug. 2018). Demography and its effects on genomic variation in crop domestication. Nature Plants 4, 512–520. issn: 2055-026X. doi: 10.1038/s41477-018-0210-1.

Hawks J (Jan. 2017). Introgression Makes Waves in Inferred Histories of Effective Population Size. HUMAN BIOLOGY 89, 67–80. issn: 0018-7143. doi: 10.13110/humanbiology.89.1.04.

Hecht LBB, PC Thompson, and BM Rosenthal (Oct. 2018). Comparative demography elucidates the longevity of parasitic and symbiotic relationships. PROCEEDINGS OF THE ROYAL SOCIETY B-BIOLOGICAL SCIENCES 285. issn: 0962-8452. doi: 10.1098/rspb.2018.1032.

Hendricks S, EC Anderson, T Antao, L Bernatchez, BR Forester, B Garner, BK Hand, PA Hohenlohe, M Kardos, B Koop, A Sethuraman, RS Waples, and G Luikart (Sept. 2018). Recent advances in conservation and population genomics data analysis. Evolutionary Applications 11, 1197–1211. issn: 1752-4571. doi: 10.1111/eva.12659.

Hobolth A and JL Jensen (Dec. 2014). Markovian approximation to the finite loci coalescent with recombination along multiple sequences. THEORETICAL POPULATION BIOLOGY 98, 48–58. issn: 0040-5809. doi: 10.1016/j.tpb.2014.01.002.

Johndrow JE and JA Palacios (Feb. 2019). Exact limits of inference in coalescent models. Theoretical Population Biology 125, 75–93. issn: 0040-5809. doi: 10.1016/j.tpb.2018.11.004.

Kardos M, A Qvarnstrom, and H Ellegren (Mar. 2017). Inferring Individual Inbreeding and Demographic History from Segments of Identity by Descent in Ficedula Flycatcher Genome Sequences. GENETICS 205, 1319–1334. issn: 0016-6731. doi: 10.1534/genetics.116.198861.

Kelleher J, AM Etheridge, and G McVean (May 2016). Efficient Coalescent Simulation and Genealogical Analysis for Large Sample Sizes. PLOS COMPUTATIONAL BIOLOGY 12. doi: 10.1371/journal.pcbi.1004842.

Kelleher J, Y Wong, AW Wohns, C Fadil, PK Albers, and G McVean (Nov. 2019). Inferring whole-genome histories in large population datasets (vol 51, pg 1330, 2019). NATURE GENETICS 51, 1660. issn: 1061-4036. doi: 10.1038/s41588-019-0523-7.

Kim Y, F Koehler, A Moitra, E Mossel, and G Ramnarayan (Apr. 2020). How Many Subpopulations Is Too Many? Exponential Lower Bounds for Inferring Population Histories. JOURNAL OF COMPUTATIONAL BIOLOGY 27, 613–625. issn: 1066-5277. doi: 10.1089/cmb.2019.0318.

Kofler R (Apr. 2018). SimulaTE: simulating complex landscapes of transposable elements of populations. BIOINFORMATICS 34, 1419–1420. issn: 1367-4803. doi: 10.1093/bioinformatics/btx772.

Lau SCY, NG Wilson, CNS Silva, and JM Strugnell (2020). Detecting glacial refugia in the Southern Ocean. ECOGRAPHY. issn: 0906-7590. doi: 10.1111/ecog.04951.

Li H and R Durbin (July 2011). Inference of human population history from individual whole-genome sequences. Nature 475, 493–U84. issn: 0028-0836. doi: 10.1038/nature10231.

Li S, B Li, C Cheng, Z Xiong, Q Liu, J Lai, HV Carey, Q Zhang, H Zheng, S Wei, H Zhang, L Chang, S Liu, S Zhang, B Yu, X Zeng, Y Hou, W Nie, Y Guo, T Chen, J Han, J Wang, J Wang, C Chen, J Liu, PJ Stambrook, M Xu, G Zhang, MTP Gilbert, H Yang, ED Jarvis, J Yu, and J Yan (2014). Genomic signatures of near-extinction and rebirth of the crested ibis and other endangered bird species. GENOME BIOLOGY 15. issn: 1474-760X. doi: 10.1186/s13059-014-0557-1.

Lynch M, R Gutenkunst, M Ackerman, K Spitze, Z Ye, T Maruki, and Z Jia(May 2017). Population Genomics of Daphnia pulex. Molecular Biology and Evolution 206, 315–332. issn: 0016-6731. doi: 10.1534/genetics.116.190611.

Lynch M, B Haubold, P Pfaffelhuber, and T Maruki (Jan. 2020). Inference of Historical Population-Size Changes with Allele-Frequency Data. G3-GENES GENOMES GENETICS 10, 211–223. issn: 2160-1836. doi: 10.1534/g3.119.400854.

Malaspinas AS, MC Westaway, C Muller, VC Sousa, O Lao, I Alves, A Bergstrom, G Athanasiadis, JY Cheng, JE Crawford, TH Heupink, E Macholdt, S Peischl, S Rasmussen, S Schiffels, S Subramanian, JL Wright, A Albrechtsen, C Barbieri, I Dupanloup, A Eriksson, A Margaryan, I Moltke, I Pugach, TS Korneliussen, IP Levkivskyi, JV Moreno-Mayar, S Ni, F Racimo, M Sikora, Y Xue, FA Aghakhanian, N Brucato, S Brunak, PF Campos, W Clark, S Ellingvag, G Fourmile, P Gerbault, D Injie, G Koki, M Leavesley, B Logan, A Lynch, EA Matisoo-Smith, PJ McAllister, AJ Mentzer, M Metspalu, AB Migliano, L Murgha, ME Phipps, W Pomat, D Reynolds, FX Ricaut, P Siba, MG Thomas, T Wales, CM Wall, SJ Oppenheimer, C Tyler-Smith, R Durbin, J Dortch, A Manica, MH Schierup, RA Foley, MM Lahr, C Bowern, JD Wall, T Mailund, M Stoneking, R Nielsen, MS Sandhu, L Excoffier, DM Lambert, and E Willerslev (Oct. 2016). A genomic history of Aboriginal Australia. NATURE 538, 207+. issn: 0028-0836. doi: 10.1038/nature18299.

Marjoram P and J Wall (Mar. 2006). Fast “coalescent” simulation. BMC Genetics 7. issn: 1471-2156. doi: 10.1186/1471-2156-7-16.

Mather N, SM Traves, and SYW Ho (Jan. 2020). A practical introduction to sequentially Markovian coalescent methods for estimating demographic history from genomic data. ECOLOGY AND EVOLUTION 10, 579–589. issn: 2045-7758. doi: 10.1002/ece3.5888.

Mattle-Greminger MP, TB Sonay, A Nater, M Pybus, T Desai, G de Valles, F Casals, A Scally, J Bertranpetit, T Marques-Bonet, CP van Schaik, M Anisimova, and M Kruetzen (Nov. 2018). Genomes reveal marked differences in the adaptive evolution between orangutan species. Genome Biology 19. issn: 1474-760X. doi: 10.1186/s13059-018-1562-6.

Mazet O, W Rodriguez, S Grusea, S Boitard, and L Chikhi (Apr. 2016). On the importance of being structured: instantaneous coalescence rates and human evolution-lessons for ancestral population size inference? Heredity 116, 362–371. issn: 0018-067X. doi: 10.1038/hdy.2015.104.

McVean G and N Cardin (July 2005). Approximating the coalescent with recombination. Philosophical Transactions of the Royal Society B-Biological Sciences 360, 1387–1393. issn: 0962-8436. doi: 10.1098/rstb.20053.1673.

Mirzaei S and Y Wu (Apr. 2017). RENT plus: an improved method for inferring local genealogical trees from haplotypes with recombination. BIOINFORMATICS 33, 1021–1030. issn: 1367-4803. doi: 10.1093/bioinformatics/btw735.

Nadachowska-Brzyska K, R Burri, L Smeds, and H Ellegren (Mar. 2016). PSMC analysis of effective population sizes in molecular ecology and its application to black-and-white Ficedula flycatchers. Molecular Ecology 25, 1058–1072. issn: 0962-1083. doi: 10.1111/mec.13540.

Nakagome S, RR Hudson, and A Di Rienzo (Feb. 2019). Inferring the model and onset of natural selection under varying population size from the site frequency spectrum and haplotype structure. PROCEEDINGS OF THE ROYAL SOCIETY B-BIOLOGICAL SCIENCES 286. issn: 0962-8452. doi: 10.1098/rspb.2018.2541.

Nelson MG, RS Linheiro, and CM Bergman (Aug. 2017). McClintock: An Integrated Pipeline for Detecting Transposable Element Insertions in Whole-Genome Shotgun Sequencing Data. G3-GENES GENOMES GENETICS 7, 2763–2778. issn: 2160-1836. doi: 10.1534/g3.117.043893.

Oh KP, CL Aldridge, JS Forbey, CY Dadabay, and SJ Oyler-McCance (July 2019). Conservation Genomics in the Sagebrush Sea: Population Divergence, Demographic History, and Local Adaptation in Sage-Grouse (Centrocercus spp.) GENOME BIOLOGY AND EVOLUTION 11, 2023–2034. issn: 1759-6653. doi: 10.1093/gbe/evz112.

Palacios JA, J Wakeley, and S Ramachandran (Sept. 2015). Bayesian Nonparametric Inference of Population Size Changes from Sequential Genealogies. Genetics 201, 281+. issn: 0016-6731. doi: 10.1534/genetics.115.177980.

Palamara PF, J Terhorst, YS Song, and AL Price (Sept. 2018). High-throughput inference of pairwise coalescence times identifies signals of selection and enriched disease heritability. NATURE GENETICS 50, 1311+. issn: 1061-4036. doi: 10.1038/s41588-018-0177-x.

Palkopoulou E, M Lipson, S Mallick, S Nielsen, N Rohland, S Baleka, E Karpinski, AM Ivancevici, TH To, D Kortschak, JM Raison, Z Qu, TJ Chin, KW Alt, S Claesson, L Dalen, RDE MacPhee, H Meller, AL Rocar, OA Ryder, D Heiman, S Young, M Breen, C Williams, BL Aken, M Ruffier, E Karlsson, J Johnson, F Di Palma, J Alfoldi, DL Adelsoni, T Mailund, K Munch, K LindbladToh, M Hofreiter, H Poinar, and D Reich (Mar. 2018). A comprehensive genomic history of extinct and living elephants. Proceedings of the National Academy of Sciences of the United States of America 115, E2566–E2574. issn: 0027-8424. doi: 10.1073/pnas.1720554115.

Palkopoulou E, S Mallick, P Skoglund, J Enk, N Rohland, H Li, A Omrak, S Vartanyan, H Poinar, A Gotherstrom, D Reich, and L Dalen (May 2015). Complete Genomes Reveal Signatures of Demographic and Genetic Declines in the Woolly Mammoth. Current Biology 25, 1395–1400. issn: 0960-9822. doi: 10.1016/j.cub.2015.04.007.

Patton AH, MJ Margres, AR Stahlke, S Hendricks, K Lewallen, RK Hamede, M Ruiz-Aravena, O Ryder, H McCallum I, ME Jones, PA Hohenlohe, and A Storfer(Dec. 2019). Contemporary Demographic Reconstruction Methods Are Robust to Genome Assembly Quality: A Case Study in Tasmanian Devils. MOLECULAR BIOLOGY AND EVOLUTION 36, 2906–2921. issn: 0737-4038. doi: 10.1093/molbev/msz191.

Pfeifer SP (Feb. 2017). From next-generation resequencing reads to a high-quality variant data set. HEREDITY 118, 111–124. issn: 0018-067X. doi: 10.1038/hdy.2016.102.

Platt II RN, L Blanco-Berdugo, and DA Ray (Feb. 2016). Accurate Transposable Element Annotation Is Vital When Analyzing New Genome Assemblies. GENOME BIOLOGY AND EVOLUTION 8, 403–410. issn: 1759-6653. doi: 10.1093/gbe/evw009.

Prado-Martinez J, PH Sudmant, JM Kidd, H Li, JL Kelley, B Lorente-Galdos, KR Veeramah, AE Woerner, T. O’Connor, G Santpere, A Cagan, C Theunert, F Casals, H Laayouni, K Munch, A Hobolth, AE Halager, M Malig, J Hernandez-Rodriguez, I Hernando-Herraez, K Pruefer, M Pybus, L Johnstone, M Lachmann, C Alkan, D Twigg, N Petit, C Baker, F Hormozdiari, M Fernandez-Callejo, M Dabad, ML Wilson, L Stevison, C Camprubi, T Carvalho, A Ruiz-Herrera, L Vives, M Mele, T Abello, I Kondova, RE Bontrop, A Pusey, F Lankester, JA Kiyang, RA Bergl, E Lonsdorf, S Myers, M Ventura, P Gagneux, D Comas, H Siegismund, J Blanc, L Agueda-Calpena, M Gut, L Fulton, SA Tishkoff, JC Mullikin, RK Wilson, IG Gut, MK Gonder, OA Ryder, BH Hahn, A Navarro, JM Akey, J Bertranpetit, D Reich, T Mailund, MH Schierup, C Hvilsom, AM Andres, JD Wall, CD Bustamante, MF Hammer, EE Eichler, and T Marques-Bonet (July 2013). Great ape genetic diversity and population history. NATURE 499, 471–475. issn: 0028-0836. doi: 10.1038/nature12228.

Rodriguez W, O Mazet, S Grusea, A Arredondo, JM Corujo, S Boitard, and L Chikhi (Dec. 2018). The IICR and the non-stationary structured coalescent: towards demographic inference with arbitrary changes in population structure. Heredity 121, 663–678. issn: 0018-067X. doi: 10.1038/s41437-018-0148-0.

Schiffels S and R Durbin (Aug. 2014). Inferring human population size and separation history from multiple genome sequences. Nature Genetics 46, 919–925. issn: 1061-4036. doi: 10.1038/ng.3015.

Schraiber JG and JM Akey (Dec. 2015). Methods and models for unravelling human evolutionary history. NATURE REVIEWS GENETICS 16, 727–740. issn: 1471-0056. doi: 10.1038/nrg4005.

Schrider DR, AG Shanku, and AD Kern (Nov. 2016). Effects of Linked Selective Sweeps on Demographic Inference and Model Selection. GENETICS 204, 1207+. issn: 0016-6731. doi: 10.1534/genetics.116.190223.

Sellinger TPP, D Abu Awad, M Moest, and A Tellier (Apr. 2020). Inference of past demography, dormancy and self-fertilization rates from whole genome sequence data. PLOS GENETICS 16. issn: 1553-7404. doi: 10.1371/journal.pgen.1008698;10.1371/journal.pgen.1008698.r001;10.1371/journal.pgen.1008698.r002;10.1371/journal.pgen.1008698.r003;10.1371/journal.pgen.1008698.r004;10.1371/journal.pgen.1008698.r005;10.1371/journal.pgen.1008698.r006.

Sheehan S, K Harris, and YS Song (July 2013). Estimating Variable Effective Population Sizes from Multiple Genomes: A Sequentially Markov Conditional Sampling Distribution Approach. Molecular Biology and Evolution 194, 647+. issn: 1943-2631. doi: 10.1534/genetics.112.149096.

Sheehan S and YS Song (Mar. 2016). Deep Learning for Population Genetic Inference. PLOS Computational Biology 12. issn: 1553-734X. doi: 10.1371/journal.pcbi.1004845.

Slatkin M (Dec. 2016). Statistical methods for analyzing ancient DNA from hominins. CURRENT OPINION IN GENETICS & DEVELOPMENT 41, 72–76. issn: 0959-437X. doi: 10.1016/j.gde.2016.08.004.

Smith CCR and SM Flaxman (Jan. 2020). Leveraging whole genome sequencing data for demographic inference with approximate Bayesian computation. MOLECULAR ECOLOGY RESOURCES 20, 125–139. issn: 1755-098X. doi: 10.1111/1755-0998.13092.

Speidel L, M Forest, S Shi, and SR Myers (Sept. 2019). A method for genome-wide genealogy estimation for thousands of samples. NATURE GENETICS 51, 1321+. issn: 1061-4036. doi: 10.1038/s41588-019-0484-x.

Spence JP, M Steinrucken, J Terhorst, and YS Song (Dec. 2018). Inference of population history using coalescent HMMs: review and outlook. Current Opinion in Genetics & Development 53, 70–76. issn: 0959-437X. doi: 10.1016/j.gde.2018.07.002.

Staab PR, S Zhu, D Metzler, and G Lunter (May 2015). scrm: efficiently simulating long sequences using the approximated coalescent with recombination. Bioinformatics 31, 1680–1682. issn: 1367-4803. doi: 10.1093/bioinformatics/btu861.

Stam R, T Nosenko, AC Hoerger, W Stephan, M Seidel, JMM Kuhn, G Haberer, and A Tellier (Dec. 2019). The de Novo Reference Genome and Transcriptome Assemblies of the Wild Tomato Species Solanum chilense Highlights Birth and Death of NLR Genes Between Tomato Species. G3-GENES GENOMES GENETICS 9, 3933–3941. issn: 2160-1836. doi: 10.1534/g3.119.400529.

Steinrucken M, J Kamm, JP Spence, and YS Song (Aug. 2019). Inference of complex population histories using whole-genome sequences from multiple populations. PROCEEDINGS OF THE NATIONAL ACADEMY OF SCIENCES OF THE UNITED STATES OF AMERICA 116, 17115–17120. issn: 0027-8424. doi: 10.1073/pnas.1905060116.

Terhorst J, JA Kamm, and YS Song (Feb. 2017). Robust and scalable inference of population history froth hundreds of unphased whole genomes. Nature Genetics 49, 303–309. issn: 1061-4036. doi: 10.1038/ng.3748.

Terhorst J and YS Song (June 2015). Fundamental limits on the accuracy of demographic inference based on the sample frequency spectrum. Proceedings of the National Academy of Sciences of the United States of America 112, 7677–7682. issn: 0027-8424. doi: 10.1073/pnas.1503717112.

Waltoft BL and A Hobolth (June 2018). Non-parametric estimation of population size changes from the site frequency spectrum. Statistical Applications in Genetics and Molecular Biology 17. issn: 2194-6302. doi: 10.1515/sagmb-2017-0061.

Wang K, I Mathieson, J O’Connell, and S Schiffels (Mar. 2020). Tracking human population structure through time from whole genome sequences. PLOS GENETICS 16. issn: 1553-7404. doi: 10.1371/journal.pgen.1008552.

Wang P, H Yao, KJ Gilbert, Q Lu, Y Hao, Z Zhang, and N Wang (Dec. 2018). Glaciation-based isolation contributed to speciation in a Palearctic alpine biodiversity hotspot: Evidence from endemic species. Molecular Phylogenetics and Evolution 129, 315–324. issn: 1055-7903. doi: 10.1016/j.ympev.2018.09.006.

Williams RC, MB Blanco, JW Poelstra, KE Hunnicutt, AA Comeault, and AD Yoder (Jan. 2020). Conservation genomic analysis reveals ancient introgression and declining levels of genetic diversity in Madagascar’s hibernating dwarf lemurs. HEREDITY 124, 236–251. issn: 0018-067X. doi: 10.1038/s41437-019-0260-9.

Wiuf C and J Hein (June 1999). Recombination as a point process along sequences. Theoretical Population Biology 55, 248–259. issn: 0040-5809. doi: 10.1006/tpbi.1998.1403.

Yew CW, D Lu, L Deng, LP Wong, RTH Ong, Y Lu, X Wang, Y Yunus, F Aghakhanian, SS Mokhtar, MZ Hoque, CLY Voo, TA Rahman, J Bhak, ME Phipps, S Xu, YY Teo, SV Kumar, and BP Hoh (Feb. 2018). Genomic structure of the native inhabitants of Peninsular Malaysia and North Borneo suggests complex human population history in Southeast Asia. Human Genetics 137, 161–173. issn: 0340-6717. doi: 10.1007/s00439-018-1869-0.

