## Supplementary material for "Limits and Convergence properties of the Sequentially Markovian Coalescent": command lines for simulation: Figure_1.R

```
###################
# Source function #
###################
install.packages("scrm")
library(scrm)
########
#Script#
########

for(data in c(1,2,3,4)){
  if(data==1){
    L=10^7
    M=2
  }
  if(data==2){
    L=10^8
    M=2
    
  }
  if(data==3){
    L=10^9
    M=2
  }
  if(data==4){
    L=10^10
    M=2
    
  }
  nsim=10
  Pop=10^4
  mu=1.25*10^-8
  r=1*10^-8
  theta=mu*4*Pop*L
  rho=r*4*Pop*L
  shartp_v=c(2,5,10,50)
        for(sharp in c(1,2,3,4)){
          sharp_param=shartp_v[sharp]
              sum=0
              count=0
              mat_t=vector()
              for(i in seq(1,6,1)){
                if((i %% 2) == 0) {
                  time_e=(10^i)/(2*Pop)
                  time_e_1=((10^i)+0.5*(10^i))/(2*Pop)
                  v_t=c(time_e,log((1/sharp_param))/abs(time_e-time_e_1))
                  
                  mat_t=rbind(mat_t,v_t)
                  time_e=((10^i)+0.5*(10^i))/(2*Pop)
                  time_e_1=((10^(i+1)))/(2*Pop)
                  v_t=c(time_e,log(sharp_param)/abs(time_e-time_e_1))
                  mat_t=rbind(mat_t, v_t)
                } else {
                  time_e=(10^i)/(2*Pop)
                  time_e_1=((10^i)+0.5*(10^i))/(2*Pop)
                  v_t=c(time_e,log(sharp_param)/abs(time_e-time_e_1))
                  
                  mat_t=rbind(mat_t,v_t)
                  time_e=((10^i)+0.5*(10^i))/(2*Pop)
                  time_e_1=((10^(i+1)))/(2*Pop)
                  v_t=c(time_e,log((1/sharp_param))/abs(time_e-time_e_1))
                  mat_t=rbind(mat_t, v_t)
                }
              }
              v_t=c((10^(i+1))/(2*Pop),1)
              mat_t=rbind(mat_t, v_t)
            command_simu=paste(M,1,"-t",theta,"-r",rho,L,sep=" ")
            for(i in 1:(dim(mat_t)[1]-1)){
              command_simu=paste(command_simu,"-eG",mat_t[i,1],mat_t[i,2],sep=" ")
            }
            command_simu=paste(command_simu,"-eN",mat_t[dim(mat_t)[1],1],mat_t[dim(mat_t)[1],2],sep=" ")
            command_simu=paste(command_simu," -T -p 10 ",sep="")
            for(x in 1:nsim){
            # scrm(command_simu,file=paste("Figure_1",sharp,data,x,".txt",sep="")) #
            system(paste("scrm ",command_simu," > ",paste("Figure_1",sharp,data,x,".txt",sep=""))) # Assuming scrm is installed on your machine
            }
          
        }
      
}
```
