## Supplementary material for "Limits and Convergence properties of the Sequentially Markovian Coalescent": command lines for simulation: Figure_3.py

```
import matplotlib
import numpy as np
import pandas as pd
import scipy.special
import scipy.stats
from matplotlib import lines as mlines
from matplotlib import pyplot

import msprime
import msprime.cli as cli
sample_size=20
Ne=10**4
r=1*1e-8
m=1.25*1e-8
L=10 ** 8
population_configurations=[msprime.PopulationConfiguration(initial_size=Ne,sample_size=sample_size)]
for strength in [2]:
    demographic_events = [
                msprime.PopulationParametersChange(
                    time=20, growth_rate=2772.58872223978/(4*Ne)
                ),
                msprime.PopulationParametersChange(
                    time=30, growth_rate=-163.093454249399/(4*Ne)
                ),
                msprime.PopulationParametersChange(
                    time=200, growth_rate=-277.258872223978/(4*Ne)
                ),
                msprime.PopulationParametersChange(
                    time=300, growth_rate=16.3093454249399/(4*Ne)
                ),
                msprime.PopulationParametersChange(
                    time=2000, growth_rate=27.7258872223978/(4*Ne)
                ),
                msprime.PopulationParametersChange(
                    time=3000, growth_rate=-1.63093454249399/(4*Ne)
                ),
                msprime.PopulationParametersChange(
                    time=20000, growth_rate=-2.77258872223978/(4*Ne)
                ),
                msprime.PopulationParametersChange(
                    time=30000, growth_rate=0.163093454249399/(4*Ne)
                ),
                msprime.PopulationParametersChange(
                    time=200000, growth_rate=0.277258872223978/(4*Ne)
                ),
                msprime.PopulationParametersChange(
                    time=300000, growth_rate=-0.0163093454249399/(4*Ne)
                ),
                msprime.PopulationParametersChange(
                    time=2000000, growth_rate=-0.0277258872223978/(4*Ne)
                ),
                msprime.PopulationParametersChange(
                    time=3000000, growth_rate=0.00163093454249399/(4*Ne)
                ),
                msprime.PopulationParametersChange(
                    time=20000000, growth_rate=0, initial_size=Ne
                ),
            ]
    for x in range(1,11):
        ts=msprime.simulate(population_configurations=population_configurations,recombination_rate=r,mutation_rate=m,length=L,demographic_events=demographic_events)
        name="Figure_3"+str(strength)+"_"+str(x)+".vcf"
        with open(name, "w") as vcf_file:
            ts.write_vcf(vcf_file, 2)
        vcf_file.close()

for strength in [5]:
    demographic_events = [
                msprime.PopulationParametersChange(
                    time=20, growth_rate=6437.7516497364/(4*Ne)
                ),
                msprime.PopulationParametersChange(
                    time=30, growth_rate=-378.691273513906/(4*Ne)
                ),
                msprime.PopulationParametersChange(
                    time=200, growth_rate=-643.77516497364/(4*Ne)
                ),
                msprime.PopulationParametersChange(
                    time=300, growth_rate=37.8691273513906/(4*Ne)
                ),
                msprime.PopulationParametersChange(
                    time=2000, growth_rate=64.377516497364/(4*Ne)
                ),
                msprime.PopulationParametersChange(
                    time=3000, growth_rate=-3.78691273513906/(4*Ne)
                ),
                msprime.PopulationParametersChange(
                    time=20000, growth_rate=-6.4377516497364/(4*Ne)
                ),
                msprime.PopulationParametersChange(
                    time=30000, growth_rate=0.378691273513906/(4*Ne)
                ),
                msprime.PopulationParametersChange(
                    time=200000, growth_rate=0.64377516497364/(4*Ne)
                ),
                msprime.PopulationParametersChange(
                    time=300000, growth_rate=-0.0378691273513906/(4*Ne)
                ),
                msprime.PopulationParametersChange(
                    time=2000000, growth_rate=-0.064377516497364/(4*Ne)
                ),
                msprime.PopulationParametersChange(
                    time=3000000, growth_rate=0.00378691273513906/(4*Ne)
                ),
                msprime.PopulationParametersChange(
                    time=20000000, growth_rate=0, initial_size=Ne
                ),
            ]
    for x in range(1,11):
        ts=msprime.simulate(population_configurations=population_configurations,recombination_rate=r,mutation_rate=m,length=L,demographic_events=demographic_events)
        name="Figure_3"+str(strength)+"_"+str(x)+".vcf"
        with open(name, "w") as vcf_file:
            ts.write_vcf(vcf_file, 2)
        vcf_file.close()

for strength in [10]:
    demographic_events = [
                msprime.PopulationParametersChange(
                    time=20, growth_rate=9210.34037197618/(4*Ne)
                ),
                msprime.PopulationParametersChange(
                    time=30, growth_rate=-541.784727763305/(4*Ne)
                ),
                msprime.PopulationParametersChange(
                    time=200, growth_rate=-921.034037197618/(4*Ne)
                ),
                msprime.PopulationParametersChange(
                    time=300, growth_rate=54.1784727763305/(4*Ne)
                ),
                msprime.PopulationParametersChange(
                    time=2000, growth_rate=92.1034037197618/(4*Ne)
                ),
                msprime.PopulationParametersChange(
                    time=3000, growth_rate=-5.41784727763305/(4*Ne)
                ),
                msprime.PopulationParametersChange(
                    time=20000, growth_rate=-9.21034037197618/(4*Ne)
                ),
                msprime.PopulationParametersChange(
                    time=30000, growth_rate=0.541784727763305/(4*Ne)
                ),
                msprime.PopulationParametersChange(
                    time=200000, growth_rate=0.921034037197618/(4*Ne)
                ),
                msprime.PopulationParametersChange(
                    time=300000, growth_rate=-0.0541784727763305/(4*Ne)
                ),
                msprime.PopulationParametersChange(
                    time=2000000, growth_rate=-0.0921034037197618/(4*Ne)
                ),
                msprime.PopulationParametersChange(
                    time=3000000, growth_rate=0.00541784727763305/(4*Ne)
                ),
                msprime.PopulationParametersChange(
                    time=20000000, growth_rate=0, initial_size=Ne
                ),
            ]
    for x in range(1,11):
        ts=msprime.simulate(population_configurations=population_configurations,recombination_rate=r,mutation_rate=m,length=L,demographic_events=demographic_events)
        name="Figure_3"+str(strength)+"_"+str(x)+".vcf"
        with open(name, "w") as vcf_file:
            ts.write_vcf(vcf_file, 2)
        vcf_file.close()

for strength in [50]:
    demographic_events = [
                msprime.PopulationParametersChange(
                    time=20, growth_rate=15648.0920217126/(4*Ne)
                ),
                msprime.PopulationParametersChange(
                    time=30, growth_rate=-920.476001277211/(4*Ne)
                ),
                msprime.PopulationParametersChange(
                    time=200, growth_rate=-1564.80920217126/(4*Ne)
                ),
                msprime.PopulationParametersChange(
                    time=300, growth_rate=92.0476001277211/(4*Ne)
                ),
                msprime.PopulationParametersChange(
                    time=2000, growth_rate=156.480920217126/(4*Ne)
                ),
                msprime.PopulationParametersChange(
                    time=3000, growth_rate=-9.20476001277211/(4*Ne)
                ),
                msprime.PopulationParametersChange(
                    time=20000, growth_rate=-15.6480920217126/(4*Ne)
                ),
                msprime.PopulationParametersChange(
                    time=30000, growth_rate=0.920476001277211/(4*Ne)
                ),
                msprime.PopulationParametersChange(
                    time=200000, growth_rate=1.56480920217126/(4*Ne)
                ),
                msprime.PopulationParametersChange(
                    time=300000, growth_rate=-0.0920476001277211/(4*Ne)
                ),
                msprime.PopulationParametersChange(
                    time=2000000, growth_rate=-0.156480920217126/(4*Ne)
                ),
                msprime.PopulationParametersChange(
                    time=3000000, growth_rate=0.00920476001277211/(4*Ne)
                ),
                msprime.PopulationParametersChange(
                    time=20000000, growth_rate=0, initial_size=Ne
                ),
            ]
    for x in range(1,11):
        ts=msprime.simulate(population_configurations=population_configurations,recombination_rate=r,mutation_rate=m,length=L,demographic_events=demographic_events)
        name="Figure_3"+str(strength)+"_"+str(x)+".vcf"
        with open(name, "w") as vcf_file:
            ts.write_vcf(vcf_file, 2)
        vcf_file.close()
```
