## Supplementary material for "Limits and Convergence properties of the Sequentially Markovian Coalescent": command lines for simulation: Figure_4.R

```
###################
# Source function #
###################
install.packages("scrm")
library(scrm)
########
#Script#
########

for(M in c(2,4)){
for(r in c(0.1,1,10)){
  L=10^8
  nsim=10
  Pop=10^4
  mu=1.25*10^-8
  r=mu*r
  theta=mu*4*Pop*L
  rho=r*4*Pop*L
  shartp_v=c(2,5,10,50)
  for(sharp in c(3)){
    sharp_param=shartp_v[sharp]
    command_simu=paste(M,1,"-t",theta,"-r",rho,L,sep=" ")
    command_simu=paste(command_simu,"-eN",0.3,(1/sharp_param),sep=" ")
    command_simu=paste(command_simu,"-eN",2,1,sep=" ")
    command_simu=paste(command_simu,"-eN",mat_t[dim(mat_t)[1],1],mat_t[dim(mat_t)[1],2],sep=" ")
    command_simu=paste(command_simu," -T -p 10",sep="")
    print(command_simu)
    for(x in 1:nsim){
     # scrm(command_simu,file=paste("Figure_4",r/mu,M,x,".txt",sep=""))
      system(paste("scrm ",command_simu," > ",paste("Figure_4",r/mu,M,x,".txt",sep="")))
    }

  }
}
}
```
