## Supplementary material for "Limits and Convergence properties of the Sequentially Markovian Coalescent": command lines for simulation: Figure_5.R

```
###################
# Source function #
###################
library(scrm)
library(eSMC2)
########
#Script#
########

L=10^8 # Sequence length
M=4 # number of sequences
    nsim=10
    Pop=10^4
mu=1.25*10^-8
r=1.25*10^-8
theta=mu*L*4*Pop
rho=r*L*4*Pop

            
              sum=0
              count=0
              mat_t=vector()
              for(i in seq(1,6,1)){
                if((i %% 2) == 0) {
                  time_e=(10^i)/(2*Pop)
                  time_e_1=((10^i)+0.5*(10^i))/(2*Pop)
                  v_t=c(time_e,log(0.2)/abs(time_e-time_e_1))
                  
                  mat_t=rbind(mat_t,v_t)
                  time_e=((10^i)+0.5*(10^i))/(2*Pop)
                  time_e_1=((10^(i+1)))/(2*Pop)
                  v_t=c(time_e,log(5)/abs(time_e-time_e_1))
                  mat_t=rbind(mat_t, v_t)
                } else {
                  time_e=(10^i)/(2*Pop)
                  time_e_1=((10^i)+0.5*(10^i))/(2*Pop)
                  v_t=c(time_e,log(5)/abs(time_e-time_e_1))
                  
                  mat_t=rbind(mat_t,v_t)
                  time_e=((10^i)+0.5*(10^i))/(2*Pop)
                  time_e_1=((10^(i+1)))/(2*Pop)
                  v_t=c(time_e,log(0.2)/abs(time_e-time_e_1))
                  mat_t=rbind(mat_t, v_t)
                }
                
                
              }
              v_t=c((10^(i+1))/(2*Pop),1)
              mat_t=rbind(mat_t, v_t)
            setwd("~/Documents/testcpp/scrm_full")
            command_simu=paste(M,nsim,"-t",theta,"-r",rho,L,sep=" ")
            for(i in 1:(dim(mat_t)[1]-1)){
              command_simu=paste(command_simu,"-eG",mat_t[i,1],mat_t[i,2],sep=" ")  
            }
            command_simu=paste(command_simu,"-eN",mat_t[dim(mat_t)[1],1],mat_t[dim(mat_t)[1],2],sep=" ")  
            command_simu=paste(command_simu,"-p 8 ",sep=" ")  
            

#scrm(command_simu,"Figure_5.txt")
system(paste("scrm ",command_simu," > ","Figure_5.txt"))
O_total_before_trim=Get_sim_data("Figure_5.txt",L,M,nsim)
          for(problem in 1:3){
            count_p=0
            for(p in c(0,0.05,0.1,0.25)){
              count_p=count_p+1
              for(x in 1:nsim){
                Os_=as.matrix(O_total_before_trim[[x]])
                if(p>0){
                  nb_extra=floor(p*dim(Os_)[2])
                  
                  pos_rm=as.numeric(unique(round(runif(nb_extra,1,dim(Os_)[2]))))
                  if(problem!=1){
                    while (length(pos_rm)<nb_extra) {
                      pos_rm=as.numeric(unique(c(pos_rm,round(runif((nb_extra-length(pos_rm)),1,dim(Os_)[2])))))
                    }
                    for(ppos in  1:length(pos_rm)){
                      Os_[round(runif(1,0,length(which(Os_[c(1:M),pos_rm[ppos]]==1)))),pos_rm[ppos]]=0
                    }
                  }
                  if(problem!=2){
                    pos=round(runif(nb_extra,1,L))
                    pos=sort(pos)
                    for (ii in pos) {
                      if(!ii%in%Os_[dim(Os_)[1],]){
                        mut=rep(0,(dim(Os_)[1]-1))
                        mut[round(runif(1,1,(dim(Os_)[1]-1)))]<-1
                        mut=c(mut,ii)
                        if(ii<Os_[dim(Os_)[1],1]){
                          Os_=cbind(mut,Os_)
                        }else{
                          if(ii<Os_[dim(Os_)[1],dim(Os_)[2]]){
                            pm=min(which(Os_[dim(Os_)[1],]>ii))
                            Os_=cbind(Os_[,1:(pm-1)],mut,Os_[,(pm:(dim(Os_)[2]))])
                          }else{
                            Os_=cbind(Os_,mut)
                          }
                          
                        }
                        
                      }else{
                        Os_[round(runif(1,1,(dim(Os_)[1]-1))),which(Os_[dim(Os_)[1],]==ii)]<-1
                      }
                    }
                  }
                  
                  
                  
                }
                Os_=as.matrix(Os_)
                create_realinput_msmc2(Os_,paste("Figure_5",problem,x,p,sep=""),num=T)
              }
            }
            
          }
        
      
    
    
  
  
```
