## Supplementary material for "Limits and Convergence properties of the Sequentially Markovian Coalescent": command lines for simulation: Figure_6.R

```
###################
# Source function #
###################
library(scrm)
library(eSMC2)
################################
source(file="~/script/Script_papier2/source_esmc2.R")
########
#Script#
########

L=2*10^6
M=2
NC=20
    nsim=10
    Pop=10^4
    mu=1.25*10^-8
    r=10^-8
    theta=4*Pop*L*mu
    rho=4*Pop*L*r
          for(x in 1:nsim){
            change_r=rep(1,NC)
            change_mu=rep(1,NC)
            for(chr in 1:NC){
              mat=matrix(0)
              while(dim(mat)[2]<3){
                scale_r=change_r[chr]
                scale_mu=change_mu[chr]
                if(scale_r<0){
                  scale_r=abs(1/scale_r)
                }
                
                if(scale_mu<0){
                  scale_mu=abs(1/scale_mu)
                }
                command_simu=paste(M,1,"-t ",scale_mu*theta,"-r",scale_r*rho,L,command_simu_o,sep=" ")
                command_simu=paste(command_simu," -p 7 ",sep="")    
                # scrm(command_simu,paste("Figure_6",x,chr,".txt",sep=""))
                system(paste("scrm ",command_simu," > ",paste("Figure_6",x,chr,".txt",sep="")))
              }
            }

            change_r=rep(1,NC)
            change_mu=runif(NC,-5,5)
         
            for(chr in 1:NC){
              scale_r=change_r[chr]
              scale_mu=change_mu[chr]
              if(scale_r<0){
                scale_r=abs(1/scale_r)
              }
              
              if(scale_mu<0){
                scale_mu=abs(1/scale_mu)
              }
              command_simu=paste(M,1,"-t ",scale_mu*theta,"-r",scale_r*rho,L,command_simu_o,sep=" ")
              command_simu=paste(command_simu," -p 7 ",sep="")    
              #scrm(command_simu,paste("Figure_6_rec_issue",x,chr,".txt",sep=""))
              system(paste("scrm ",command_simu," > ",paste("Figure_6_rec_issue",x,chr,".txt",sep="")))
            }
            
           
            
            change_r=runif(NC,-5,5)
            change_mu=rep(1,NC)
           
            for(chr in 1:NC){
              scale_r=change_r[chr]
              scale_mu=change_mu[chr]
              if(scale_r<0){
                scale_r=abs(1/scale_r)
              }
              
              if(scale_mu<0){
                scale_mu=abs(1/scale_mu)
              }
              command_simu=paste(M,1,"-t ",scale_mu*theta,"-r",scale_r*rho,L,command_simu_o,sep=" ")
              command_simu=paste(command_simu," -p 7 ",sep="")    

              #scrm(command_simu,paste("Figure_6_mut_issue",x,chr,".txt",sep=""))
              system(paste("scrm ",command_simu," > ",paste("Figure_6_mut_issue",x,chr,".txt",sep="")))
            }
            
            
   
            
            change_r=runif(NC,-5,5)
            change_mu=runif(NC,-5,5)

            for(chr in 1:NC){
              scale_r=change_r[chr]
              scale_mu=change_mu[chr]
              if(scale_r<0){
                scale_r=abs(1/scale_r)
              }
              
              if(scale_mu<0){
                scale_mu=abs(1/scale_mu)
              }
              command_simu=paste(M,1,"-t ",scale_mu*theta,"-r",scale_r*rho,L,command_simu_o,sep=" ")
              command_simu=paste(command_simu," -p 7 ",sep="")    
              #scrm(command_simu,paste("Figure_6_mut_and_rec_issue",x,chr,".txt",sep=""))
              system(paste("scrm ",command_simu," > ",paste("Figure_6_mut_and_rec_issue",x,chr,".txt",sep="")))
            }
            
          }

    
  
```
