## Supplementary material for "Limits and Convergence properties of the Sequentially Markovian Coalescent": command lines for simulation: Figure_7.R

```
###################
# Source function #
###################
library(scrm)
library(eSMC2)
########
#Script#
########
l=20*10^7
M=4
nsim=10
Pop=10^4
mu=1.25*10^-8
r=1.25*10^-8
theta=4*Pop*L*mu
rho=4*Pop*L*r

              sum=0
              count=0
              mat_t=vector()
              for(i in seq(1,6,1)){
                if((i %% 2) == 0) {
                  time_e=(10^i)/(2*Pop)
                  time_e_1=((10^i)+0.5*(10^i))/(2*Pop)
                  v_t=c(time_e,log(0.2)/abs(time_e-time_e_1))
                  
                  mat_t=rbind(mat_t,v_t)
                  time_e=((10^i)+0.5*(10^i))/(2*Pop)
                  time_e_1=((10^(i+1)))/(2*Pop)
                  v_t=c(time_e,log(5)/abs(time_e-time_e_1))
                  mat_t=rbind(mat_t, v_t)
                } else {
                  time_e=(10^i)/(2*Pop)
                  time_e_1=((10^i)+0.5*(10^i))/(2*Pop)
                  v_t=c(time_e,log(5)/abs(time_e-time_e_1))
                  
                  mat_t=rbind(mat_t,v_t)
                  time_e=((10^i)+0.5*(10^i))/(2*Pop)
                  time_e_1=((10^(i+1)))/(2*Pop)
                  v_t=c(time_e,log(0.2)/abs(time_e-time_e_1))
                  mat_t=rbind(mat_t, v_t)
                }
                
                
              }
              v_t=c((10^(i+1))/(2*Pop),1)
              mat_t=rbind(mat_t, v_t)
            setwd("~/Documents/testcpp/scrm_full")
            command_simu=paste(M,nsim,"-t",theta,"-r",rho,L,sep=" ")
            for(i in 1:(dim(mat_t)[1]-1)){
              command_simu=paste(command_simu,"-eG",mat_t[i,1],mat_t[i,2],sep=" ")  
            }
            command_simu=paste(command_simu,"-eN",mat_t[dim(mat_t)[1],1],mat_t[dim(mat_t)[1],2],sep=" ")  
            command_simu=paste(command_simu,"-p 8 ",sep=" ")  

scrm(command_simu,"Figure_7.txt")

          data_before_trim=Get_sim_data("Figure_7.txt",L,M,nsim)
          
      
          for(p in c(0.05,0.1,0.25,0.50)){

            for(kb in c(1000)){

              
              for(trim in 1:4){
                
                if(trim==1){
                  data=data_before_trim
                }
                if(trim==2){
                  data=list()
                  for(x in 1:nsim){
                    length_cut=vector()
                    while ((length(length_cut)*kb/L)<p){
                      pos_t=unique(floor(runif((p-(length(length_cut)*kb/L))*(2*L/kb),0,(L/kb)))) 
                      length_cut=unique(c(length_cut,pos_t))
                    }
                    if((length(length_cut)*kb/L)>p){
                      length_cut=sort(length_cut[1:ceiling(p/(kb/L))])
                    }
                    pos_r=vector()
                    Seg_mat=data_before_trim[[x]]
                    for(xx in length_cut ){
                      yy=(xx+1)*kb
                      xx=xx*kb
                      pos_r=c(pos_r,which(Seg_mat[(M+1),]>=xx&Seg_mat[(M+1),]<=yy))
                    }
                    pos_r=sort(unique(pos_r))
                    Seg_mat=Seg_mat[,-pos_r]
                    data[[x]]=Seg_mat
                  }
                }
                if(trim==3){
                  data=list()
                  for(x in 1:nsim){
                    length_cut=vector()
                    while ((length(length_cut)*kb/L)<p){
                      pos_t=unique(floor(runif((p-(length(length_cut)*kb/L))*(2*L/kb),0,(L/kb))))
                      length_cut=unique(c(length_cut,pos_t))
                    }
                    if((length(length_cut)*kb/L)>p){
                      length_cut=sort(length_cut[1:ceiling(p/(kb/L))])
                    }
                    length_cut=sort(length_cut,decreasing =T)
                    Seg_mat=matrix(0,nrow = (M+1),ncol = L)
                    Seg_mat[(1:M),data_before_trim[[x]][(M+1),]]=data_before_trim[[x]][(1:M),]
                    for(xx in length_cut ){
                      yy=(xx+1)*kb
                      xx=xx*kb
                      Seg_mat=Seg_mat[,-c(xx:yy)]
                    }
                    pos=which(colSums(Seg_mat)>0)
                    Seg_mat=Seg_mat[,pos]
                    Seg_mat[(M+1),]=pos
                    data[[x]]=Seg_mat
                  }
                }
                if(trim==4){
                  data=list()
                  for(x in 1:nsim){
                    length_cut=vector()
                    while ((length(length_cut)*kb/L)<(p*0.5)){
                      pos_t=unique(floor(runif(((p*0.5)-(length(length_cut)*kb/L))*(2*L/kb),0,(L/kb))))
                      length_cut=unique(c(length_cut,pos_t))
                    }
                    if((length(length_cut)*kb/L)>(p*0.5)){
                      length_cut=sort(length_cut[1:ceiling((p*0.5)/(kb/L))])
                    }
                    pos_r=vector()
                    Seg_mat=data_before_trim[[x]]
                    for(xx in length_cut ){
                      yy=(xx+1)*kb
                      xx=xx*kb
                      pos_r=c(pos_r,which(Seg_mat[(M+1),]>=xx&Seg_mat[(M+1),]<=yy))
                    }
                    pos_r=sort(unique(pos_r))
                    Seg_mat_i=Seg_mat[,-pos_r]
                    rm(Seg_mat)
                    length_cut=vector()
                    while ((length(length_cut)*kb/L)<(p*0.5)){
                      pos_t=unique(sort(floor(runif(((p*0.5)-(length(length_cut)*kb/L))*(2*L/kb),0,(L/kb))))) 
                      length_cut=sort(unique(c(length_cut,pos_t)))
                    }
                    if((length(length_cut)*kb/L)>(p*0.5)){
                      length_cut=sort(length_cut[1:ceiling((p*0.5)/(kb/L))])
                    }
                    length_cut=sort(length_cut,decreasing =T)
                    Seg_mat=matrix(0,nrow = (M+1),ncol = L)
                    Seg_mat[(1:M),Seg_mat_i[(M+1),]]=Seg_mat_i[(1:M),]
                    rm(Seg_mat_i)
                    for(xx in length_cut ){
                      yy=(xx+1)*kb
                      xx=xx*kb
                      Seg_mat=Seg_mat[,-c(xx:yy)]
                    }
                    pos=which(colSums(Seg_mat)>0)
                    Seg_mat=Seg_mat[,pos]
                    Seg_mat[(M+1),]=pos
                    data[[x]]=Seg_mat
                  }
                }

                for(x in 1:nsim){
                  Os_=as.matrix(data[[x]])
                  create_realinput_msmc2(Os_,paste("Figure_7",x,p,kb,trim,".txt",sep=""),num=T)
                }

              }
            }
          }

        
      

  
```
