## Supplementary material for "Limits and Convergence properties of the Sequentially Markovian Coalescent": command lines for simulation: S4_fig.R

```
###################
# Source function #
###################
install.packages("scrm")
library(scrm)
########
#Script#
########
L=10^10
M=2
nsim=10
Pop=10^4
mu=1.25*10^-8
r=1*10^-8
theta=mu*4*Pop*L
rho=r*4*Pop*L
for(scenario in c(1,2,3,4)){
shartp_v=c(2,5,10,50)
for(sharp in c(4)){
sharp_param=shartp_v[sharp]
      if(scenario==1){
          sum=0
          count=0
          mat_t=vector()
          for(i in seq(1,6,1)){
            if((i %% 2) == 0) {
              time_e=(10^i)/(2*Pop)
              time_e_1=((10^i)+0.5*(10^i))/(2*Pop)
              v_t=c(time_e,log((1/sharp_param))/abs(time_e-time_e_1))
              
              mat_t=rbind(mat_t,v_t)
              time_e=((10^i)+0.5*(10^i))/(2*Pop)
              time_e_1=((10^(i+1)))/(2*Pop)
              v_t=c(time_e,log(sharp_param)/abs(time_e-time_e_1))
              mat_t=rbind(mat_t, v_t)
            } else {
              time_e=(10^i)/(2*Pop)
              time_e_1=((10^i)+0.5*(10^i))/(2*Pop)
              v_t=c(time_e,log(sharp_param)/abs(time_e-time_e_1))
              
              mat_t=rbind(mat_t,v_t)
              time_e=((10^i)+0.5*(10^i))/(2*Pop)
              time_e_1=((10^(i+1)))/(2*Pop)
              v_t=c(time_e,log((1/sharp_param))/abs(time_e-time_e_1))
              mat_t=rbind(mat_t, v_t)
            }
          }
          v_t=c((10^(i+1))/(2*Pop),1)
          mat_t=rbind(mat_t, v_t)
        
        
        command_simu=paste(M,1,"-t",theta,"-r",rho,L,sep=" ")
        for(i in 1:(dim(mat_t)[1]-1)){
          command_simu=paste(command_simu,"-eG",mat_t[i,1],mat_t[i,2],sep=" ")
        }
        command_simu=paste(command_simu,"-eN",mat_t[dim(mat_t)[1],1],mat_t[dim(mat_t)[1],2],sep=" ")
        
      }
      if(scenario==2){
        command_simu=paste(M,1,"-t",theta,"-r",rho,L,sep=" ")
        command_simu=paste(command_simu,"-eN",0.3,(1/sharp_param),sep=" ")
        command_simu=paste(command_simu,"-eN",2,1,sep=" ")
      }
      if(scenario==3){
        sum=0
        count=0
        mat_t=vector()
        for(i in seq(3,3,1)){
          if((i %% 2) == 0) {
            time_e=(10^i)/(2*Pop)
            time_e_1=10^(i+1)/(2*Pop)
            v_t=c(time_e,log((1/sharp_param))/abs(time_e-time_e_1))
            mat_t=rbind(mat_t,v_t)
          } else {
            time_e=(10^i)/(2*Pop)
            time_e_1=10^(i+1)/(2*Pop)
            v_t=c(time_e,log(sharp_param)/abs(time_e-time_e_1))
            mat_t=rbind(mat_t,v_t)
          }
        }
        v_t=c((10^(i+1))/(2*Pop),(1/sharp_param))
        mat_t=rbind(mat_t, v_t)
        command_simu=paste(M,1,"-t",theta,"-r",rho,L,sep=" ")
        for(i in 1:(dim(mat_t)[1]-1)){
          command_simu=paste(command_simu,"-eG",mat_t[i,1],mat_t[i,2],sep=" ")
        }
        command_simu=paste(command_simu,"-eN",mat_t[dim(mat_t)[1],1],mat_t[dim(mat_t)[1],2],sep=" ")
        
      }
      if(scenario==4){
        sum=0
        count=0
        mat_t=vector()
        for(i in seq(3,3,1)){
          if((i %% 2) == 0) {
            time_e=(10^i)/(2*Pop)
            time_e_1=10^(i+1)/(2*Pop)
            v_t=c(time_e,log(sharp_param)/abs(time_e-time_e_1))
            mat_t=rbind(mat_t,v_t)
          } else {
            time_e=(10^i)/(2*Pop)
            time_e_1=10^(i+1)/(2*Pop)
            v_t=c(time_e,log((1/sharp_param))/abs(time_e-time_e_1))
            mat_t=rbind(mat_t,v_t)
          }
        }
        v_t=c((10^(i+1))/(2*Pop),sharp_param)
        command_simu=paste(M,1,"-t",theta,"-r",rho,L,sep=" ")
        for(i in 1:(dim(mat_t)[1]-1)){
          command_simu=paste(command_simu,"-eG",mat_t[i,1],mat_t[i,2],sep=" ")
        }
        command_simu=paste(command_simu,"-eN",mat_t[dim(mat_t)[1],1],mat_t[dim(mat_t)[1],2],sep=" ")
        
      }
       command_simu=paste(command_simu," -T -p 10",sep="")
       print(command_simu)
       for(x in 1:nsim){
        # scrm(command_simu,file=paste("S4_fig",sharp,scenario,x,".txt",sep=""))
         system(paste("scrm ",command_simu," > ",paste("S4_fig",sharp,scenario,x,".txt",sep="")))
       }

    }

  }

  
  
```
