## Supplementary material for "Limits and Convergence properties of the Sequentially Markovian Coalescent": command lines for simulation: S6_fig.R

```
###################
# Source function #
###################
install.packages("scrm")
library(scrm)
########
#Script#
########

for(data in c(1,2,3,4)){
  if(data==1){
    L=10^7
    M=4
  }
  if(data==2){
    L=10^8
    M=4
    
  }
  if(data==3){
    L=10^9
    M=4
  }
  if(data==4){
    L=10^10
    M=4
    
  }
  nsim=10
  Pop=10^4
  mu=1.25*10^-8
  r=1*10^-8
  theta=mu*4*Pop*L
  rho=r*4*Pop*L
  
    shartp_v=c(2,5,10,50)
    for(sharp in c(1,2,3,4)){
      sharp_param=shartp_v[sharp]
      command_simu=paste(M,1,"-t",theta,"-r",rho,L,sep=" ")
      command_simu=paste(command_simu,"-eN",0.3,(1/sharp_param),sep=" ")
      command_simu=paste(command_simu,"-eN",2,1,sep=" ")
      command_simu=paste(command_simu,"-eN",mat_t[dim(mat_t)[1],1],mat_t[dim(mat_t)[1],2],sep=" ")
      command_simu=paste(command_simu," -T -p 10",sep="")
      print(command_simu)
      for(x in 1:nsim){
       # scrm(command_simu,file=paste("S6_fig",sharp,data,x,".txt",sep=""))
        system(paste("scrm ",command_simu," > ",paste("S6_fig",sharp,data,x,".txt",sep="")))
      }

    }
  
}
```
