## Supplementary material for "Limits and Convergence properties of the Sequentially Markovian Coalescent": command lines for simulation: S11_fig.R

```
###################
# Source function #
###################
install.packages("scrm")
library(scrm)
########
#Script#
########

for(M in c(2,4)){
  L=10^8
  nsim=10
  Pop=10^4
  mu=1.25*10^-8
  r=1*10^-8
  theta=mu*4*Pop*L
  rho=r*4*Pop*L
    command_simu=paste(M,1,"-t",theta,"-r",rho,L,sep=" ")
    command_simu=paste(command_simu," -T -p 10",sep="")
    print(command_simu)
    for(x in 1:nsim){
      #scrm(command_simu,file=paste("S11_fig",sharp,M,x,".txt",sep="")) 
      system(paste("scrm ",command_simu," > ",paste("S11_fig",M,x,".txt",sep="")))
    }
  
}
```
