## Supplementary material for "Limits and Convergence properties of the Sequentially Markovian Coalescent": command lines for simulation: S14_fig.py

```
import matplotlib
import numpy as np
import pandas as pd
import scipy.special
import scipy.stats
from matplotlib import lines as mlines
from matplotlib import pyplot

import msprime
import msprime.cli as cli
sample_size=20
Ne=10**4
r=1*1e-8
m=1.25*1e-8
L=10 ** 8
population_configurations=[msprime.PopulationConfiguration(initial_size=Ne,sample_size=sample_size)]
for strengh in [2]:
    demographic_events = [
                msprime.PopulationParametersChange(
                    time=2000, growth_rate=7.70163533955495*1e-5/2
                ),
                msprime.PopulationParametersChange(
                    time=20000, growth_rate=0,initial_size=Ne/strengh
                ),
            ]
    for x in range(1,11):
        ts=msprime.simulate(population_configurations=population_configurations,recombination_rate=r,mutation_rate=m,length=L,demographic_events=demographic_events)
        name="S12_fig_Expansion_strength_"+str(strengh)+"_"+str(x)+".vcf"
        with open(name, "w") as vcf_file:
            ts.write_vcf(vcf_file, 2)
        vcf_file.close()

for strengh in [5]:
    demographic_events = [
                msprime.PopulationParametersChange(
                    time=2000, growth_rate=0.0001788264347149/2
                ),
                msprime.PopulationParametersChange(
                    time=20000, growth_rate=0,initial_size=Ne/strengh
                ),
            ]
    for x in range(1,11):
        ts=msprime.simulate(population_configurations=population_configurations,recombination_rate=r,mutation_rate=m,length=L,demographic_events=demographic_events)
        name="S12_fig_Expansion_strength_"+str(strengh)+"_"+str(x)+".vcf"
        with open(name, "w") as vcf_file:
            ts.write_vcf(vcf_file, 2)
        vcf_file.close()

for strengh in [10]:
    demographic_events = [
                msprime.PopulationParametersChange(
                    time=2000, growth_rate=0.00025584278811045/2
                ),
                msprime.PopulationParametersChange(
                    time=20000, growth_rate=0,initial_size=Ne/strengh
                ),
            ]
    for x in range(1,11):
        ts=msprime.simulate(population_configurations=population_configurations,recombination_rate=r,mutation_rate=m,length=L,demographic_events=demographic_events)
        name="S12_fig_Expansion_strength_"+str(strengh)+"_"+str(x)+".vcf"
        with open(name, "w") as vcf_file:
            ts.write_vcf(vcf_file, 2)
        vcf_file.close()

for strengh in [50]:
    demographic_events = [
                msprime.PopulationParametersChange(
                    time=2000, growth_rate=0.00043466922282535/2
                ),
                msprime.PopulationParametersChange(
                    time=20000, growth_rate=0,initial_size=Ne/strengh
                ),
            ]
    for x in range(1,11):
        ts=msprime.simulate(population_configurations=population_configurations,recombination_rate=r,mutation_rate=m,length=L,demographic_events=demographic_events)
        name="S12_fig_Expansion_strength_"+str(strengh)+"_"+str(x)+".vcf"
        with open(name, "w") as vcf_file:
            ts.write_vcf(vcf_file, 2)
        vcf_file.close()
```
