## Supplementary material for "Limits and Convergence properties of the Sequentially Markovian Coalescent": command lines for simulation: S16_fig.R

```
###################
# Source function #
###################
install.packages("scrm")
library(scrm)
########
#Script#
########
M=4
L=50*10^8
nsim=10
Pop=10^4
mu=1.25*10^-8
r=10^-8
theta=4*Pop*mu*L
rho=4*Pop*r*L
        for(scenario in c(1,2)){

          if(scenario==1){
            
              sum=0
              count=0
              mat_t=vector()
              for(i in seq(1,6,1)){
                if((i %% 2) == 0) {
                  time_e=(10^i)/(2*Pop)
                  time_e_1=((10^i)+0.5*(10^i))/(2*Pop)
                  v_t=c(time_e,log(0.2)/abs(time_e-time_e_1))
                  
                  mat_t=rbind(mat_t,v_t)
                  time_e=((10^i)+0.5*(10^i))/(2*Pop)
                  time_e_1=((10^(i+1)))/(2*Pop)
                  v_t=c(time_e,log(5)/abs(time_e-time_e_1))
                  mat_t=rbind(mat_t, v_t)
                } else {
                  time_e=(10^i)/(2*Pop)
                  time_e_1=((10^i)+0.5*(10^i))/(2*Pop)
                  v_t=c(time_e,log(5)/abs(time_e-time_e_1))
                  
                  mat_t=rbind(mat_t,v_t)
                  time_e=((10^i)+0.5*(10^i))/(2*Pop)
                  time_e_1=((10^(i+1)))/(2*Pop)
                  v_t=c(time_e,log(0.2)/abs(time_e-time_e_1))
                  mat_t=rbind(mat_t, v_t)
                }
                
                
              }
              v_t=c((10^(i+1))/(2*Pop),1)
              mat_t=rbind(mat_t, v_t)
            
            setwd("~/Documents/testcpp/scrm_full")
            command_simu=paste(M,nsim,"-t",theta,"-r",rho,L,sep=" ")
            for(i in 1:(dim(mat_t)[1]-1)){
              command_simu=paste(command_simu,"-eG",mat_t[i,1],mat_t[i,2],sep=" ")  
            }
            command_simu=paste(command_simu,"-eN",mat_t[dim(mat_t)[1],1],mat_t[dim(mat_t)[1],2],sep=" ")  
            command_simu=paste(command_simu,"-p",ceiling(log10(L)),sep=" ")  
            
            
          }
          if(scenario==2){
            
            Pop_size=rep(Pop,length(x_ab))
            command_simu=paste(M,nsim,"-t",theta,"-r",rho,L,sep=" ")
            command_simu=paste(command_simu,"-p",ceiling(log10(L)),sep=" ")
            
          }
          for(x in 1:nsim){
            scrm(command_simu,paste("S14_fig",scenario,x,".txt",sep=""))
          }


        }

    
    
  
  
```
