## Supplementary material for "Limits and Convergence properties of the Sequentially Markovian Coalescent": command lines for simulation: S22_fig.R

```
###################
# Source function #
###################
install.packages("scrm")
library(scrm)
########
#Script#
########

for(M in c(2,4)){
  for(r in c(10)){
    L=10^8
    nsim=10
    Pop=10^4
    mu=1.25*10^-8
    r=mu*r
    theta=mu*4*Pop*L
    rho=r*4*Pop*L
    shartp_v=c(2,5,10,50)
    for(sharp in c(3)){
      sharp_param=shartp_v[sharp]
      command_simu=paste(M,nsim,"-t",theta,"-r",rho,L,sep=" ")
      command_simu=paste(command_simu,"-eN",0.3,(1/sharp_param),sep=" ")
      command_simu=paste(command_simu,"-eN",2,1,sep=" ")
      command_simu=paste(command_simu,"-eN",mat_t[dim(mat_t)[1],1],mat_t[dim(mat_t)[1],2],sep=" ")
      command_simu=paste(command_simu," -p 10",sep="")
      print(command_simu)
      for(x in 1:nsim){
        #scrm(command_simu,file=paste("S22_fig",r/mu,M,x,".txt",sep=""))  
        system(paste("scrm ",command_simu," > ",paste("S22_fig",r/mu,M,x,".txt",sep=""))
      }
    }
  }
}
```
