## Supplementary material for "Limits and Convergence properties of the Sequentially Markovian Coalescent": command lines for simulation: S25_fig.R

```
###################
# Source function #
###################
library(eSMC2)
########
#Script#
########
results_esmc=list()
results_msmc=list()
results_msmc2=list()
results_smcpp=list()
mu_list=list()
r_list=list()
L=10^8
nsim=10
Pop=10^4
mu=1.25*10^-8
r=1*10^-8
theta=mu*4*Pop*L
rho=r/mu
pos_sim=list()
M=100
system("python3 S25_fig.py")
if(T){
for(x in 1:nsim){
  path=paste("S25_fig_",x,".vcf",sep="")

  for(MAF in c(0,0.01,0.05,0.1)){
    O_total=Process_vcf_data(path)
    if(MAF==0){
     create_realinput_msmc2(O_total,paste("S25_fig",MAF,x,sep=""),num=T)
      data=Get_vcf_data(path)
      data_written=matrix(NA,nrow =length(data),ncol = 19 )
       for(cc in 1:length(data)){
         truc=data[[cc]][1:min(19,length(data[[cc]]))]
         data_written[cc,1:length(truc)]=truc
       }      
      write.table( data_written,paste("S25_fig",MAF,x,".vcf",sep=""),na = "",append = F, quote = FALSE, row.names=FALSE, col.names=FALSE,sep="\t")
    }else{
      pos=c()
      for(cc in 1:dim(O_total)[2]){
        if(sum(as.numeric(O_total[1:M,cc]))<=(MAF*M)|sum(as.numeric(O_total[1:M,cc]))>=((1-MAF)*M)){
          pos=c(pos,cc)
        }
      }
      create_realinput_msmc2(O_total[,-c(pos)],paste("S25_fig",MAF,x,sep=""),num=T)

      data=Get_vcf_data(path)
      pos=pos+6
      data=data[-c(pos)]
      data_written=matrix(NA,nrow =length(data),ncol = 19 )
      for(cc in 1:length(data)){
        truc=data[[cc]][1:min(19,length(data[[cc]]))]
        data_written[cc,1:length(truc)]=truc
      }      
      write.table( data_written,paste("S25_fig",MAF,x,".vcf",sep=""), na = "", quote = FALSE, row.names=FALSE, col.names=FALSE,sep="\t")
    }
    rm(O_total)
  }
}

}
```
