## Supplementary material for "Limits and Convergence properties of the Sequentially Markovian Coalescent": command lines for simulation: S26_fig.R

```
###################
# Source function #
###################
library(scrm)
library(eSMC2)
################################
########
#Script#
########

L=40*10^6
M=2
mu=1.25*10^-8
r=1.25*10^-8
nsim=10
Pop=10^4
rho=4*Pop*L*r
theta=4*Pop*L*mu
command_simu=paste(M,1,"-r",rho,L,sep=" ")
command_simu_o=command_simu
          for(x in 1:nsim){
            command_simu=paste(command_simu_o," -t ",theta," -p 8 ",sep="")    
            scrm(command_simu,paste("S21_fig",x,".txt",sep=""))
            n_r=floor(L/(2*10^6))
            change_r=runif(n_r,-5,5)
            vect_r=" "
            count_rr=0
            for(scale_r in change_r){
              count_rr=count_rr+1
              if(scale_r<0){
                scale_r=abs(1/scale_r)
              }
              vect_r=paste(vect_r,"-sr",(count_rr-1)*(2*10^6),rho*scale_r,sep = " " )
            }
            command_simu=paste(command_simu_o," -t ",theta,vect_r," -p 8 ",sep="")    
            scrm(command_simu,paste("S26_fig_rec_issue",x,".txt",sep=""))
          }
  
```
