## Supplementary material for "Limits and Convergence properties of the Sequentially Markovian Coalescent": command lines for simulation: S35_fig.R

```
###################
# Source function #
###################
library(eSMC2)
library(scrm)
########
#Script#
########
L=20*10^7
M=4
nsim=10
Pop=10^4
mu=1.25*10^-8
r=1.25*10^-8
theta=4*Pop*L*mu
rho=4*Pop*L*r

sum=0
count=0
mat_t=vector()
for(i in seq(1,6,1)){
  if((i %% 2) == 0) {
    time_e=(10^i)/(2*Pop)
    time_e_1=((10^i)+0.5*(10^i))/(2*Pop)
    v_t=c(time_e,log(0.2)/abs(time_e-time_e_1))
    
    mat_t=rbind(mat_t,v_t)
    time_e=((10^i)+0.5*(10^i))/(2*Pop)
    time_e_1=((10^(i+1)))/(2*Pop)
    v_t=c(time_e,log(5)/abs(time_e-time_e_1))
    mat_t=rbind(mat_t, v_t)
  } else {
    time_e=(10^i)/(2*Pop)
    time_e_1=((10^i)+0.5*(10^i))/(2*Pop)
    v_t=c(time_e,log(5)/abs(time_e-time_e_1))
    
    mat_t=rbind(mat_t,v_t)
    time_e=((10^i)+0.5*(10^i))/(2*Pop)
    time_e_1=((10^(i+1)))/(2*Pop)
    v_t=c(time_e,log(0.2)/abs(time_e-time_e_1))
    mat_t=rbind(mat_t, v_t)
  }
  
  
}
v_t=c((10^(i+1))/(2*Pop),1)
mat_t=rbind(mat_t, v_t)
command_simu=paste("scrm",M,nsim,"-t",theta,"-r",rho,L,sep=" ")
for(i in 1:(dim(mat_t)[1]-1)){
  command_simu=paste(command_simu,"-eG",mat_t[i,1],mat_t[i,2],sep=" ")  
}
command_simu=paste(command_simu,"-eN",mat_t[dim(mat_t)[1],1],mat_t[dim(mat_t)[1],2],sep=" ")  
command_simu=paste(command_simu,"-p 8 > S35_fig.txt ",sep=" ")  

system(command_simu)

data_before_trim=Get_sim_data("S35_fig.txt",L,M,nsim)


for(p in c(0.05,0.1,0.25,0.50)){
  
  for(kb in c(1000,10000,100000)){


        data=list()
        for(x in 1:nsim){
          length_cut=vector()
          while ((length(length_cut)*kb/L)<p){
            pos_t=unique(floor(runif((p-(length(length_cut)*kb/L))*(2*L/kb),0,(L/kb)))) 
            length_cut=unique(c(length_cut,pos_t))
          }
          if((length(length_cut)*kb/L)>p){
            length_cut=sort(length_cut[1:ceiling(p/(kb/L))])
          }
          pos_r=vector()
          Seg_mat=data_before_trim[[x]]
          for(xx in length_cut ){
            yy=(xx+1)*kb
            xx=xx*kb
            pos_r=c(pos_r,which(Seg_mat[(M+1),]>=xx&Seg_mat[(M+1),]<=yy))

          }
          pos_r=sort(unique(pos_r))
          Seg_mat=Seg_mat[,-pos_r]
          data[[x]]=Seg_mat
          Os_=as.matrix(data[[x]])
          length_cut=length_cut[which((length_cut*kb)<Seg_mat[(M+1),dim(Seg_mat)[2]])]
          mask=list()
          for(pp in 1:length(length_cut)){
            mask[[pp]]=c((length_cut[pp]*kb),(length_cut[pp]+1)*kb)
          }
          create_realinput_msmc2_masked(Os_,paste("S35_fig",x,p,kb,"mask",sep=""),num=T,mask=mask)

      }
    
  }
}
```
