## Supplementary material for "Limits and Convergence properties of the Sequentially Markovian Coalescent": command lines for analyses: Figure_1.R

```
###################
# Source function #
###################

library(eSMC2)
########
#Script#
########
results=list()
for(data in c(1,2,3,4)){
  if(data==1){
    L=10^7
    M=2
  }
  if(data==2){
    L=10^8
    M=2
    
  }
  if(data==3){
    L=10^9
    M=2
  }
  if(data==4){
    L=10^10
    M=2
    
  }
  nsim=10
  Pop=10^4
  mu=1.25*10^-8
  r=1*10^-8
  theta=mu*4*Pop*L
        for(sharp in c(1,2,3,4)){
          for(x in 1:nsim){
            results[[(1+length(results))]]=Optimize_N(file=paste("Figure_1",sharp,data,x,".txt",sep=""),theta = theta,Rho=theta,L=L,n=40,ER=T,Pop=T,SB=F,SF=F,BoxP=c(3,3),Boxr=c(1,1),pop_vect=rep(2,20),Big_Window=F,Correct_window =T,mut=T)

          }

        }

      

}
```
