## Supplementary material for "Limits and Convergence properties of the Sequentially Markovian Coalescent": command lines for analyses: Figure_2.R

```
###################
# Source function #
###################
library('plot.matrix')
library(eSMC2)
########
#Script#
########
if(T){
  nsim=10
  theta=500*10^4
  rho=0.8
  L=10000*10^6
  Sharp=c(2,5,10,50)
    #eSMC
    
    pdf("Figure_2.pdf")
    par(mfrow=c(2,2))
    for(sharp in 1:4){
        N=matrix(nrow = 40*40,ncol = nsim)
        for(x in 1:nsim){
          BW_mat=get_Mat(file=paste("Figure_2",sharp,x,".txt",sep=""),theta=theta,Rho=theta,L=L,n=40,BoxB=c(0.1,1),BoxP=c(3,3),Boxr=c(1,1),Boxs=c(0,0.97),pop_vect=rep(2,20),window_scaling=c(1,0),Big_Window=0,mut=T,Correct_window = T)
          N[,x]=as.numeric(BW_mat$N)
          N[which(is.na(as.numeric(N[,x]))),x]<-0
        }
        N_heat=matrix(c(apply(N,1,sd)/apply(N,1,mean)),nrow = 40,ncol = 40)
        plot(N_heat,main=paste( letters[data],") Amplitude : ",Sharp[sharp],sep=""))
    }
    dev.off()
}
```
