## Supplementary material for "Limits and Convergence properties of the Sequentially Markovian Coalescent": command lines for analyses: Figure_3.R

```
###################
# Source function #
###################

library(eSMC2)
########
#Script#
########
results_esmc=list()
L=10^8
nsim=10
Pop=10^4
mu=1.25*10^-8
r=1*10^-8
theta=mu*4*Pop*L
strentgh=c(2,5,10,50)
for(sharp in c(1,2,3,4)){
for(method in c(1,3,4)){
  if(method<4){
  if(method==1){
    M=2
    L=10^8
    for(x in 1:nsim){
      path=paste("Figure_3",sharp,M,x,".txt",sep="")
      O_total=Get_sim_data(path,L,M,1)
      # esmc
      results_esmc[[(1+length(results_esmc))]]=eSMC2(n=40,rho=1,O_total[[1]],mu_r=mu,maxit =20,symbol_number=25,BoxB=c(0.05,1),BoxP=c(3,3),Boxr=c(1,1),Boxs=c(0,0.97),pop=F,SB=F,SF=F,Rho=T,Check=F,BW=F,NC=1) 

      
      # MSMC2
      setwd("~/MSMC2")
      Os_=as.matrix(O_total[[1]])
      create_realinput_msmc2(Os_,paste("Figure_3_MSMC2",sharp,x,sep=""),num=T)
      command_line=paste("msmc2_linux64bit -o",paste("Figure_3_MSMC2_output",sharp,x,sep=""),paste("Figure_3_MSMC2",sharp,x,".txt",sep=""),"-p 20*2 -t 1 -r ",1,sep = " ")
      system(command_line)

      
      
    }
  }
  
  if(method==3){
    M=4
    L=10^8
    
    for(x in 1:nsim){
      path=paste("Figure_3",sharp,M,x,".txt",sep="")
      O_total=Get_sim_data(path,L,M,1)
      Os_=as.matrix(O_total[[1]])
      
      # MSMC
      
      setwd("~/MSMC")
      create_realinput_msmc2(Os_,paste("Figure_3_MSMC",sharp,x,sep=""),num=T)
      command_line=paste("msmc_1.0.0_linux64bit -o",paste("Figure_3_MSMC_output",sharp,x,sep=""),paste("Figure_3_MSMC",sharp,x,".txt",sep="") ,"-p 20*2  -t 1 -r ",1,sep = " ")
      system(command_line)
    }
  }
  
}else{
  M=20
  L=10^8
  # SMC ++ 
  for(x in 1:nsim){
    name=paste("Figure_3","_",strentgh[sharp],"_",x,".vcf",sep="")
    command_line=paste("bcftools view  ",name," -Oz -o ",paste(name,".gz",sep=""),sep="")
    system(command_line)
    command_line=paste("bcftools index ",paste(name,".gz",sep=""),sep="")
    system(command_line)
    
    name_file=paste(name,".gz",sep="")
    output_name_file=paste("Figure_3",strentgh[sharp],x,"output.txt",sep="_")
    sample=paste("1:",paste(paste("tsk",seq(0,((M/2)-1),1),sep="_"),collapse = ","),sep = "")
    command_line= paste(paste("smc++ vcf2smc",name_file,output_name_file,"1",sample,"-l",as.integer(L),sep=" "),paste("smc++ estimate  ",mu," ",output_name_file," --base ",paste("Figure_3_SMC++",sharp,x,sep="")," --cores 2  --unfold --spline cubic --timepoints 100 200000 --algorithm Powell  ", sep=""),paste("smc++ plot ",paste("Figure_3_SMC++",sharp,x,".png",sep=""),paste("Figure_3_SMC++",sharp,x,".final.json",sep=""),"--csv",sep=" "),sep=";")
    system(command_line)
  }
}
}
}

      
    

  
```
