## Supplementary material for "Limits and Convergence properties of the Sequentially Markovian Coalescent": command lines for analyses: Figure_4.R

```
###################
# Source function #
###################
library(eSMC2)
########
#Script#
########
results_esmc=list()
L=10^8
nsim=10
Pop=10^4
mu=1.25*10^-8
theta=mu*4*Pop*L
for(rho in c(0.1,1,10)){
  r=rho*10^-8
          for(method in c(1,3,4)){
            if(method<4){
              
              if(method==1){
                M=2
                L=10^8
                
                for(x in 1:nsim){
                  path=paste("Figure_4",r/mu,M,x,".txt",sep="")
                  DNAseqfile=Get_data(path)
                  O_total=Seqlist2data(DNAseqfile,L,M,1)
                  # esmc
                  results_esmc[[(length(results_esmc)+1)]]=eSMC2(n=40,rho=1,O_total[[1]],mu_r=mu,maxit =20,symbol_number=25,BoxB=c(0.05,1),BoxP=c(3,3),Boxr=c(1,1),Boxs=c(0,0.97),pop=F,SB=F,SF=F,Rho=T,Check=F,BW=F,NC=1) 
                  
                  Os_=as.matrix(O_total[[1]])
                  # MSMC2
                  setwd("~/MSMC2")
                  create_realinput_msmc2(Os_,paste("Figure_4_MSMC2",r/mu,x,sep=""),num=T)
                  command_line=paste("msmc2_linux64bit -o",paste("Figure_4_MSMC2",r/mu,x,"output",sep=""),paste("Figure_4_MSMC2",r/mu,x,".txt",sep="") ,"-p 20*2 -t 1 -r ",1,sep = " ")
                  system(command_line)
                 
                  
                }
              }
              
              if(method==3){
                M=4
                L=10^8
                
                for(x in 1:nsim){
                  path=paste("Figure_4",r/mu,M,x,".txt",sep="")
                  DNAseqfile=Get_data(path)
                  O_total=Seqlist2data(DNAseqfile,L,M,1)
                  Os_=as.matrix(O_total[[1]])
                  
                  # MSMC
                  
                  setwd("~/MSMC")
                  create_realinput_msmc2(Os_,paste("Figure_4_MSMC2",r/mu,x,sep=""),num=T)
                  command_line=paste("msmc_1.0.0_linux64bit -o",paste("Figure_4_MSMC2",r/mu,x,"output",sep=""),paste("Figure_4_MSMC2",r/mu,x,".txt",sep=""),"-p 20*2  -t 1 -r ",1,sep = " ")
                  system(command_line)
                  
                }
              }
              
            }else{
              M=20
              L=10^8
              # SMC ++ 
              setwd("~/Documents/New_msprime/msprime")
              rho_n=c("0.1","1.0","10.0")
              
              for(x in 1:nsim){
                name=paste("Figure_4_rho_",rho_n[count_rho],paste(x,".vcf",sep=""),sep="_")
                command_line=paste("bcftools view  ",name," -Oz -o ",paste(name,".gz",sep=""),sep="")
                system(command_line)
                command_line=paste("bcftools index ",paste(name,".gz",sep=""),sep="")
                system(command_line)
                
                name_file=paste(name,".gz",sep="")
                output_name_file=paste(paste("Figure_4_r_",vect_rho[count_rho],sep=""),x,".txt",sep="_")
                sample=paste("1:",paste(paste("tsk_",seq(0,((M/2)-1),2),sep=""),collapse = ","),sep = "")
                command_line= paste(paste("smc++ vcf2smc",name_file,output_name_file,"1",sample,"-l",as.integer(L),sep=" "),paste("smc++ estimate  ",mu," ",output_name_file," --base ",paste("Figure_4_r_",vect_rho[count_rho],x,sep="")," --cores 1  --unfold --spline cubic --timepoints 100 200000  ", sep=""),paste("smc++ plot ",paste("Figure_4_r_",vect_rho[count_rho],x,".png",sep=""),paste("Figure_4_r_",vect_rho[count_rho],x,".final.json",sep=""),"--csv",sep=" "),sep=";")
                system(command_line)
                
              }
            }
          }
  }
```
