## Supplementary material for "Limits and Convergence properties of the Sequentially Markovian Coalescent": command lines for analyses: Figure_6.R

```
###################
# Source function #
###################
library(eSMC2)
########
#Script#
########
results_esmc=list()
L=2*10^6
nsim=10
Pop=10^4
mu=1.25*10^-8
r=1*10^-8
theta=mu*4*Pop*L
nsim=10
Pop=10^4
M=2
name_issue=c("Figure_6","Figure_6_rec_issue","Figure_6_mu_issue","Figure_6_mut_and_rec_issue")
for(problem in 1:4){
  name=name_issue[problem]
  
    n=40
    for(x in 1:nsim){
      Os_=list()
      for(chr in 1:20){
        path=paste(name,x,chr,".txt",sep="")
      Os_[[chr]]mat=Get_sim_data(path,L,M,1)[[1]]
      }
      # esmc
      results_esmc[[(1+length(results_esmc))]]=eSMC2(n=n,rho=1,Os_,mu_r=mu,maxit =20,symbol_number=25,BoxP=c(3,3),Boxr=c(1,1),pop=F,SB=F,SF=F,Rho=T,NC=20) 
      
      
      results_esmc[[(1+length(results_esmc))]]=eSMC2(n=n,rho=1,Os_,mu_r=mu,maxit =20,symbol_number=25,BoxP=c(3,3),Boxr=c(1,1),pop=F,SB=F,SF=F,Rho=T,NC=20,Share_r = F) 
      

      
    }
  
}
```
