## Supplementary material for "Limits and Convergence properties of the Sequentially Markovian Coalescent": command lines for analyses: Figure_7.R

```
###################
# Source function #
###################
library(eSMC2)
########
#Script#
########
results_esmc=list()
L=20*10^6
nsim=10
Pop=10^4
mu=1.25*10^-8
r=1.25*10^-8
theta=mu*4*Pop*L
nsim=10
Pop=10^4
M=4

for(p in c(0.05,0.1,0.25,0.50)){
  for(kb in c(1000)){
    for(trim in 1:4){
    n=30
    for(x in 1:nsim){
      Os_=Get_real_data(M=M,filename=paste("Figure_7",x,p,kb,trim,".txt",sep=""))
      # esmc
      results_esmc[[(1+length(results_esmc))]]=eSMC2(n=n,rho=1,O_total[[x]],mu_r=mu,maxit =20,symbol_number=30,BoxP=c(3,3),Boxr=c(1,1),pop=F,SB=F,SF=F,Rho=T,NC=1)
      }
  }
  }
}
```
