## Supplementary material for "Limits and Convergence properties of the Sequentially Markovian Coalescent": command lines for analyses: S4_fig.R

```
###################
# Source function #
###################

library(eSMC2)
########
#Script#
########
results=list()
  L=10^9
  M=2
  nsim=10
  Pop=10^4
  mu=1.25*10^-8
  r=1*10^-8
  theta=mu*4*Pop*L
  sharp=4
  for(scenario in c(1,2,3,4)){
    for(x in 1:nsim){
      results[[(1+length(results))]]=Optimize_N(file=paste("S4_fig",sharp,scenario,x,".txt",sep=""),theta = theta,Rho=theta,L=L,n=30,ER=T,Pop=T,SB=F,SF=F,BoxP=c(3,3),Boxr=c(1,1),pop_vect=rep(2,20),Big_Window=F,Correct_window =T,mut=T)
      results[[(1+length(results))]]=Optimize_N(file=paste("S4_fig",sharp,scenario,x,".txt",sep=""),theta = theta,Rho=theta,L=L,n=50,ER=T,Pop=T,SB=F,SF=F,BoxP=c(3,3),Boxr=c(1,1),pop_vect=rep(2,20),Big_Window=F,Correct_window =T,mut=T)
      
    }
    
  }
```
