## Supplementary material for "Limits and Convergence properties of the Sequentially Markovian Coalescent": command lines for analyses: S5_fig.R

```
###################
# Source function #
###################

library(eSMC2)
########
#Script#
########
results=list()
for(data in c(1,2,3,4)){
  if(data==1){
    L=10^7
    M=4
  }
  if(data==2){
    L=10^8
    M=4
    
  }
  if(data==3){
    L=10^9
    M=4
  }
  if(data==4){
    L=10^10
    M=4
    
  }
  nsim=10
  Pop=10^4
  mu=1.25*10^-8
  r=1*10^-8
  theta=mu*4*Pop*L
  for(sharp in c(1,2,3,4)){
    for(x in 1:nsim){
      results[[(1+length(results))]]=Optimize_MM_N(file=paste("S5_fig",sharp,data,x,".txt",sep=""),theta=theta_o,gamma=1,L=L,n=40,ER=T,Pop=T,SB=F,SF=F,BoxB=c(0.1,1),BoxP=c(3,3),Boxr=c(1,1),Boxs=c(0,0.97),pop_vect=NA,window_scaling=c(1,0),sigma=0,beta=1,NC=1,M_a=M,mut=T,Correct_window=T)
            
    }
  }
}
```
