## Supplementary material for "Limits and Convergence properties of the Sequentially Markovian Coalescent": command lines for analyses: S9_fig.R

```
###################
# Source function #
###################

library(eSMC2)
########
#Script#
########
results=list()
    L=10^9
    M=4
  nsim=10
  Pop=10^4
  mu=1.25*10^-8
  r=1*10^-8
  theta=mu*4*Pop*L
  sharp=4
  for(scenario in c(1,2,3,4)){
    for(x in 1:nsim){
      results[[(1+length(results))]]=Optimize_MM_N(file=paste("S9_fig",sharp,scenario,x,".txt",sep=""),theta=theta_o,gamma=1,L=L,n=30,ER=T,Pop=T,SB=F,SF=F,BoxP=c(3,3),Boxr=c(1,1),sigma=0,beta=1,NC=1,M_a=M,mut=T,Correct_window=T)
      results[[(1+length(results))]]=Optimize_MM_N(file=paste("S9_fig",sharp,scenario,x,".txt",sep=""),theta=theta_o,gamma=1,L=L,n=50,ER=T,Pop=T,SB=F,SF=F,BoxP=c(3,3),Boxr=c(1,1),sigma=0,beta=1,NC=1,M_a=M,mut=T,Correct_window=T)
      
    }
  }
```
