## Supplementary material for "Limits and Convergence properties of the Sequentially Markovian Coalescent": command lines for analyses: S12_fig.R

```
###################
# Source function #
###################
library(eSMC2)
########
#Script#
########
results_esmc=list()
L=10^8
nsim=10
Pop=10^4
mu=1.25*10^-8
r=1*10^-8
theta=mu*4*Pop*L
for(method in c(1,3,4)){
  if(method<4){
    if(method==1){
      M=2
      L=10^8
      for(x in 1:nsim){
        path=paste("S12_fig",M,x,".txt",sep="")
        O_total=Get_sim_data(path,L,M,1)
        # esmc
        results_esmc[[(1+length(results_esmc))]]=eSMC2(n=40,rho=1,O_total[[1]],mu_r=mu,maxit =20,symbol_number=25,BoxB=c(0.05,1),BoxP=c(3,3),Boxr=c(1,1),Boxs=c(0,0.97),pop=F,SB=F,SF=F,Rho=T,Check=F,BW=F,NC=1) 
        
        
        # MSMC2
        setwd("~/MSMC2")
        Os_=as.matrix(O_total[[1]])
        create_realinput_msmc2(Os_,paste("S12_fig_MSMC2",x,sep=""),num=T)
        command_line=paste("msmc2_linux64bit -o",paste("S12_fig_MSMC2_output",x,sep=""),paste("S12_fig_MSMC2",x,".txt",sep=""),"-p 20*2 -t 1 -r ",1,sep = " ")
        system(command_line)
        
        
        
      }
    }
    
    if(method==3){
      M=4
      L=10^8
      
      for(x in 1:nsim){
        path=paste("S12_fig",M,x,".txt",sep="")
        O_total=Get_sim_data(path,L,M,1)
        Os_=as.matrix(O_total[[1]])
        
        # MSMC
        
        setwd("~/MSMC")
        create_realinput_msmc2(Os_,paste("S12_fig_MSMC",x,sep=""),num=T)
        command_line=paste("msmc_1.0.0_linux64bit -o",paste("S12_fig_MSMC_output",x,sep=""),paste("S12_fig_MSMC",x,".txt",sep="") ,"-p 20*2  -t 1 -r ",1,sep = " ")
        system(command_line)
      }
    }
    
  }else{
    M=20
    L=10^8
    # SMC ++ 
    for(x in 1:nsim){
      name=paste("S12_fig","_","_",x,".vcf",sep="")
      command_line=paste("bcftools view  ",name," -Oz -o ",paste(name,".gz",sep=""),sep="")
      system(command_line)
      command_line=paste("bcftools index ",paste(name,".gz",sep=""),sep="")
      system(command_line)
      
      name_file=paste(name,".gz",sep="")
      output_name_file=paste("S12_fig",x,"output.txt",sep="_")
      sample=paste("1:",paste(paste("tsk",seq(0,((M/2)-1),1),sep="_"),collapse = ","),sep = "")
      command_line= paste(paste("smc++ vcf2smc",name_file,output_name_file,"1",sample,"-l",as.integer(L),sep=" "),paste("smc++ estimate  ",mu," ",output_name_file," --base ",paste("S12_fig_SMC++",x,sep="")," --cores 2  --unfold --spline cubic --timepoints 100 200000 --algorithm Powell  ", sep=""),paste("smc++ plot ",paste("S12_fig_SMC++",x,".png",sep=""),paste("S12_fig_SMC++",x,".final.json",sep=""),"--csv",sep=" "),sep=";")
      system(command_line)
    }
  }
}
```
