## Supplementary material for "Limits and Convergence properties of the Sequentially Markovian Coalescent": command lines for analyses: S13_fig.R

```
###################
# Source function #
###################

library(eSMC2)
########
#Script#
########
results_esmc=list()
L=10^8
nsim=10
Pop=10^4
mu=1.25*10^-8
r=1*10^-8
theta=mu*4*Pop*L
strentgh=c(2,5,10,50)
for(sharp in c(1,2,3,4)){
  for(method in c(1,3,4)){
    if(method<4){
      if(method==1){
        M=2
        L=10^8
        for(x in 1:nsim){
          path=paste("S13_fig",sharp,M,x,".txt",sep="")
          O_total=Get_sim_data(path,L,M,1)
          # esmc
          results_esmc[[(1+length(results_esmc))]]=eSMC2(n=40,rho=1,O_total[[1]],mu_r=mu,maxit =20,symbol_number=25,BoxB=c(0.05,1),BoxP=c(3,3),Boxr=c(1,1),Boxs=c(0,0.97),pop=F,SB=F,SF=F,Rho=T,Check=F,BW=F,NC=1) 
          
          
          # MSMC2
          setwd("~/MSMC2")
          Os_=as.matrix(O_total[[1]])
          create_realinput_msmc2(Os_,paste("S13_fig_MSMC2",sharp,x,sep=""),num=T)
          command_line=paste("msmc2_linux64bit -o",paste("S13_fig_MSMC2_output",sharp,x,sep=""),paste("S13_fig_MSMC2",sharp,x,".txt",sep=""),"-p 20*2 -t 1 -r ",1,sep = " ")
          system(command_line)
          
          
          
        }
      }
      
      if(method==3){
        M=4
        L=10^8
        
        for(x in 1:nsim){
          path=paste("S13_fig",sharp,M,x,".txt",sep="")
          O_total=Get_sim_data(path,L,M,1)
          Os_=as.matrix(O_total[[1]])
          
          # MSMC
          
          setwd("~/MSMC")
          create_realinput_msmc2(Os_,paste("S13_fig_MSMC",sharp,x,sep=""),num=T)
          command_line=paste("msmc_1.0.0_linux64bit -o",paste("S13_fig_MSMC_output",sharp,x,sep=""),paste("S13_fig_MSMC",sharp,x,".txt",sep="") ,"-p 20*2  -t 1 -r ",1,sep = " ")
          system(command_line)
        }
      }
      
    }else{
      M=20
      L=10^8
      # SMC ++ 
      for(x in 1:nsim){
        name=paste("S13_fig","_",strentgh[sharp],"_",x,".vcf",sep="")
        command_line=paste("bcftools view  ",name," -Oz -o ",paste(name,".gz",sep=""),sep="")
        system(command_line)
        command_line=paste("bcftools index ",paste(name,".gz",sep=""),sep="")
        system(command_line)
        
        name_file=paste(name,".gz",sep="")
        output_name_file=paste("S13_fig",strentgh[sharp],x,"output.txt",sep="_")
        sample=paste("1:",paste(paste("tsk",seq(0,((M/2)-1),1),sep="_"),collapse = ","),sep = "")
        command_line= paste(paste("smc++ vcf2smc",name_file,output_name_file,"1",sample,"-l",as.integer(L),sep=" "),paste("smc++ estimate  ",mu," ",output_name_file," --base ",paste("S13_fig_SMC++",sharp,x,sep="")," --cores 2  --unfold --spline cubic --timepoints 100 200000 --algorithm Powell  ", sep=""),paste("smc++ plot ",paste("S13_fig_SMC++",sharp,x,".png",sep=""),paste("S13_fig_SMC++",sharp,x,".final.json",sep=""),"--csv",sep=" "),sep=";")
        system(command_line)
      }
    }
  }
}
```
