## Supplementary material for "Limits and Convergence properties of the Sequentially Markovian Coalescent": command lines for analyses: S15_fig.R

```
###################
# Source function #
###################
library('plot.matrix')
library(eSMC2)
########
#Script#
########
if(T){
  gen <- 1
  nsim=10
  mu=1.25*10^-8
  r=1*10^-8
  M=2
  L=10^8
  pdf("S15_fig.pdf")
  par(mfrow=c(2,2))
  sharp_v=c(2,5,10,50)
  for(sharp in 1:4){
    N_theo=matrix(0,nrow = 40*40,ncol = 10)
    N=matrix(0,nrow = 40*40,ncol = 10)
    for(x in 1:nsim){
      path=paste("S15_fig",sharp,M,x,".txt",sep="")
      O_total=Get_sim_data(path,L,M,1)
      # esmc
      results=eSMC2(n=40,rho=1,O_total[[1]],mu_r=mu,maxit =20,symbol_number=25,BoxB=c(0.05,1),BoxP=c(3,3),Boxr=c(1,1),Boxs=c(0,0.97),pop=F,SB=F,SF=F,Rho=T,NC=1) 
      
      N=results$N
      BW_mat=get_Mat(file=paste("S15_fig",sharp,x,".txt",sep=""),theta=theta,Rho=theta,L=L,n=40,BoxB=c(0.1,1),BoxP=c(3,3),Boxr=c(1,1),Boxs=c(0,0.97),pop_vect=rep(2,20),window_scaling=c(1,0),Big_Window=0,mut=T,Correct_window = T)
      N_theo[,x]=as.numeric(BW_mat$N)
      
      N[which(is.na(as.numeric(N[,x]))),x]<-0
      N_theo[which(is.na(as.numeric(N_theo[,x]))),x]<-0
    }
    dist_vect=numeric(length = dim(N)[1])
    for(ii in 1:dim(N_theo)[1]){
      dist_vect[ii]=mean((N[ii,]-N_theo[ii,]))
    }
    N_heat=matrix(dist_vect,nrow = 40,ncol = 40)
    if(sharp<2){
      plot(N_heat,border=NA,breaks=seq(-2*10^7,10^7,2*10^6),  axis.col=NULL, axis.row=NULL, xlab='', ylab='',main = "",key=list( cex.axis=0.75))
    }else{
      plot(N_heat,border=NA,breaks=seq(-2*10^7,10^7,2*10^6),  axis.col=NULL, axis.row=NULL , xlab='', ylab='' ,main = "",key=NULL)
    }
    title(paste(letters[sharp],")"," Amplitde: ",sharp_v[sharp],sep=""),adj = 0)
    
  }
  dev.off()
  
}
```
