## Supplementary material for "Limits and Convergence properties of the Sequentially Markovian Coalescent": command lines for analyses: S16_fig.R

```
###################
# Source function #
###################
library(eSMC2)
########
#Script#
########
results_esmc=list()
M=4
L=50*10^8
nsim=10
Pop=10^4
mu=1.25*10^-8
r=10^-8
theta=4*Pop*mu*L
rho=4*Pop*r*L
for(scenario in c(1,2)){
  for(BW in c(TRUE,FALSE)){
      for(x in 1:nsim){
        path=paste("S16_fig",scenario,x,".txt",sep="")
        O_total=Get_sim_data(path,L,M,1)
        
        results_esmc[[(1+length(results_esmc))]]=eSMC2(n=40,rho=1,O_total[[1]],mu_r=mu,maxit =20,symbol_number=30,BoxP=c(3,3),Boxr=c(1,1),pop=F,SB=F,SF=F,Rho=T,BW,NC=1)

        results_esmc[[(1+length(results_esmc))]]=eSMC2(n=40,rho=1,O_total[[1]],mu_r=mu,maxit =20,symbol_number=30,BoxP=c(3,3),Boxr=c(1,1),pop=F,SB=F,SF=F,Rho=T,BW,Big_Window=2,NC=1)
        
        results_esmc[[(1+length(results_esmc))]]=eSMC2(n=40,rho=1,O_total[[1]],mu_r=mu,maxit =20,symbol_number=30,BoxP=c(3,3),Boxr=c(1,1),pop=F,SB=F,SF=F,Rho=T,BW,window_scaling=c(0.2,0),NC=1)
        
        results_esmc[[(1+length(results_esmc))]]=eSMC2(n=40,rho=1,O_total[[1]],mu_r=mu,maxit =20,symbol_number=30,BoxP=c(3,3),Boxr=c(1,1),pop=F,SB=F,SF=F,Rho=T,BW,window_scaling=c(5,0),NC=1)
    }
  }
}
```
