## Supplementary material for "Limits and Convergence properties of the Sequentially Markovian Coalescent": command lines for analyses: S18_fig.R

```
###################
# Source function #
###################

library(eSMC2)
########
#Script#
########
results_esmc=list()
L=10^8
nsim=10
Pop=10^4
mu=1.25*10^-8
r=1*10^-8
theta=mu*4*Pop*L
strentgh=c(2,5,10,50)
for(sharp in c(1,2,3,4)){
  

    M=20
    L=10^8
    # SMC ++ 
    for(x in 1:nsim){
      name=paste("S18_fig","_",strentgh[sharp],"_",x,".vcf",sep="")
      command_line=paste("bcftools view  ",name," -Oz -o ",paste(name,".gz",sep=""),sep="")
      system(command_line)
      command_line=paste("bcftools index ",paste(name,".gz",sep=""),sep="")
      system(command_line)
      
      name_file=paste(name,".gz",sep="")
      output_name_file=paste("S18_fig",strentgh[sharp],x,"_rp1_output.txt",sep="_")
      sample=paste("1:",paste(paste("tsk",seq(0,((M/2)-1),1),sep="_"),collapse = ","),sep = "")
      command_line= paste(paste("smc++ vcf2smc",name_file,output_name_file,"1",sample,"-l",as.integer(L),sep=" "),paste("smc++ estimate  ",mu," ",output_name_file," --base ",paste("S18_fig_SMC++",sharp,x,sep="")," --cores 2 -rp 1  --unfold --spline cubic --timepoints 100 200000 --algorithm Powell  ", sep=""),paste("smc++ plot ",paste("S18_fig_rp1_SMC++",sharp,x,".png",sep=""),paste("S18_fig_rp1_SMC++",sharp,x,".final.json",sep=""),"--csv",sep=" "),sep=";")
      system(command_line)
      
      name_file=paste(name,".gz",sep="")
      output_name_file=paste("S18_fig",strentgh[sharp],x,"_rp5_output.txt",sep="_")
      sample=paste("1:",paste(paste("tsk",seq(0,((M/2)-1),1),sep="_"),collapse = ","),sep = "")
      command_line= paste(paste("smc++ vcf2smc",name_file,output_name_file,"1",sample,"-l",as.integer(L),sep=" "),paste("smc++ estimate  ",mu," ",output_name_file," --base ",paste("S18_fig_SMC++",sharp,x,sep="")," --cores 2 -rp 5 --unfold --spline cubic --timepoints 100 200000 --algorithm Powell  ", sep=""),paste("smc++ plot ",paste("S18_fig_rp5_SMC++",sharp,x,".png",sep=""),paste("S18_fig_rp5_SMC++",sharp,x,".final.json",sep=""),"--csv",sep=" "),sep=";")
      system(command_line)
      
      name_file=paste(name,".gz",sep="")
      output_name_file=paste("S18_fig",strentgh[sharp],x,"_rp10_output.txt",sep="_")
      sample=paste("1:",paste(paste("tsk",seq(0,((M/2)-1),1),sep="_"),collapse = ","),sep = "")
      command_line= paste(paste("smc++ vcf2smc",name_file,output_name_file,"1",sample,"-l",as.integer(L),sep=" "),paste("smc++ estimate  ",mu," ",output_name_file," --base ",paste("S18_fig_SMC++",sharp,x,sep="")," --cores 2 -rp 10 --unfold --spline cubic --timepoints 100 200000 --algorithm Powell  ", sep=""),paste("smc++ plot ",paste("S18_fig_rp10_SMC++",sharp,x,".png",sep=""),paste("S18_fig_rp10_SMC++",sharp,x,".final.json",sep=""),"--csv",sep=" "),sep=";")
      system(command_line)
      
      name_file=paste(name,".gz",sep="")
      output_name_file=paste("S18_fig",strentgh[sharp],x,"_rp20_output.txt",sep="_")
      sample=paste("1:",paste(paste("tsk",seq(0,((M/2)-1),1),sep="_"),collapse = ","),sep = "")
      command_line= paste(paste("smc++ vcf2smc",name_file,output_name_file,"1",sample,"-l",as.integer(L),sep=" "),paste("smc++ estimate  ",mu," ",output_name_file," --base ",paste("S18_fig_SMC++",sharp,x,sep="")," --cores 2 -rp 20 --unfold --spline cubic --timepoints 100 200000 --algorithm Powell  ", sep=""),paste("smc++ plot ",paste("S18_fig_rp20_SMC++",sharp,x,".png",sep=""),paste("S18_fig_rp20_SMC++",sharp,x,".final.json",sep=""),"--csv",sep=" "),sep=";")
      system(command_line)
      
    }
  
}
```
