## Supplementary material for "Limits and Convergence properties of the Sequentially Markovian Coalescent": command lines for analyses: S23_fig.R

```
###################
# Source function #
###################
library(eSMC2)
########
#Script#
########
results_esmc=list()
L=10^8
nsim=10
Pop=10^4
mu=1.25*10^-8
r=1.25*10^-8
theta=mu*4*Pop*L
    nsim=10
    Pop=10^4
    M=4
    for(problem in 1:3){
              for(p in c(0,0.05,0.1,0.25)){
              n=30
              for(x in 1:nsim){
                Os_=Get_real_data(M=M,filename=paste("S23_fig",problem,x,p,sep=""))
                # esmc
                results_esmc[[(1+length(results_esmc))]]=eSMC2(n=n,rho=1,Os_,mu_r=mu,maxit =20,symbol_number=30,BoxB=c(0.05,1),BoxP=c(3,3),Boxr=c(1,1),Boxs=c(0,0.97),pop=F,SB=F,SF=F,Rho=T,Check=F,BW=F,NC=1) 
              }
            }
}
        
      
    
    
  
```
