## Supplementary material for "Limits and Convergence properties of the Sequentially Markovian Coalescent": command lines for analyses: S24_fig.R

```
###################
# Source function #
###################
library(eSMC2)
########
#Script#
########
results_esmc=list()
L=10^8
nsim=10
Pop=10^4
mu=1.25*10^-8
r=1.25*10^-8
theta=mu*4*Pop*L
nsim=10
Pop=10^4
M=4
for(problem in 1:3){
  for(p in c(0,0.05,0.1,0.25)){
    for(x in 1:nsim){
      command_line=paste("msmc_linux64bit -o ",paste("S24_fig",problem,x,p,"output",sep=""),paste("S20_fig",problem,x,p,".txt",sep="")," -p 15*2 -r ",1,sep = " ")
      system(command_line)
    }
  }
}
```
