## Supplementary material for "Limits and Convergence properties of the Sequentially Markovian Coalescent": command lines for analyses: S26_fig.R

```
###################
# Source function #
###################
library(eSMC2)
########
#Script#
########
results_esmc=list()
L=4*10^7
nsim=10
Pop=10^4
mu=1.25*10^-8
r=1.25*10^-8
theta=mu*4*Pop*L
nsim=10
Pop=10^4
M=2
    n=40
    for(x in 1:nsim){
      path=paste("S26_fig",x,".txt",sep="")
      O_total=Get_sim_data(path,L,M,1)
      
      # esmc
      results_esmc[[(1+length(results_esmc))]]=eSMC2(n=n,rho=1,Os_,mu_r=mu,maxit =20,symbol_number=30,BoxB=c(0.05,1),BoxP=c(3,3),Boxr=c(1,1),Boxs=c(0,0.97),pop=F,SB=F,SF=F,Rho=T,Check=F,BW=F,NC=1) 
    
      path=paste("S26_fig_rec_issue",x,".txt",sep="")
      O_total=Get_sim_data(path,L,M,1)
      
      # esmc
      results_esmc[[(1+length(results_esmc))]]=eSMC2(n=n,rho=1,Os_,mu_r=mu,maxit =20,symbol_number=30,BoxB=c(0.05,1),BoxP=c(3,3),Boxr=c(1,1),Boxs=c(0,0.97),pop=F,SB=F,SF=F,Rho=T,Check=F,BW=F,NC=1) 
      
      }
    
  
```
