## Supplementary material for "Limits and Convergence properties of the Sequentially Markovian Coalescent": command lines for analyses: S31_fig.R

```
###################
# Source function #
###################
library(eSMC2)
########
#Script#
########
results_esmc=list()
L=20*10^6
nsim=10
Pop=10^4
mu=1.25*10^-8
r=1.25*10^-8
theta=mu*4*Pop*L
nsim=10
Pop=10^4
M=4

for(p in c(0.05,0.1,0.25,0.50)){
  for(kb in c(100000)){
    for(trim in 1:4){
      n=30
      for(x in 1:nsim){
        command_line=paste("msmc_1.0.0_linux64bit -o ",paste("S31_fig",problem,x,p,"output",sep=""),paste("S31_fig",problem,x,p,".txt",sep="")," -p 15*2 -r ",1,sep = " ")
        system(command_line)
      }
    }
  }
}
```
