## Supplementary material for "Limits and Convergence properties of the Sequentially Markovian Coalescent": Contains supplementary figures and tables

Technical University of Munich

### 1 Supplementary Figures

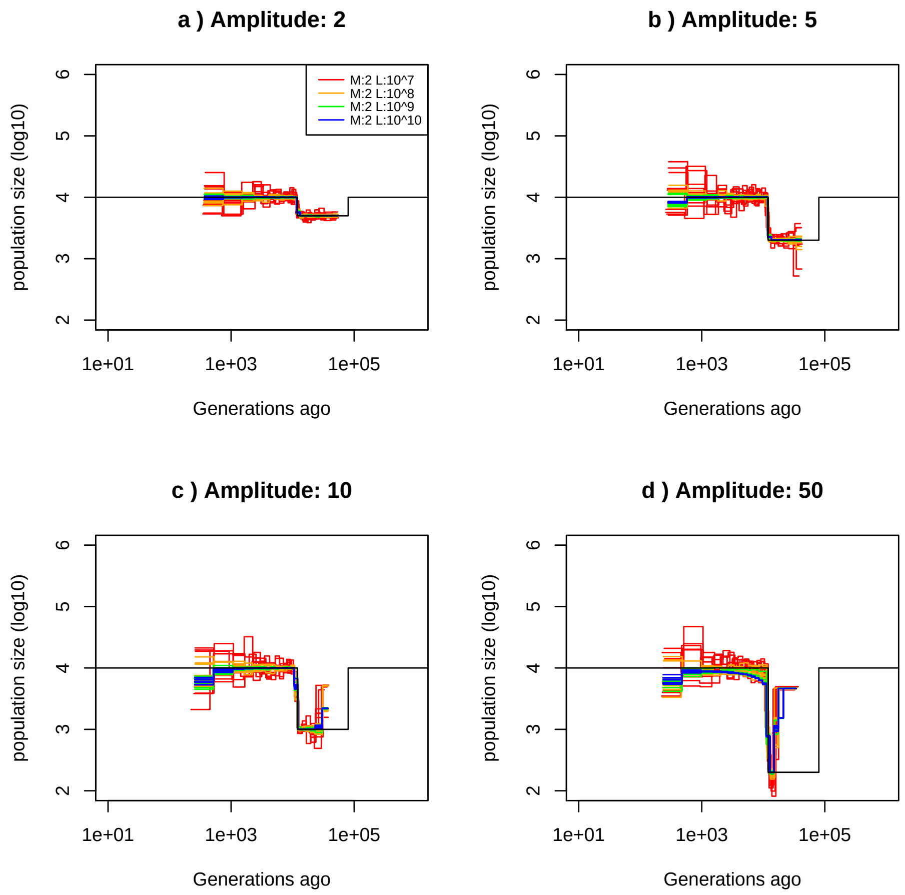


Figure 1: **Best-case convergence of PSMC'** Estimated demographic history using simulated genealogy over sequences of 10,100,1000,10000 Mb (respectively in red,orange, green and blue) under a bottleneck scenario (black) with 10 replicates for different amplitudes of size change: a) 2-fold, b) 5-fold, c) 10-fold, and d) 50-fold. The recombination rate is set to $1 \times 10^{-8}$ per generation per bp and the mutation rate to $1.25 \times 10^{-8}$ per generation per bp.

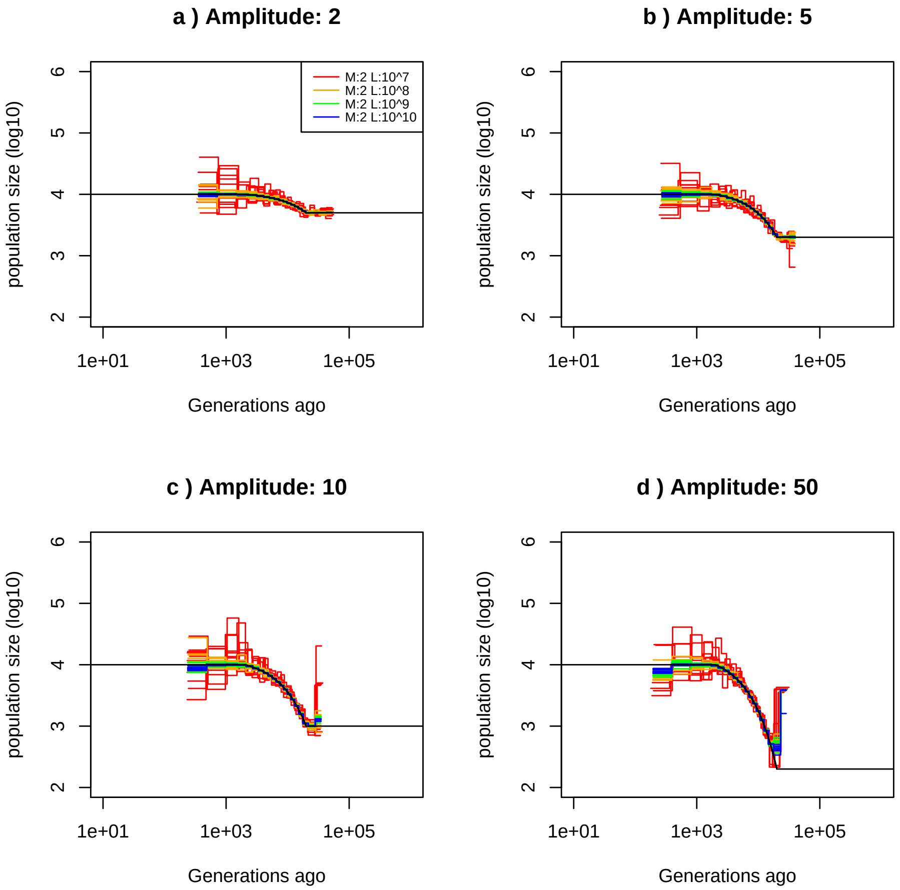


Figure 2: **Best-case convergence of PSMC'** Estimated demographic history using simulated genealogy over sequences of 10,100,1000,10000 Mb (respectively in red,orange, green and blue) under a population expansion scenario (black) with 10 replicates for different amplitudes of size change: a) 2-fold, b) 5-fold, c) 10-fold, and d) 50-fold. The recombination rate is set to $1 \times 10^{-8}$ per generation per bp and the mutation rate to $1.25 \times 10^{-8}$ per generation per bp.

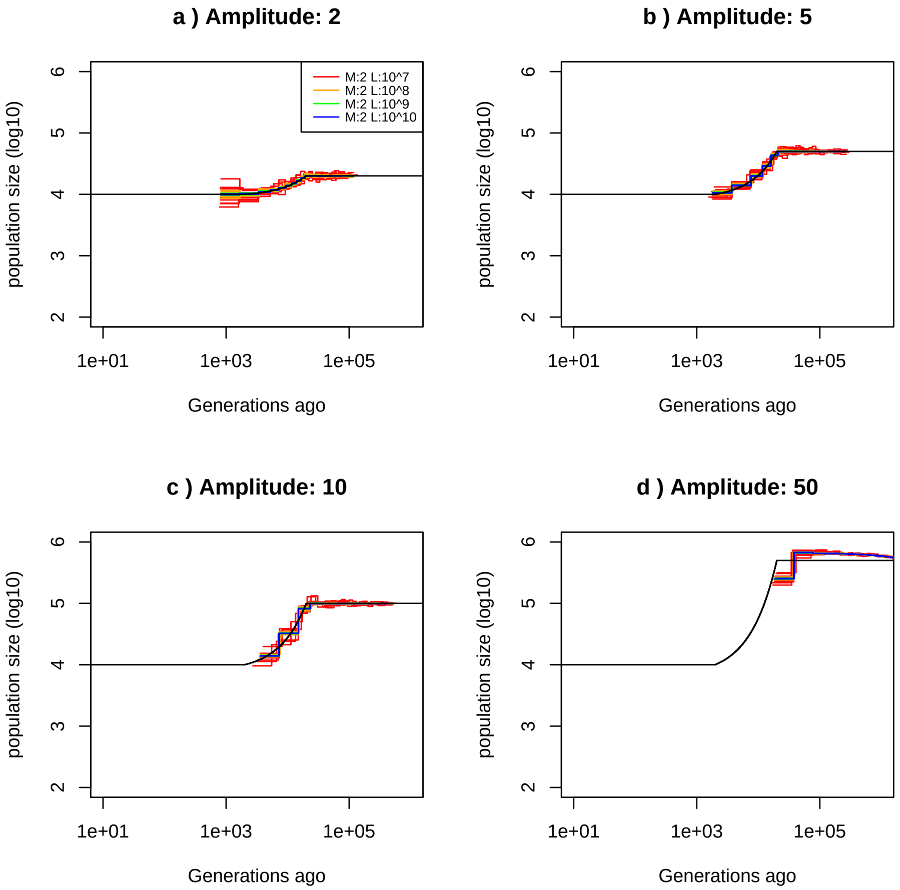


Figure 3: **Best-case convergence of PSMC'** Estimated demographic history using simulated genealogy over sequences of 10,100,1000,10000 Mb (respectively in red,orange, green and blue) under a decrease of population size scenario (black) with 10 replicates for different amplitudes of size change: a) 2-fold, b) 5-fold, c) 10-fold, and d) 50-fold. The recombination rate is set to $1 \times 10^{-8}$ per generation per bp and the mutation rate to $1.25 \times 10^{-8}$ per generation per bp.

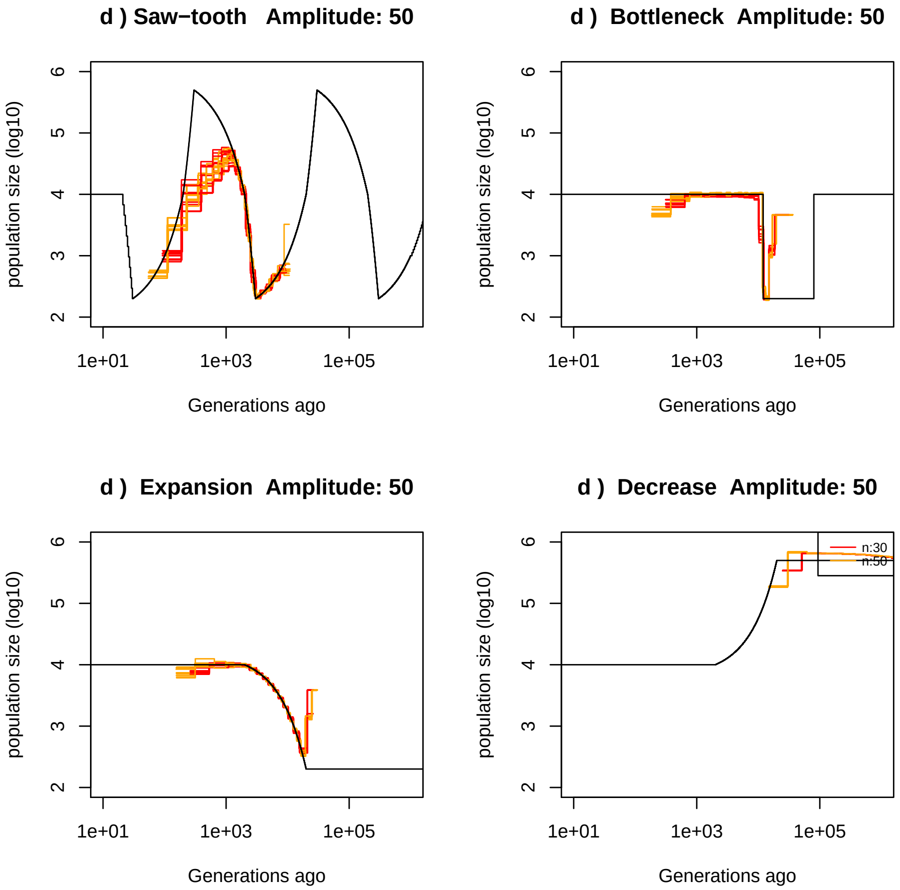


Figure 4: **Best-case convergence of PSMC'** Estimated demographic history using simulated genealogy over sequences of 1 Gb using 30 or 50 hidden states (respectively in red,orange) under scenarios with population size of fold 50 (black) with 10 replicates. Recombination rate is set to $1 \times 10^{-8}$ per generation per bp and mutation rate to $1.25 \times 10^{-8}$ per generation per bp. a) Demographic history simulated under a saw-tooth scenario of strength 50. b) Demographic history simulated under a bottleneck scenario of strength 50. c) Demographic history simulated under a population expansion scenario of strength 50. d) Demographic history simulated under a population decrease scenario of strength 50.

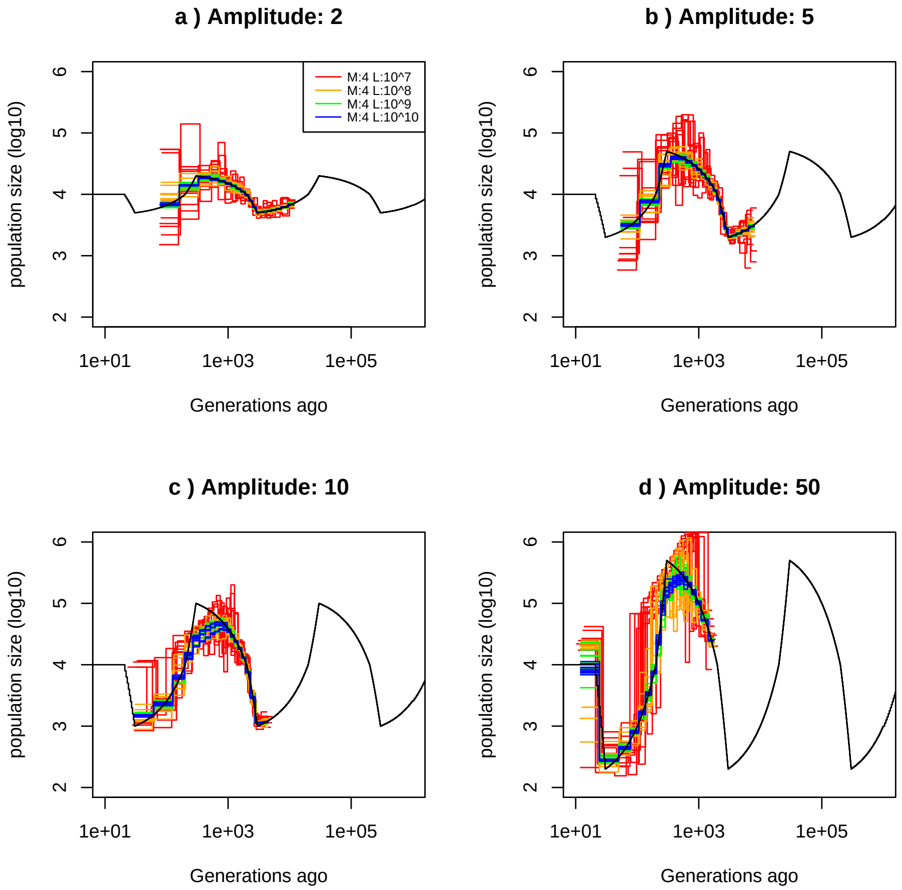


Figure 5: **Best-case convergence of MSMC** Estimated demographic history using simulated genealogies of 4 sequences of 10,100,1000,10000 Mb (respectively in red,orange, green and blue) under a saw-tooth scenario (black) with 10 replicates for different amplitudes of size change: a) 2-fold, b) 5-fold, c) 10-fold, and d) 50-fold. The recombination rate is set to $1 \times 10^{-8}$ per generation per bp and the mutation rate to $1.25 \times 10^{-8}$ per generation per bp.

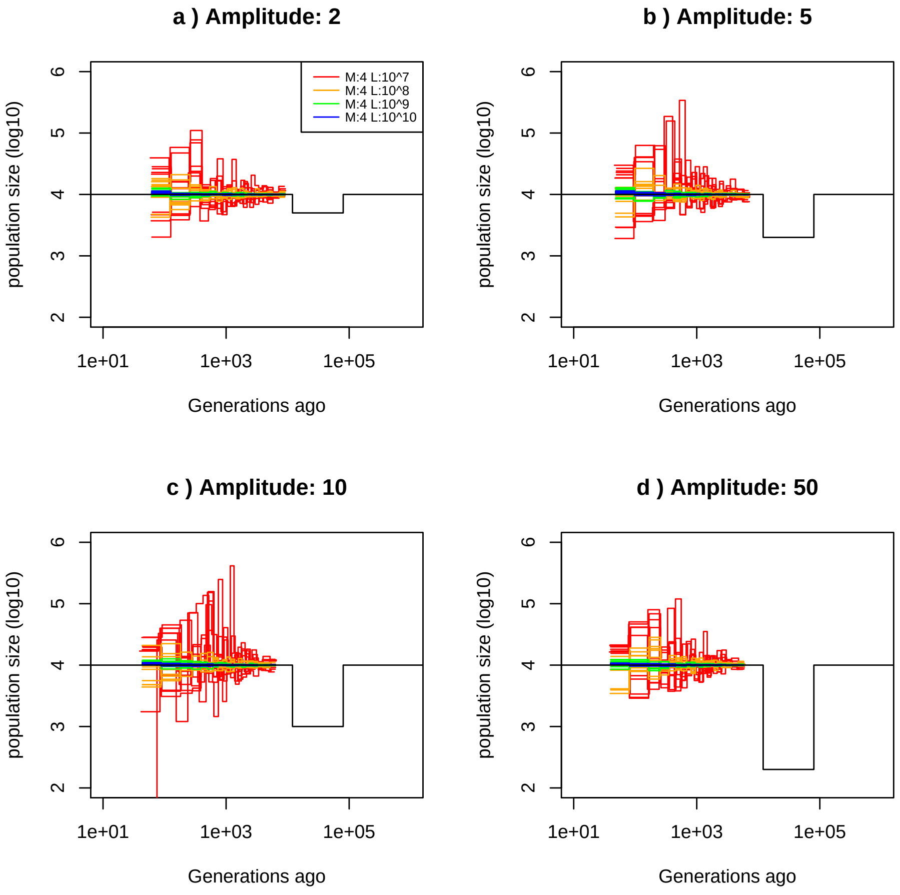


Figure 6: **Best-case convergence of MSMC** Estimated demographic history using simulated genealogies of 4 sequences of 10,100,1000,10000 Mb (respectively in red,orange, green and blue) under a bottleneck scenario (black) with 10 replicates for different amplitudes of size change: a) 2-fold, b) 5-fold, c) 10-fold, and d) 50-fold. The recombination rate is set to $1 \times 10^{-8}$ per generation per bp and the mutation rate to $1.25 \times 10^{-8}$ per generation per bp.

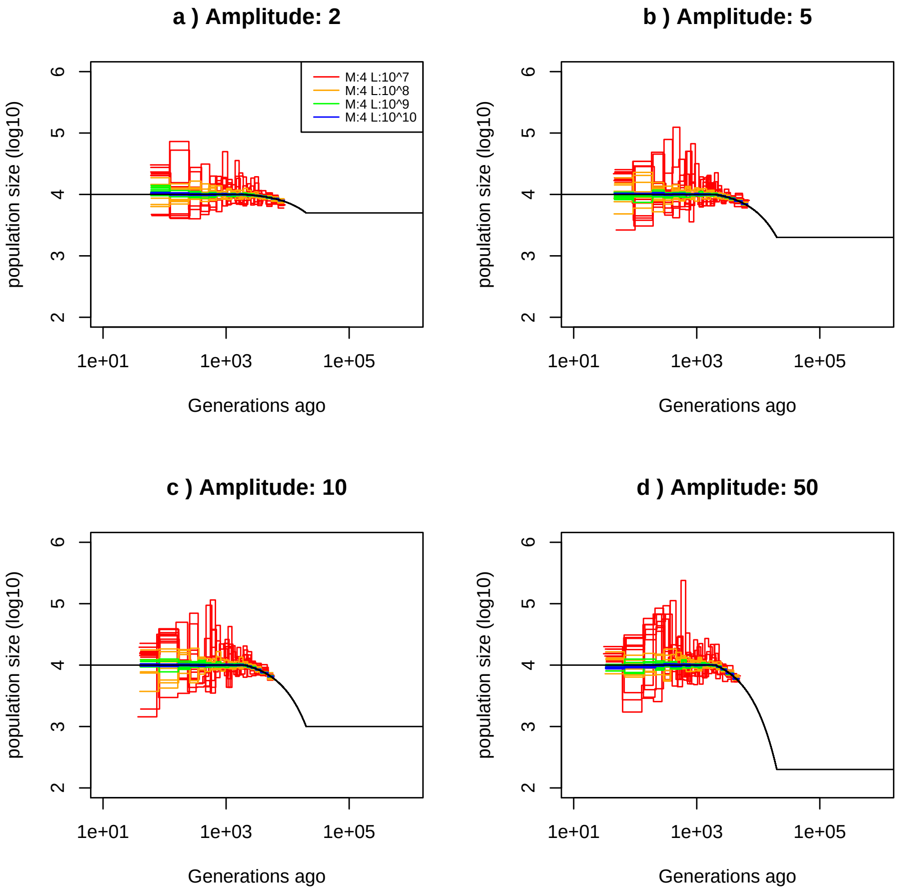


Figure 7: **Best-case convergence of MSMC** Estimated demographic history using simulated genealogies of 4 sequences of 10,100,1000,10000 Mb (respectively in red,orange, green and blue) under a population expansion scenario (black) with 10 replicates for different amplitudes of size change: a) 2-fold, b) 5-fold, c) 10-fold, and d) 50-fold. The recombination rate is set to $1 \times 10^{-8}$ per generation per bp and the mutation rate to $1.25 \times 10^{-8}$ per generation per bp.

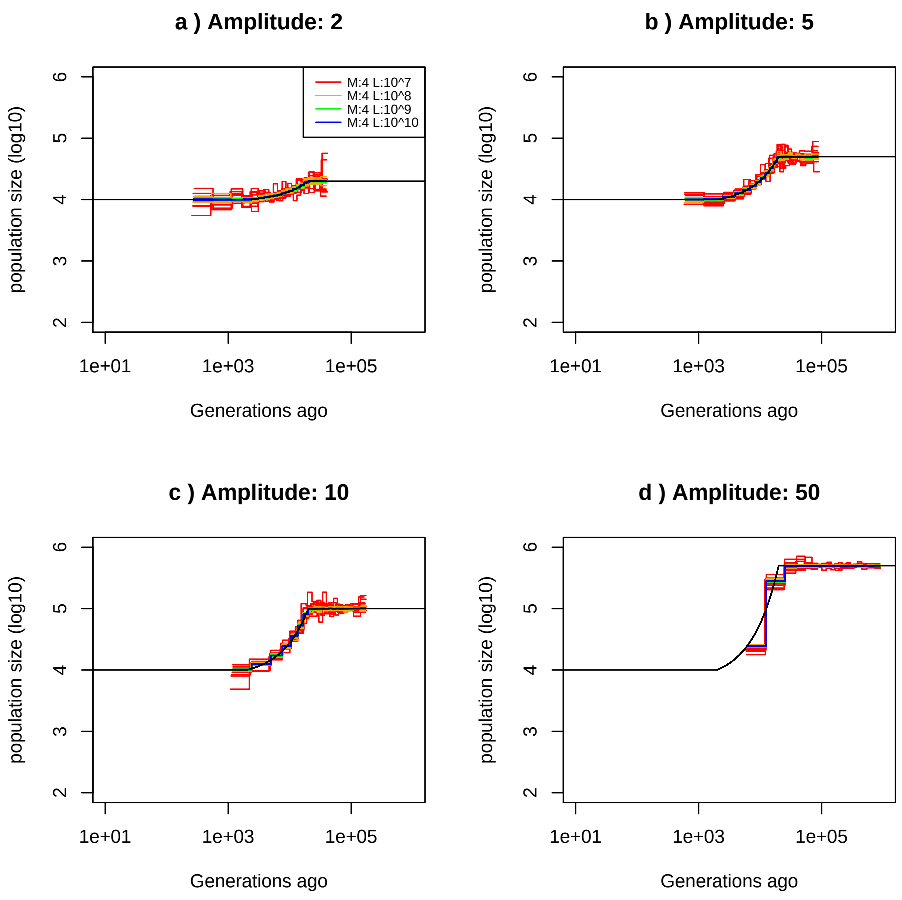


Figure 8: **Best-case convergence of MSMC** Estimated demographic history using simulated genealogies of 4 sequences of 10,100,1000,10000 Mb (respectively in red,orange, green and blue) under a decrease of population size scenario (black) with 10 replicates for different amplitudes of size change: a) 2-fold, b) 5-fold, c) 10-fold, and d) 50-fold. The recombination rate is set to $1 \times 10^{-8}$ per generation per bp and the mutation rate to $1.25 \times 10^{-8}$ per generation per bp.

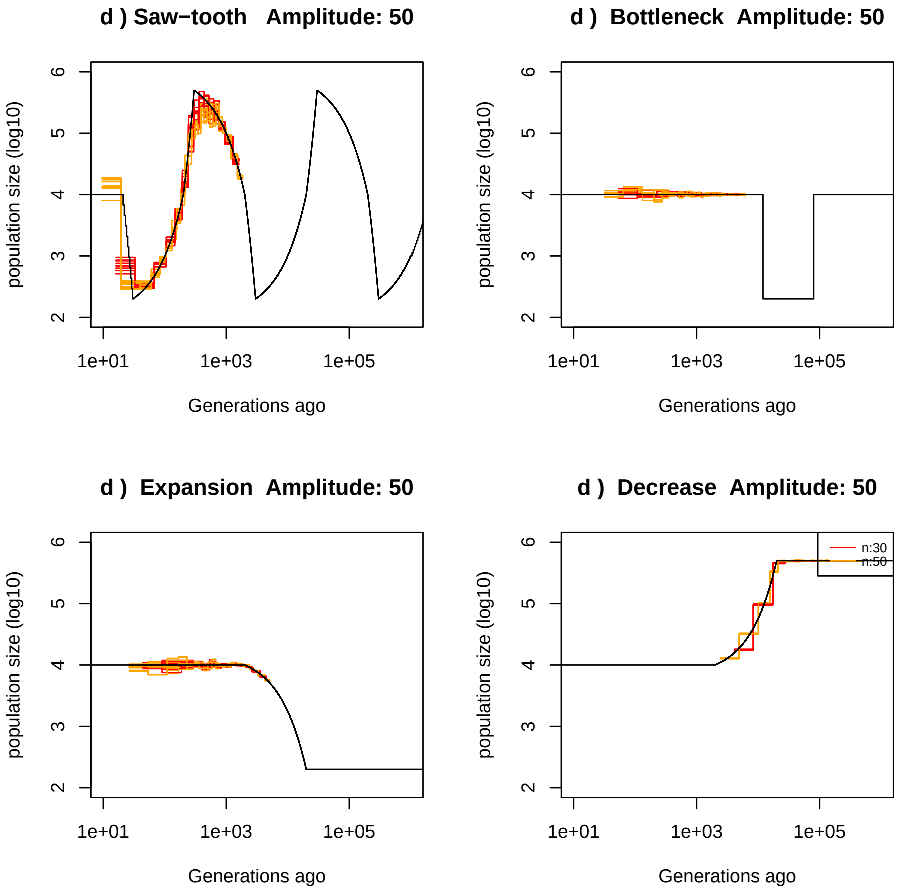


Figure 9: **Best-case convergence of MSMC** Estimated demographic history using simulated genealogy over sequences of 1 Gb using 30 or 50 hidden states (respectively in red,orange) under scenarios with variation of population size fold 50 (black) with 10 replicates. Recombination rate is set to $1 \times 10^{-8}$ per generation per bp and mutation rate to $1.25 \times 10^{-8}$ per generation per bp. a) Demographic history simulated under a saw-tooth scenario of strength 50. b) Demographic history simulated under a bottleneck scenario of strength 50. c) Demographic history simulated under a population expansion scenario of strength 50. d) Demographic history simulated under a population decrease scenario of strength 50.

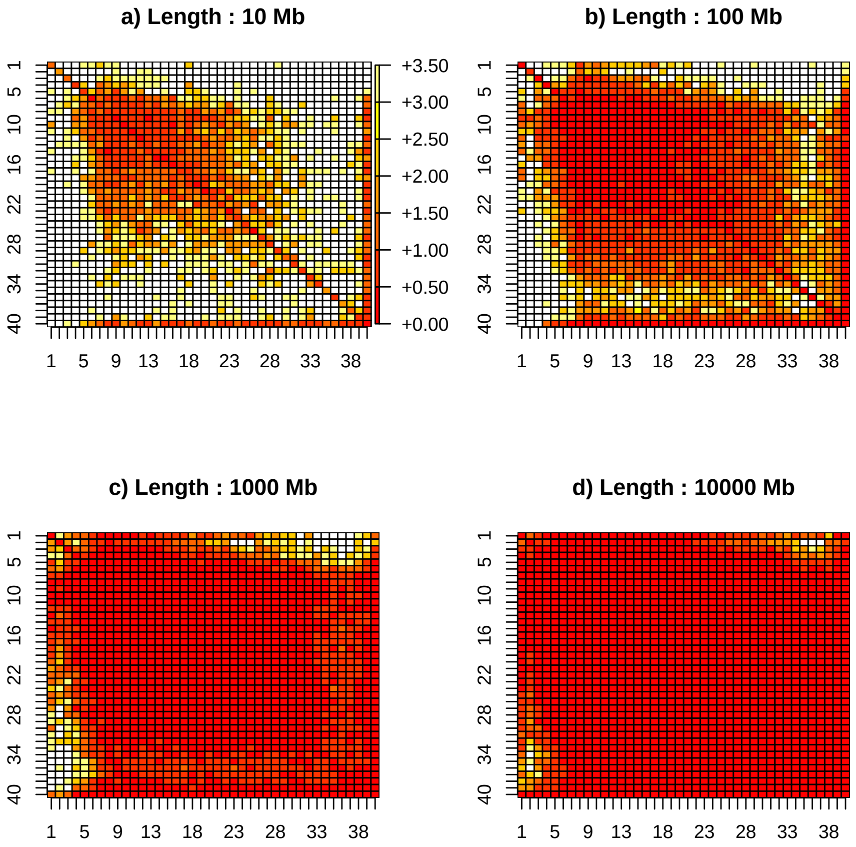


Figure 10: **Estimated Transition matrix in sawtooth scenario** Estimated transition matrix coefficient of variation using simulated genealogy over sequences of 10,100,1000,10000 (respectively in a), b) c) and d) ) Mb under a saw-tooth scenario with 10 replicates. Recombination rate is set to $1 \times 10^{-8}$ per generation per bp and mutation rate to $1.25 \times 10^{-8}$ per generation per bp. Demographic history is simulated under a saw-tooth scenario of amplitude 10.

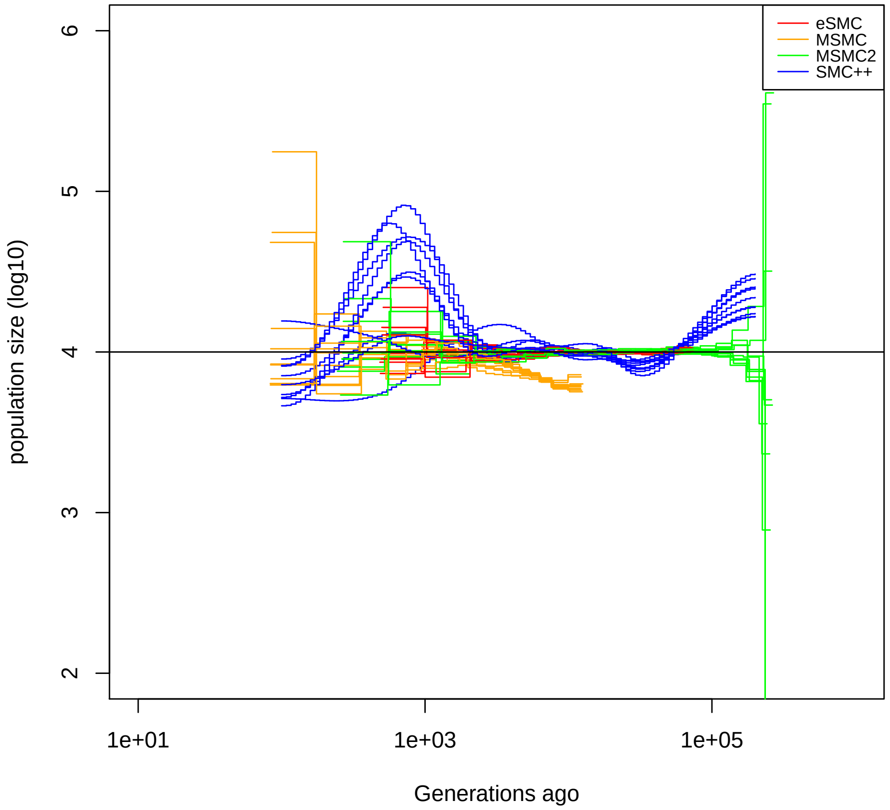


Figure 11: **Estimated demography of SMC method under a constant population size scenario.** Estimated demographic history under a constant population size with 10 replicates using simulated sequences. 2 sequences of 100 Mb for eSMC and MSMC2 (respectively in red and green ). We use 4 sequences of 100 Mb for MSMC (orange) and 20 sequences of 100 Mb for SMC++ (blue). Recombination rate is set to $1 \times 10^{-8}$ per generation per bp and mutation rate to $1.25 \times 10^{-8}$ per generation per bp. The simulated demographic history is represented in black.

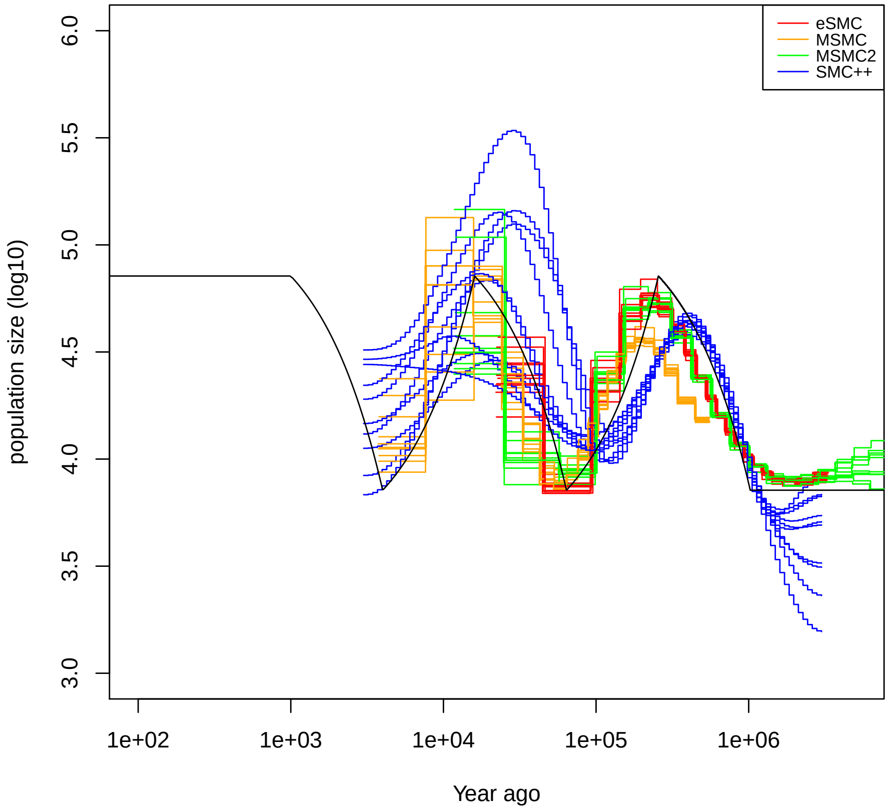


Figure 12: **Estimated demography of SMC method under sawtooth scenario.** Estimated demographic history under an sawtooth scenario with 10 replicates using simulated sequences. 2 sequences of 100 Mb for eSMC and MSMC2 (respectively in red and green ). We use 4 sequences of 100 Mb for MSMC (orange) and 20 sequences of 100 Mb for SMC++ (blue). Recombination rate is set to $1 \times 10^{-8}$ per generation per bp and mutation rate to $1.25 \times 10^{-8}$ per generation per bp. The simulated demographic history is represented in black.

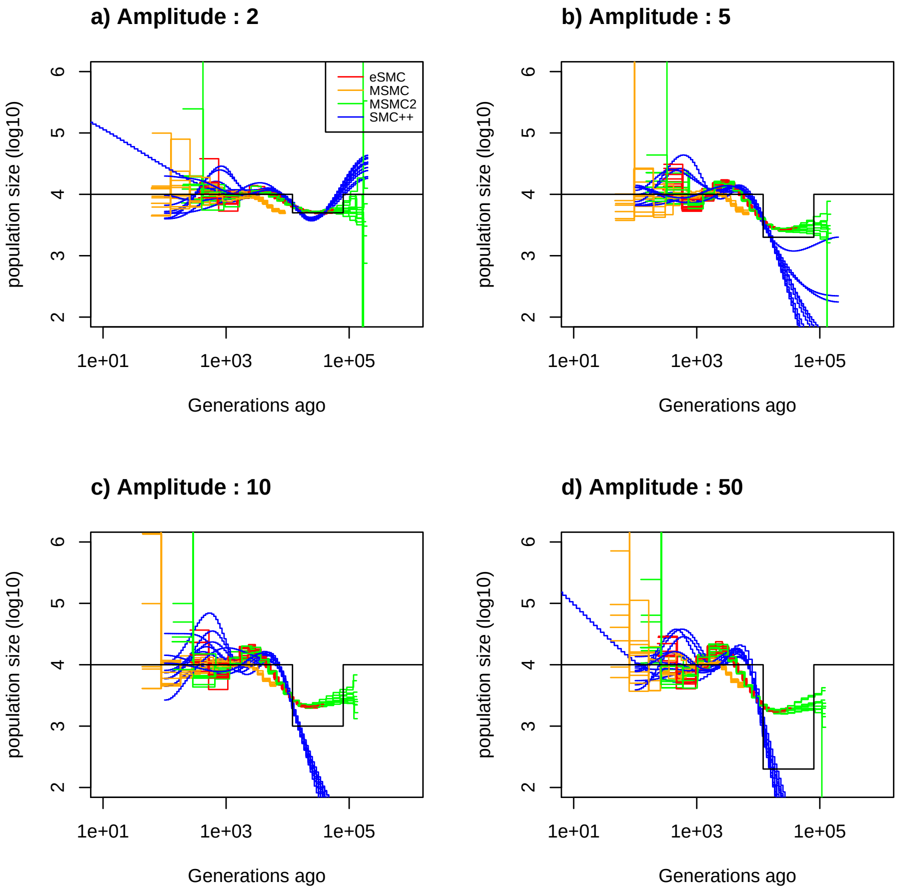


Figure 13: **Estimated demography of SMC method under a bottleneck scenario.** Estimated demographic history under a bottleneck scenario with 10 replicates using simulated sequences. 2 sequences of 100 Mb for eSMC and MSMC2 (respectively in red and green ). We use 4 sequences of 100 Mb for MSMC (orange) and 20 sequences of 100 Mb for SMC++ (blue). Recombination rate is set to $1 \times 10^{-8}$ per generation per bp and mutation rate to $1.25 \times 10^{-8}$ per generation per bp. The simulated demographic history is represented in black. a) Demographic history simulated under a bottleneck scenario of strength 2. b) Demographic history simulated under a bottleneck scenario of strength 5. c) Demographic history simulated under a bottleneck scenario of strength 10. d) Demographic history simulated under a bottleneck scenario of strength 50.

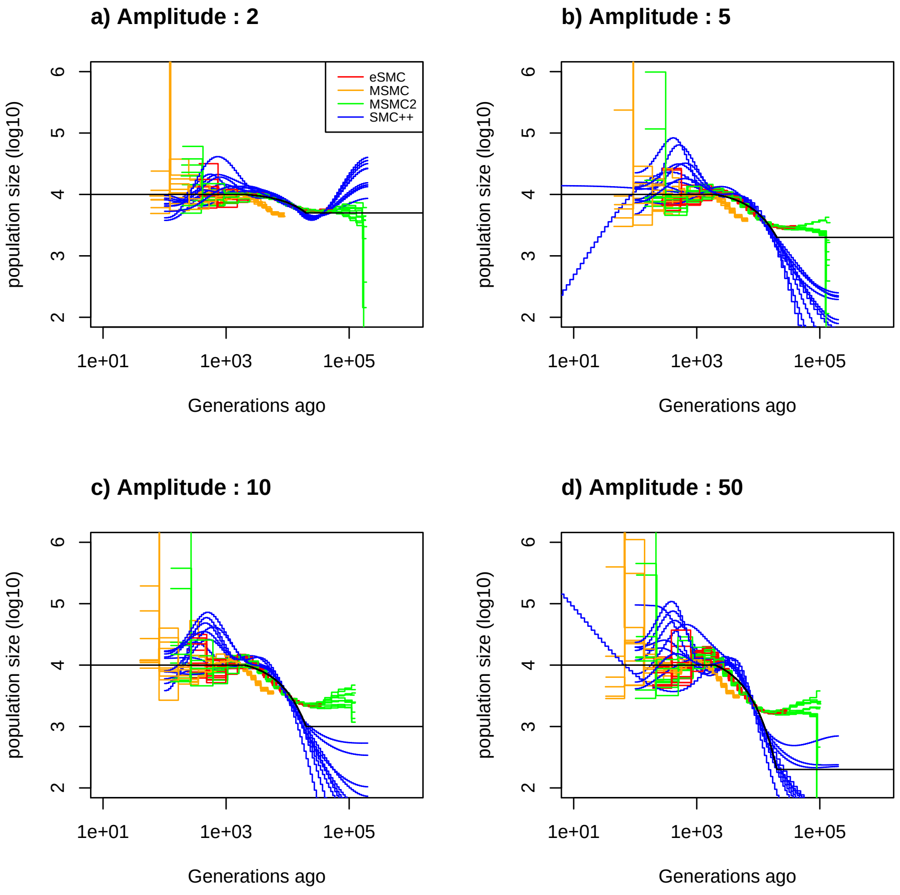


Figure 14: **Estimated demography of SMC method under a expansion scenario.** Estimated demographic history under an expansion scenario with 10 replicates using simulated sequences. 2 sequences of 100 Mb for eSMC and MSMC2 (respectively in red and green ). We use 4 sequences of 100 Mb for MSMC (orange) and 20 sequences of 100 Mb for SMC++ (blue). Recombination rate is set to $1 \times 10^{-8}$ per generation per bp and mutation rate to $1.25 \times 10^{-8}$ per generation per bp. The simulated demographic history is represented in black. a) Demographic history simulated under an expansion scenario of strength 2. b) Demographic history simulated under an expansion scenario of strength 5. c) Demographic history simulated under an expansion scenario of strength 10. d) Demographic history simulated under an expansion scenario of strength 50.

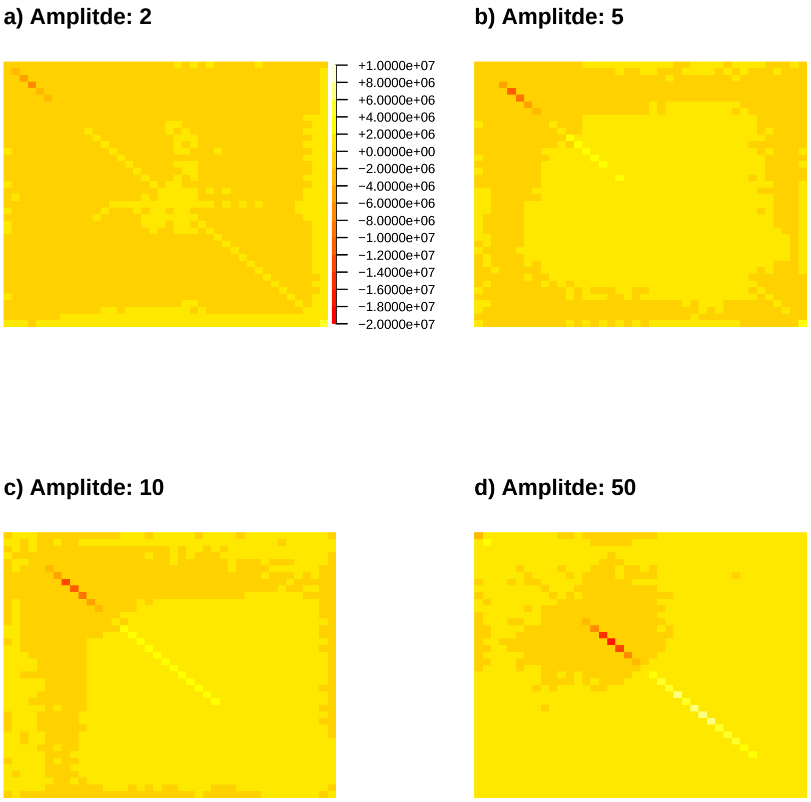


Figure 15: **Mean difference between estimated and actual transition matrix in saw-tooth scenario** Mean difference between estimated and transition matrix directly build from genealogy using sequences of 100 Mb under a saw-tooth scenario of strength 2,5,10 and 50 (respectively in a), b) c) and d) ) each with 10 replicates. Recombination rate is set to $1 \times 10^{-8}$ per generation per bp and mutation rate to $1.25 \times 10^{-8}$ per generation per bp.

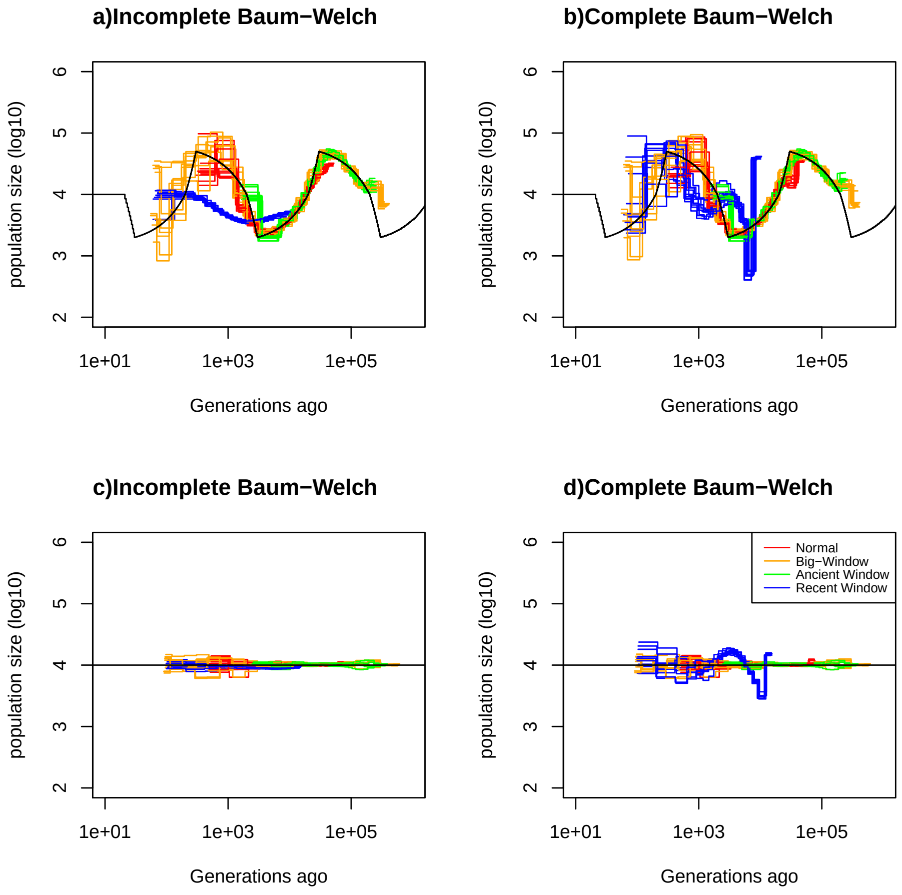


Figure 16: **Estimated demography using different window and optimization function under a saw-tooth and constant population size scenario.** Estimated demographic history under a saw-tooth (a), b)) and constant population size (c,d)) scenario with 10 replicates using 4 simulated sequences of length 50 Mb. Estimation with the incomplete optimization function are displayed in a) and c). Estimation with the complete Baum-Welch algorithm are displayed in b) and c). Recombination rate is set to $1 \times 10^{-8}$ per generation per bp and mutation rate to $1.25 \times 10^{-8}$ per generation per bp. The simulated demographic history is represented in black. Estimation with the time window of PSMC' are displayed in red, with the time window of MSMC2 in orange, estimation with a PSMC' time window shifted by fold 5 in past are displayed in green and shifted by fold 5 in recent time are displayed in blue.

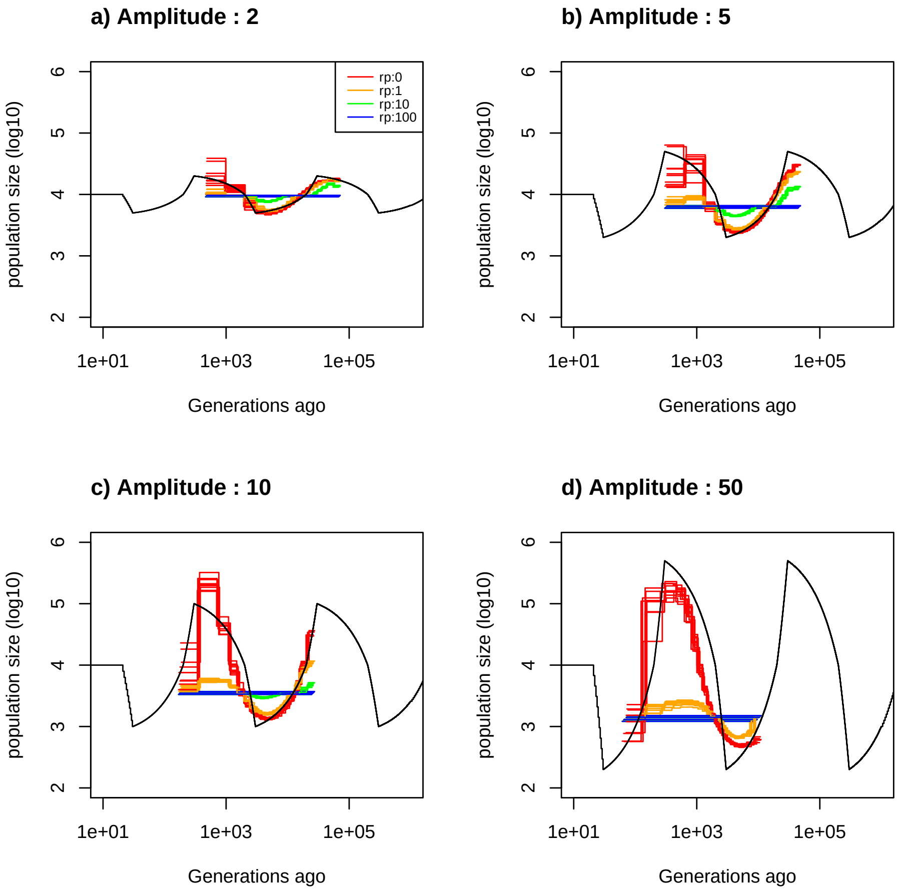


Figure 17: **Estimated demography by eSMC with regularization penalty.** Estimated demographic history under a saw-tooth scenario for different amplitudes of size change: a) 2-fold, b) 5-fold, c) 10-fold, and d) 50-fold. with 10 replicates using 2 simulated sequences of length 100 Mb. Estimation with no regularization is represented in red and with regularization penalty parameter 1,10,100 in orange, blue , green respectively.

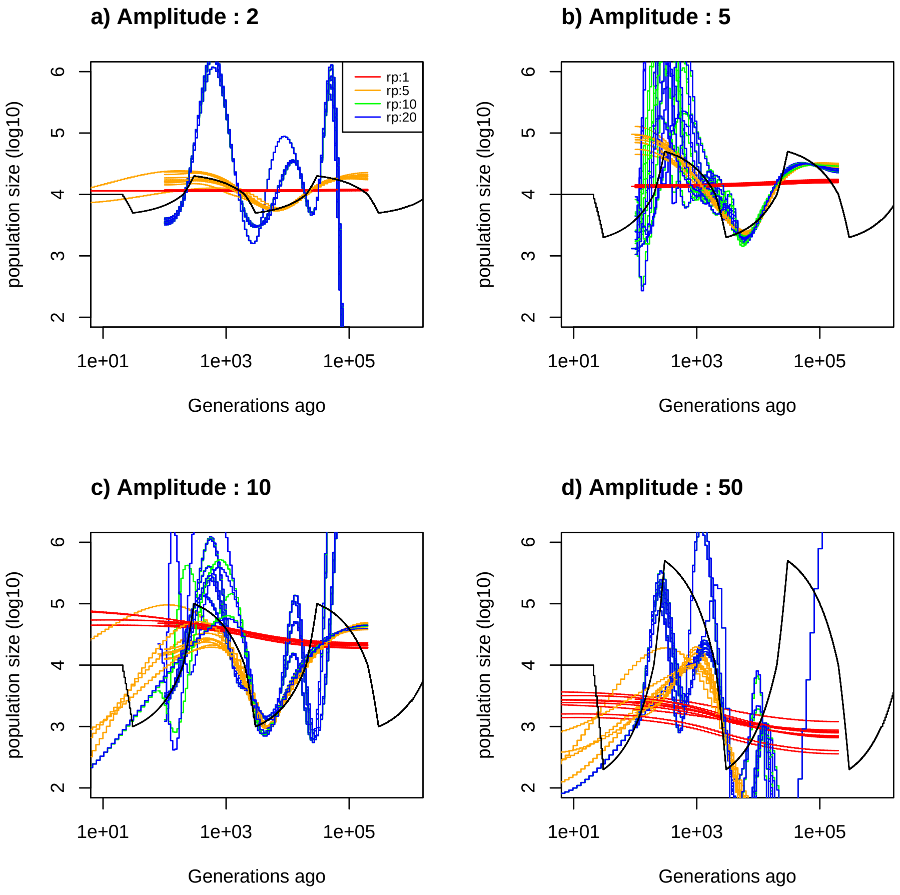


Figure 18: **Estimated demography by SMC++ with regularization penalty.** Estimated demographic history under a saw-tooth scenario for different amplitudes of size change: a) 2-fold, b) 5-fold, c) 10-fold, and d) 50-fold. with 10 replicates using 20 simulated sequences of length 100 Mb. Estimation with regularization penalty parameter 1,5,10,20 in red, orange, blue , green respectively.

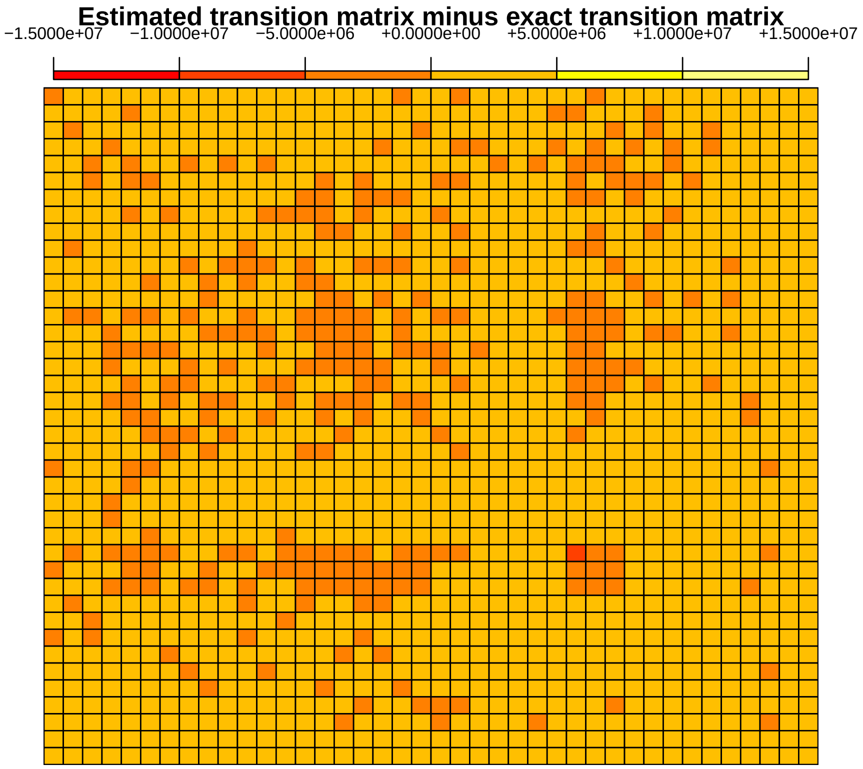


Figure 19: **Distance between estimated and exact transition matrix in bottleneck scenario** Mean distance between estimated and actual PSMC' transition matrix using 2 sequences of 100 Mb under a bottleneck scenario of strength 10 each with 10 replicates. Recombination rate is set to $1.25 \times 10^{-9}$ per generation per bp and mutation rate to $1.25 \times 10^{-8}$ per generation per bp.

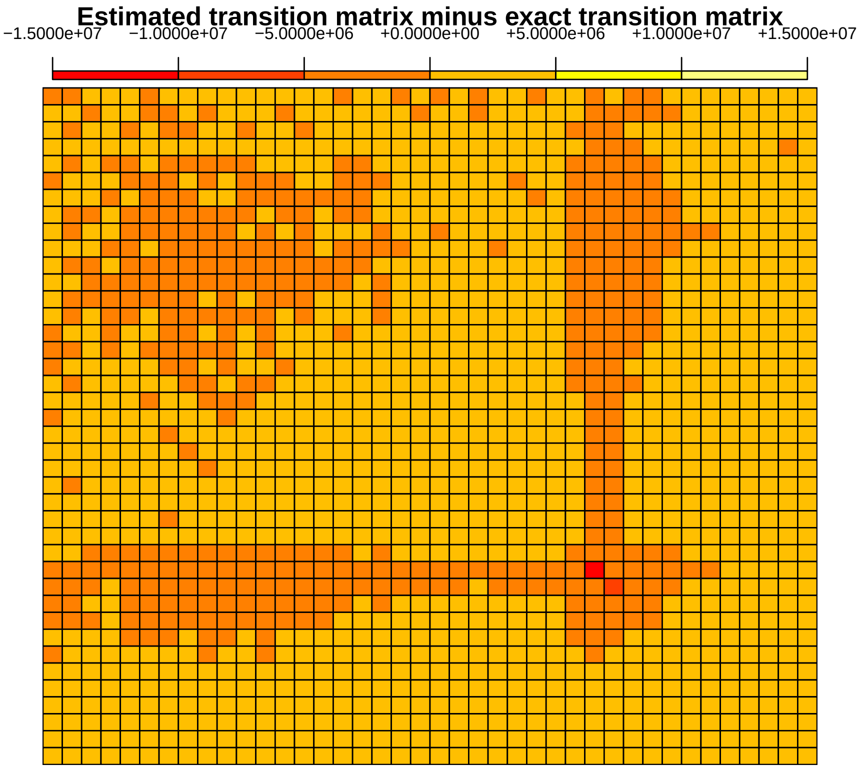


Figure 20: **Distance between estimated and exact transition matrix in bottleneck scenario** Mean distance between estimated and actual PSMC' transition matrix using 2 sequences of 100 Mb under a bottleneck scenario of strength 10 each with 10 replicates. Recombination rate is set to $1.25 \times 10^{-8}$ per generation per bp and mutation rate to $1.25 \times 10^{-8}$ per generation per bp.

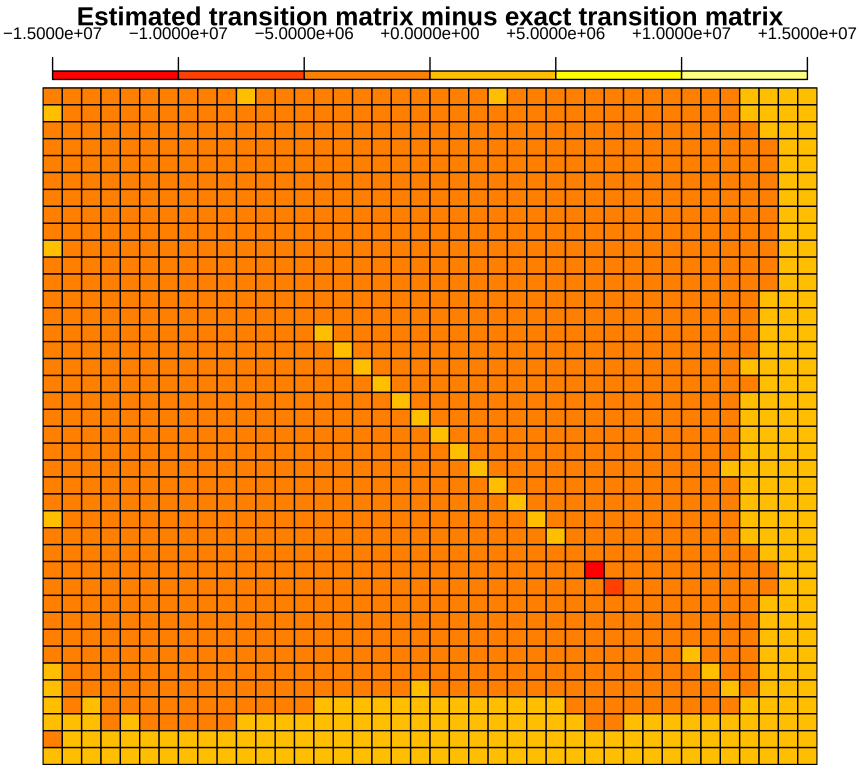


Figure 21: **Distance between estimated and exact transition matrix in bottleneck scenario** Mean distance between estimated and actual PSMC' transition matrix using 2 sequences of 100 Mb under a bottleneck scenario of strength 10 (respectively in a), b) c) and d) ) each with 10 replicates. Recombination rate is set to $1.25 \times 10^{-7}$ per generation per bp and mutation rate to $1.25 \times 10^{-8}$ per generation per bp.

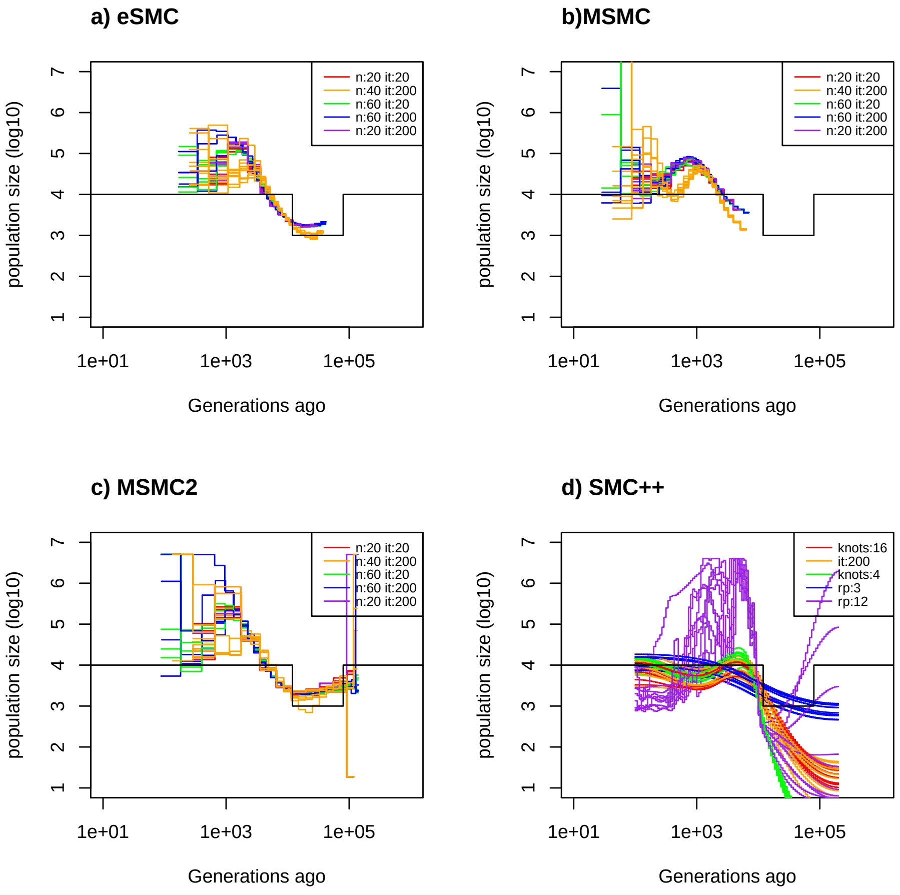


Figure 22: **Estimated demography of SMC method under a bottleneck scenario.** Estimated demographic history under a bottleneck scenario with 10 replicates using simulated sequences. 2 sequences of 100 Mb for eSMC and MSMC2 (respectively in a) and b) ). We use 4 sequences of 100 Mb for MSMC (c ) ) and 20 sequences of 100 Mb for SMC++ ( d) ). Recombination rate is set to $1.25 \times 10^{-7}$ per generation per bp and mutation rate to $1.25 \times 10^{-8}$ per generation per bp. Demographic history is simulated under a bottleneck scenario of strength 10 and is represented in black. Analysis with eSMC, MSMC and MSMC2 using 20 hidden states are in red, 200 iterations orange, 60 hidden states green, 60 hidden states and 200 iterations in blue, 20 hidden states and 200 iterations in purple. For SMC++, analysis using 16 knots are in red, 200 iterations in orange, 4 knots in green, regularization penalty set to 3 in blue and regularization-penalty set to 12 in purple.

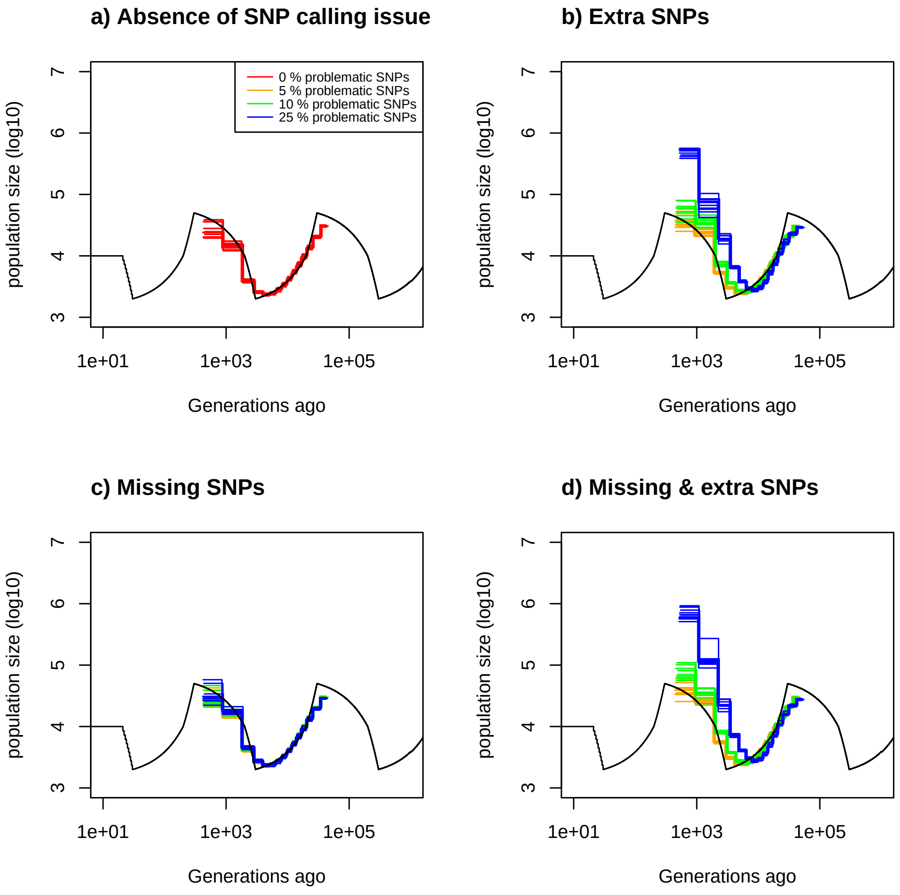


Figure 23: **Estimated demography of eSMC under a saw-tooth scenario with SNPs errors.** Estimated demographic history by eSMC under a saw-tooth scenario with 10 replicates using simulated sequences. 4 sequences of 100 Mb. Recombination rate is set to $1.25 \times 10^{-8}$ per generation per bp and mutation rate to $1.25 \times 10^{-8}$ per generation per bp. The simulated demographic history is represented in black. a) Demographic history simulated with ibsence of SNP calling issue (red). b) Demographic history simulated with 5% (orange),10% (green) and 25% (blue) missing SNPs. c) Demographic history simulated with 5% (orange),10% (green) and 25% (blue) additional SNPs. d) Demographic history simulated with 5% (orange),10% (green) and 25% (blue) of additional and missing SNPs.

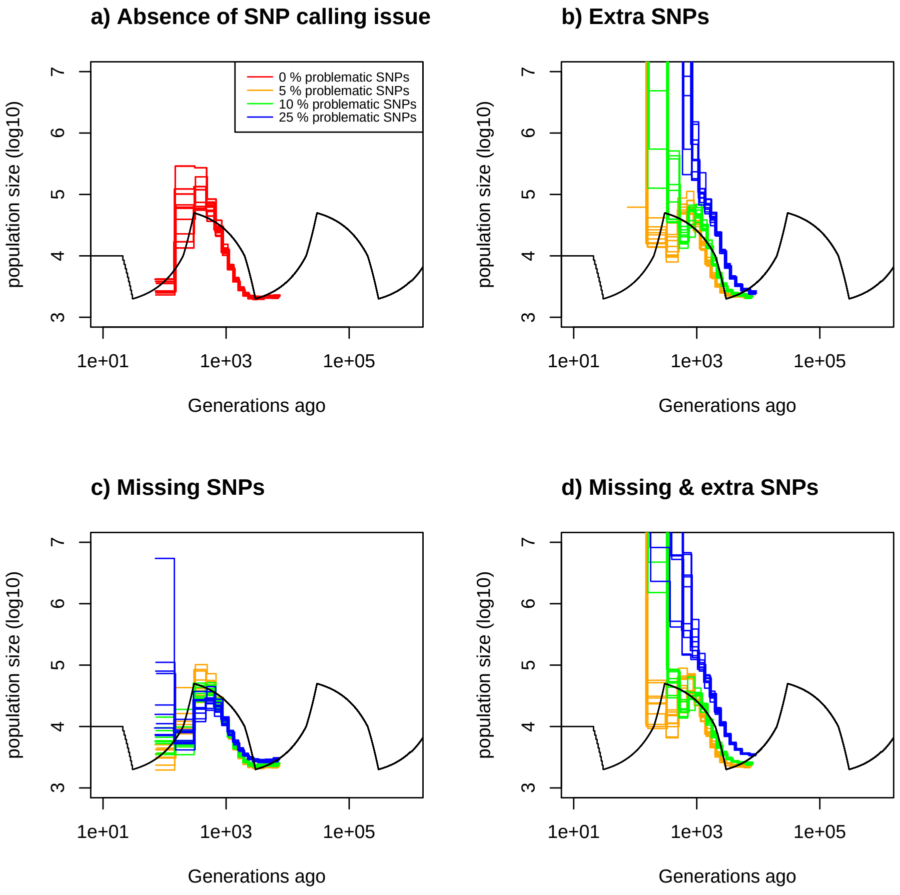


Figure 24: **Estimated demography of MSMC under a saw-tooth scenario with SNPs errors.** Estimated demographic history by eSMC under a saw-tooth scenario with 10 replicates using simulated sequences. 4 sequences of 100 Mb. Recombination rate is set to $1.25 \times 10^{-8}$ per generation per bp and mutation rate to $1.25 \times 10^{-8}$ per generation per bp. The simulated demographic history is represented in black. a) Demographic history simulated with ibsence of SNP calling issue (red). b) Demographic history simulated with 5% (orange),10% (green) and 25% (blue) missing SNPs. c) Demographic history simulated with 5% (orange),10% (green) and 25% (blue) additional SNPs. d) Demographic history simulated with 5% (orange),10% (green) and 25% (blue) of additional and missing SNPs .

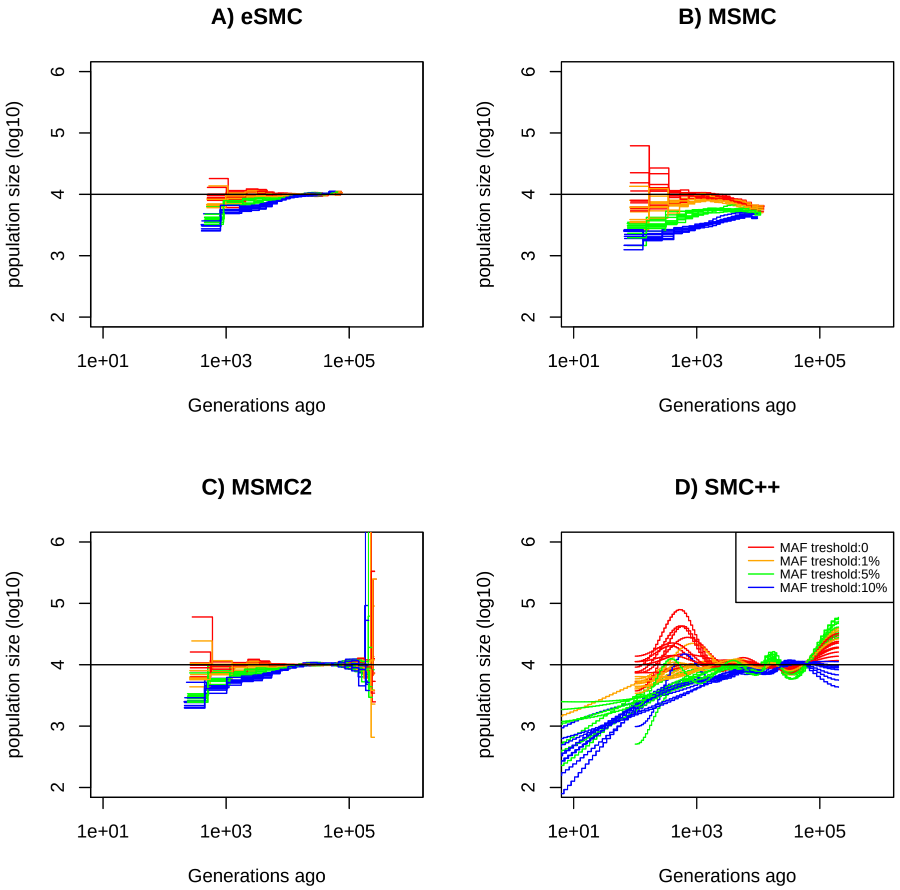


Figure 25: **Estimated demography of SMC method under a constant population size scenario.** Estimated demographic history under a constant population size with 10 replicates using simulated sequences. 2 sequences of 100 Mb for eSMC a) and MSMC2 b) . We use 4 sequences of 100 Mb for MSMC c) and 20 sequences of 100 Mb for SMC++ d). Recombination rate is set to $1 \times 10^{-8}$ per generation per bp and mutation rate to $1.25 \times 10^{-8}$ per generation per bp. The simulated demographic history is represented in black. One SNPs above a Minor Allele Frequency threshold are kept. Results with no filtering are represented in red, with threshold 1%, 5%, 10% respectively in orage, blue and green.

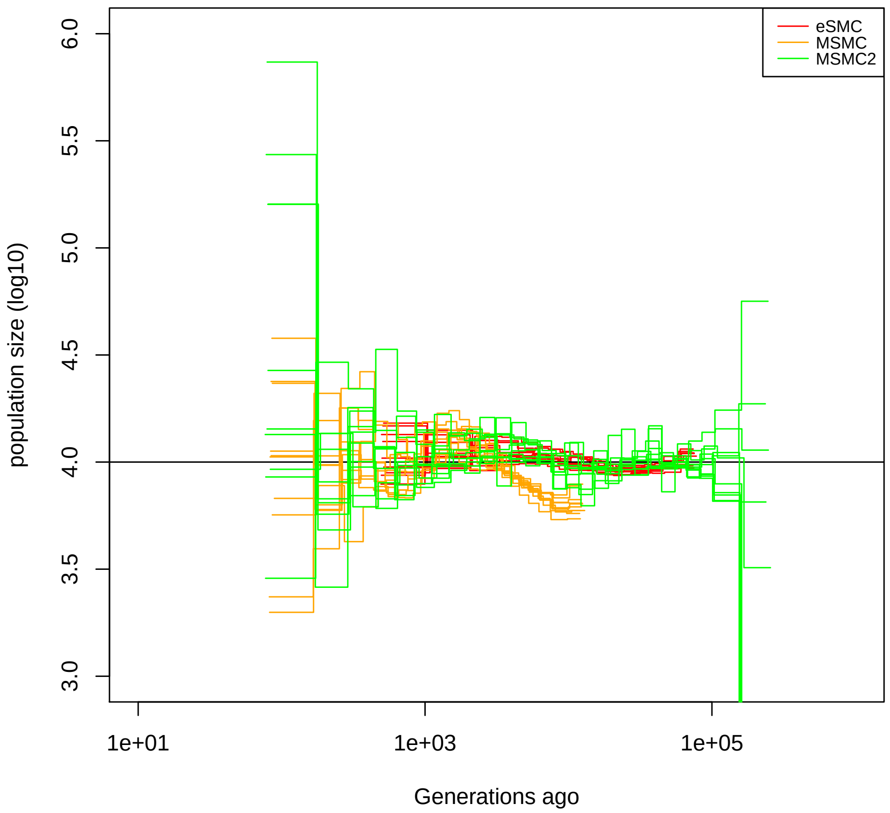


Figure 26: **Estimated demography of eSMC under a constant population size with recombination rate variation.** Estimated demographic history by eSMC under constant population size (black) with (red) or without (orange) variation of recombination rate along the sequence with 10 replicates using 2 simulated sequences of 40 Mb. Mutation rate is set to $1.25 \times 10^{-8}$ per generation per bp. Recombination rate changes randomly every 2 Mb between $2.5 \times 10^{-9}$ and $6.25 \times 10^{-8}$. T

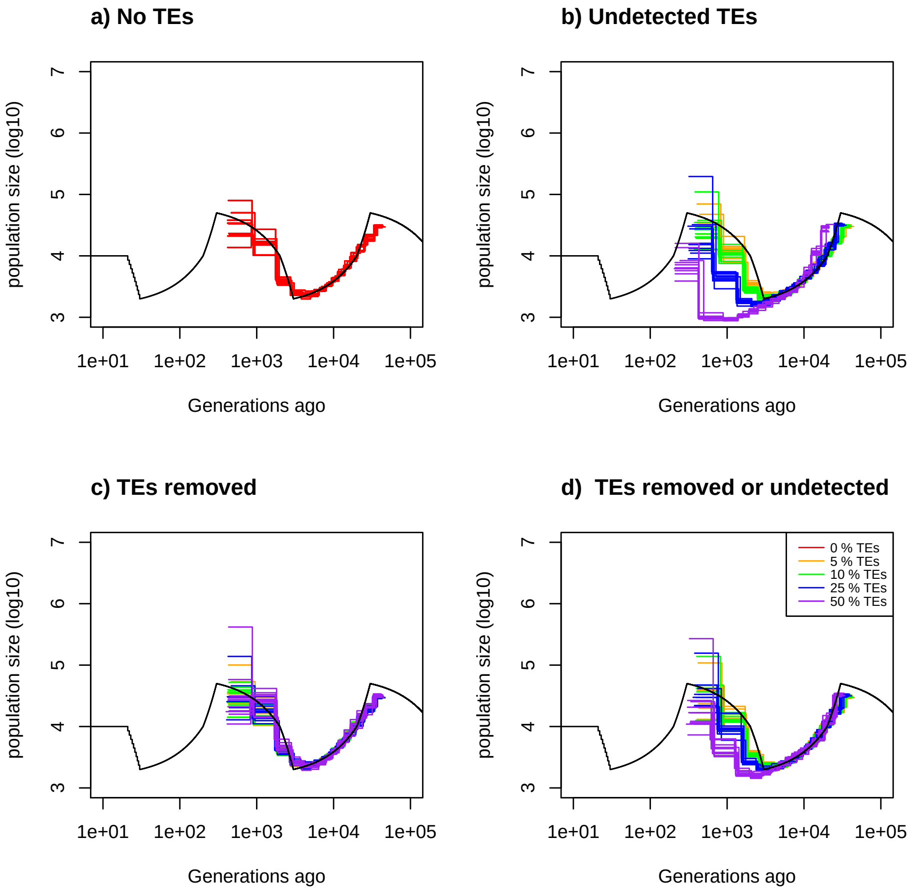


Figure 27: **Estimated demography of eSMC under a saw-tooth scenario with transposable elements.** Estimated demographic history by eSMC under a saw-tooth scenario with 10 replicates using simulated sequences. 4 sequences of 20 Mb. Recombination rate is set to $1.25 \times 10^{-8}$ per generation per bp and mutation rate to $1.25 \times 10^{-8}$ per generation per bp . The simulated demographic history is represented in black. Here tansposable element are of length 10 kbp. a) Demographic history simulated with no transposable elements. b) Demographic history simulated where transposable element are removed. c) Demographic history simulated where SNPs on transposable are removed. d) Demographic history simulated where half of transposable are removed and SNPs on the other half are removed. Proportion of transpobable element of the genome is set to 0% (red), 5% (orange), 10% (green), 25 % (blue) and 50 % (purple).

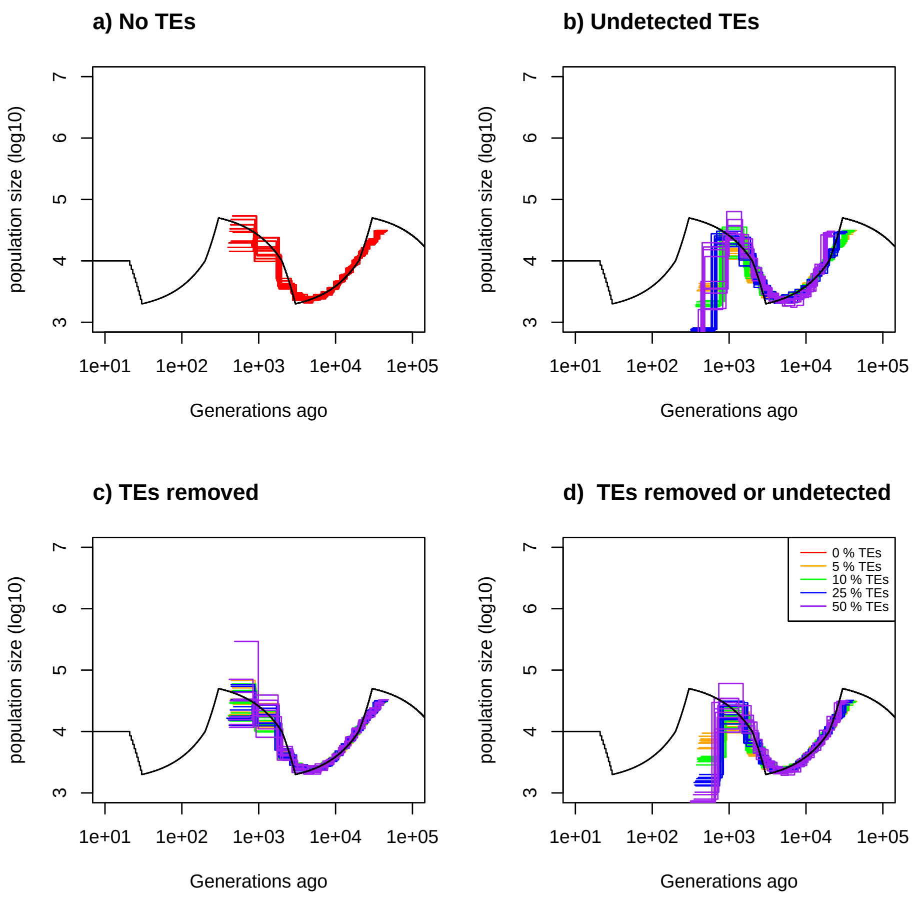


Figure 28: **Estimated demography of eSMC under a saw-tooth scenario with transposable elements.** Estimated demographic history by eSMC under a saw-tooth scenario with 10 replicates using simulated sequences. 4 sequences of 20 Mb. Recombination rate is set to $1.25 \times 10^{-8}$ per generation per bp and mutation rate to $1.25 \times 10^{-8}$ per generation per bp . The simulated demographic history is represented in black. Here tansposable element are of length 100 kbp. a) Demographic history simulated with no transposable elements. b) Demographic history simulated where transposable element are removed. c) Demographic history simulated where SNPs on transposable are removed. d) Demographic history simulated where half of transposable are removed and SNPs on the other half are removed. Proportion of transpobable element of the genome is set to 0% (red), 5% (orange), 10% (green), 25 % (blue) and 50 % (purple).

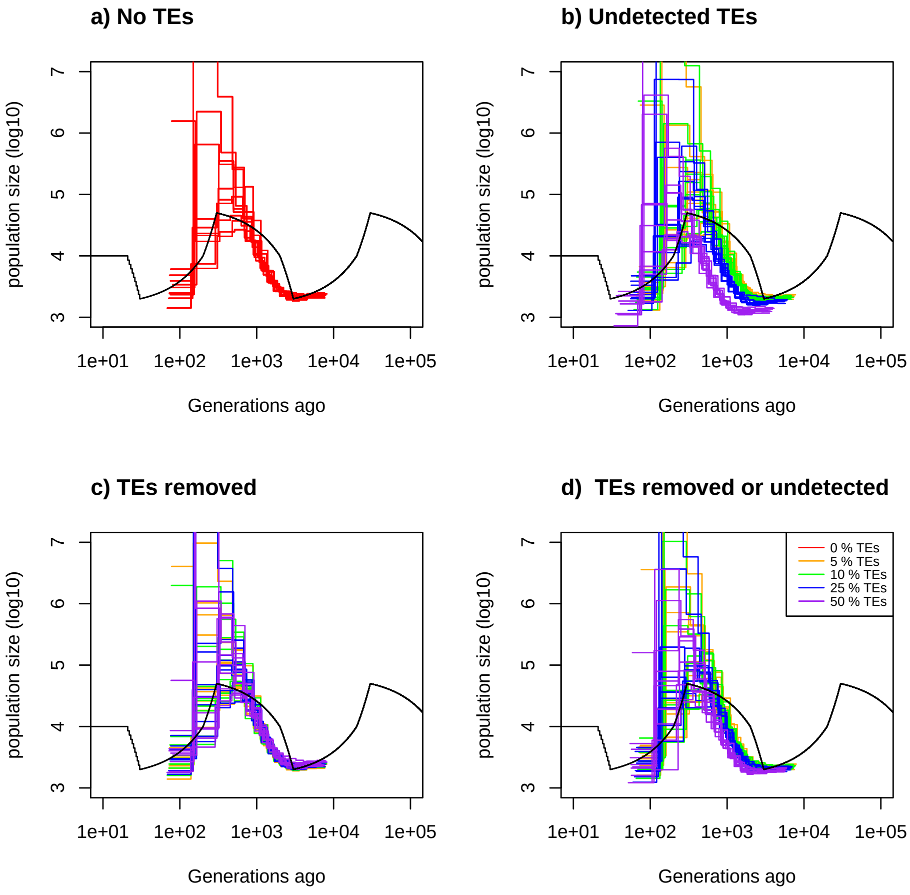


Figure 29: **Estimated demography of MSMC under a saw-tooth scenario with transposable elements.** Estimated demographic history by MSMC under a saw-tooth scenario with 10 replicates using simulated sequences. 4 sequences of 20 Mb. Recombination rate is set to $1.25 \times 10^{-8}$ per generation per bp and mutation rate to $1.25 \times 10^{-8}$ per generation per bp . The simulated demographic history is represented in black. Here tansposable element are of length 1kbp. a) Demographic history simulated with no transposable elements. b) Demographic history simulated where transposable element are removed. c) Demographic history simulated where SNPs on transposable are removed. d) Demographic history simulated where half of transposable are removed and SNPs on the other half are removed. Proportion of transpobable element of the genome is set to 0% (red), 5% (orange), 10% (green), 25 % (blue) and 50 % (purple).

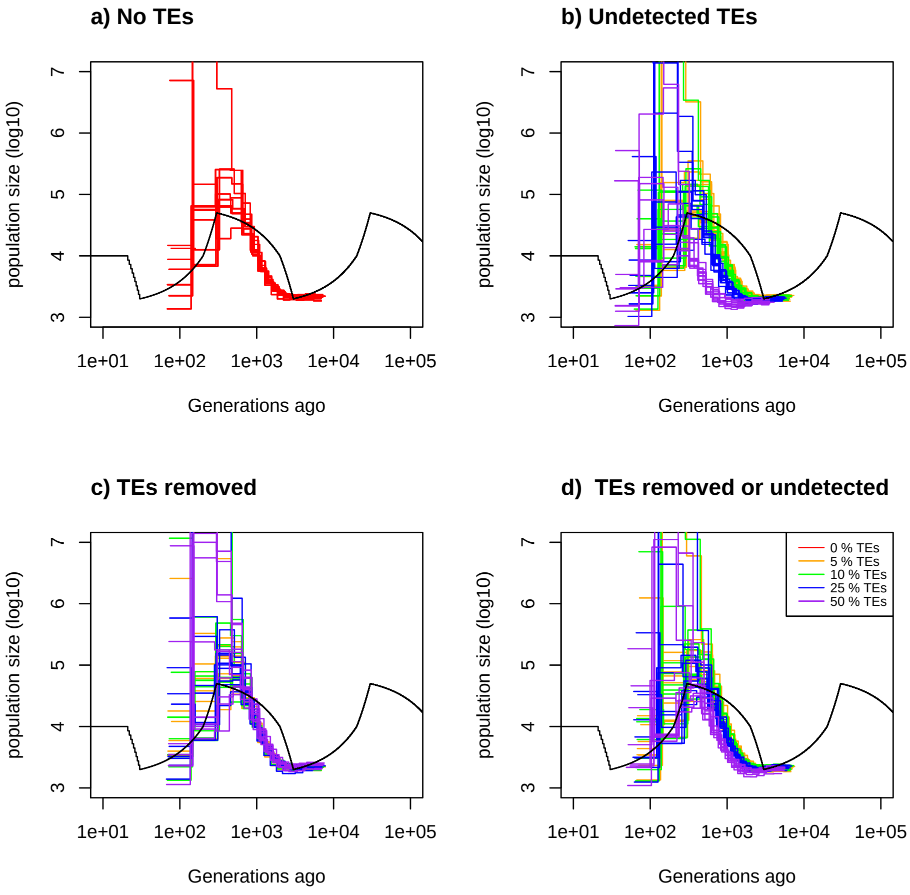


Figure 30: **Estimated demography of MSMC under a saw-tooth scenario with transposable elements.** Estimated demographic history by MSMC under a saw-tooth scenario with 10 replicates using simulated sequences. 4 sequences of 20 Mb. Recombination rate is set to $1.25 \times 10^{-8}$ per generation per bp and mutation rate to $1.25 \times 10^{-8}$ per generation per bp . The simulated demographic history is represented in black. Here tansposable element are of length 10kbp. a) Demographic history simulated with no transposable elements. b) Demographic history simulated where transposable element are removed. c) Demographic history simulated where SNPs on transposable are removed. d) Demographic history simulated where half of transposable are removed and SNPs on the other half are removed. Proportion of transpobable element of the genome is set to 0% (red), 5% (orange), 10% (green), 25 % (blue) and 50 % (purple).

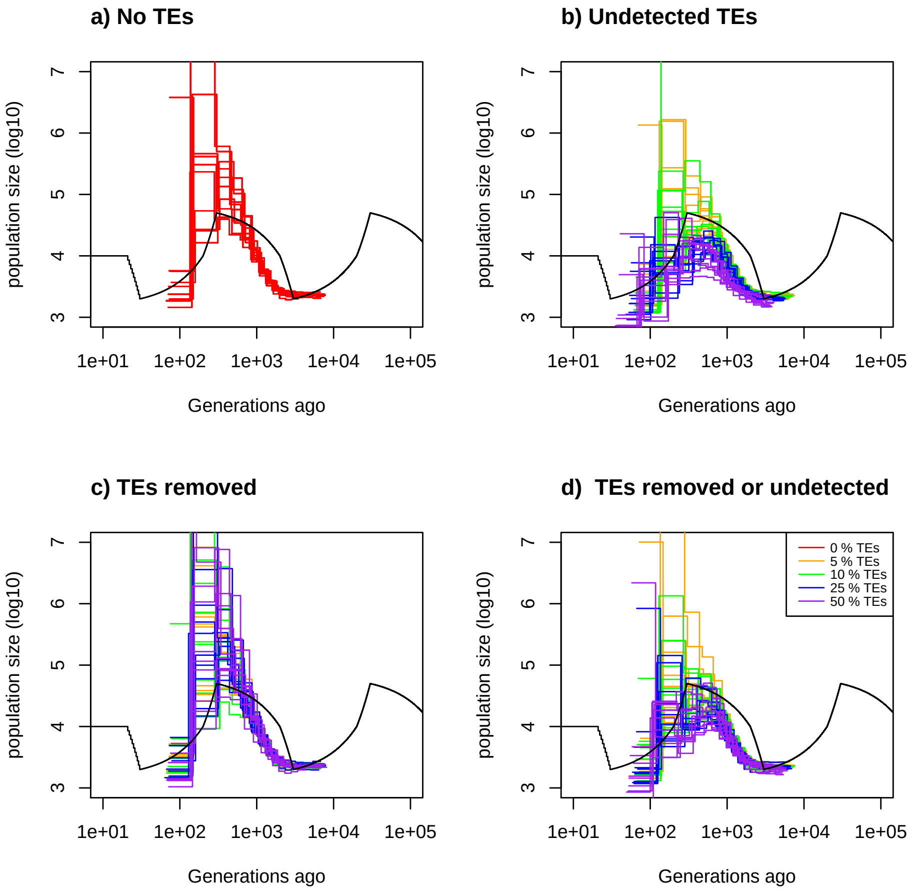


Figure 31: **Estimated demography of MSMC under a saw-tooth scenario with transposable elements.** Estimated demographic history by MSMC under a saw-tooth scenario with 10 replicates using simulated sequences. 4 sequences of 20 Mb. Recombination rate is set to $1.25 \times 10^{-8}$ per generation per bp and mutation rate to $1.25 \times 10^{-8}$ per generation per bp . The simulated demographic history is represented in black. Here tansposable element are of length 100kbp. a) Demographic history simulated with no transposable elements. b) Demographic history simulated where transposable element are removed. c) Demographic history simulated where SNPs on transposable are removed. d) Demographic history simulated where half of transposable are removed and SNPs on the other half are removed. Proportion of transpobable element of the genome is set to 0% (red), 5% (orange), 10% (green), 25 % (blue) and 50 % (purple).

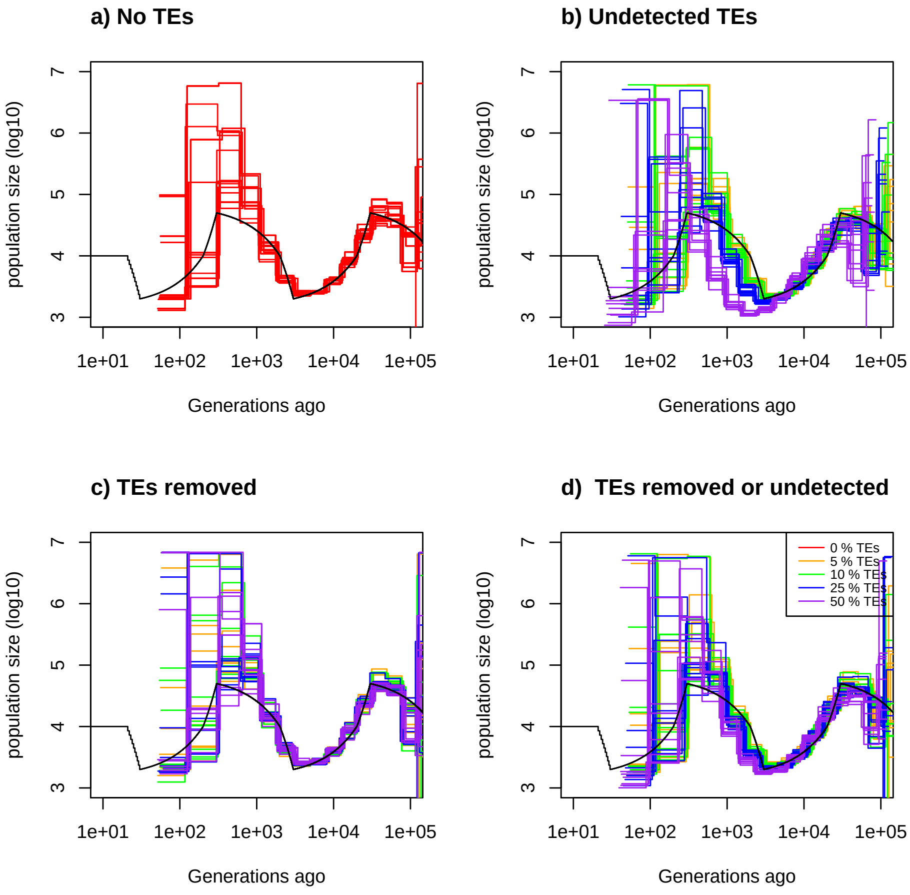


Figure 32: **Estimated demography of MSMC2 under a saw-tooth scenario with transposable elements.** Estimated demographic history by MSMC2 under a saw-tooth scenario with 10 replicates using simulated sequences. 4 sequences of 20 Mb. Recombination rate is set to $1.25 \times 10^{-8}$ per generation per bp and mutation rate to $1.25 \times 10^{-8}$ per generation per bp . The simulated demographic history is represented in black. Here tansposable element are of length 1kbp. a) Demographic history simulated with no transposable elements. b) Demographic history simulated where transposable element are removed. c) Demographic history simulated where SNPs on transposable are removed. d) Demographic history simulated where half of transposable are removed and SNPs on the other half are removed. Proportion of transpobable element of the genome is set to 0% (red), 5% (orange), 10% (green), 25 % (blue) and 50 % (purple).

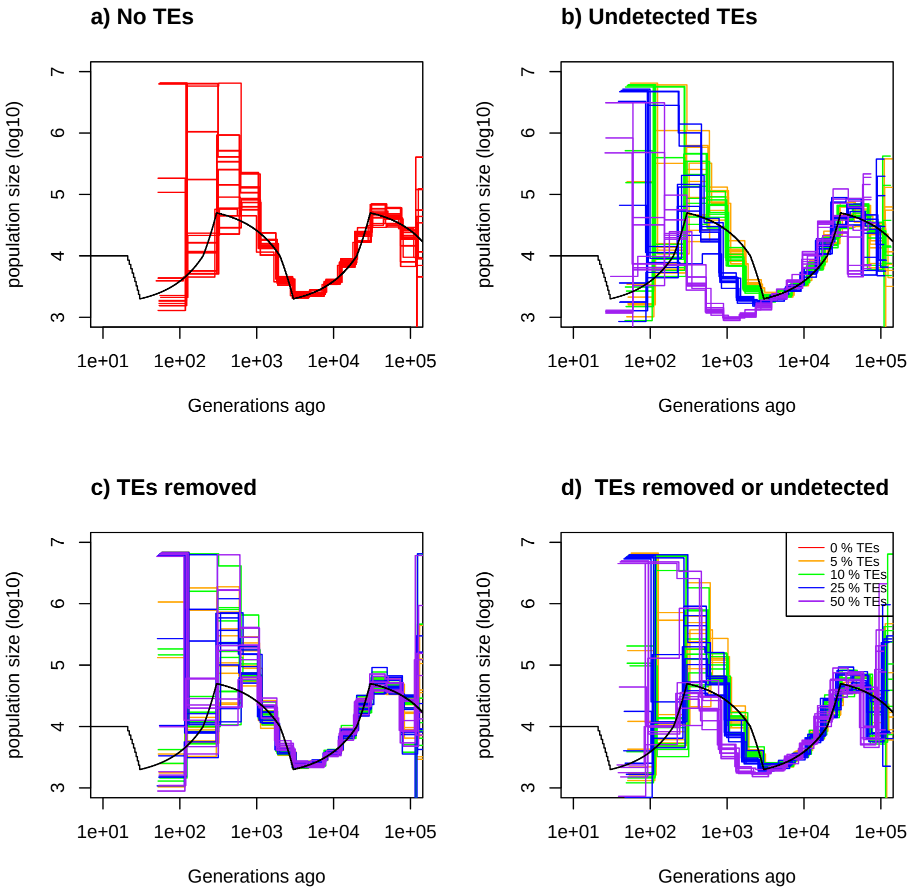


Figure 33: **Estimated demography of MSMC2 under a saw-tooth scenario with transposable elements.** Estimated demographic history by MSMC2 under a saw-tooth scenario with 10 replicates using simulated sequences. 4 sequences of 20 Mb. Recombination rate is set to $1.25 \times 10^{-8}$ per generation per bp and mutation rate to $1.25 \times 10^{-8}$ per generation per bp . The simulated demographic history is represented in black. Here tansposable element are of length 10kbp. a) Demographic history simulated with no transposable elements. b) Demographic history simulated where transposable element are removed. c) Demographic history simulated where SNPs on transposable are removed. d) Demographic history simulated where half of transposable are removed and SNPs on the other half are removed. Proportion of transpobable element of the genome is set to 0% (red), 5% (orange), 10% (green), 25 % (blue) and 50 % (purple).

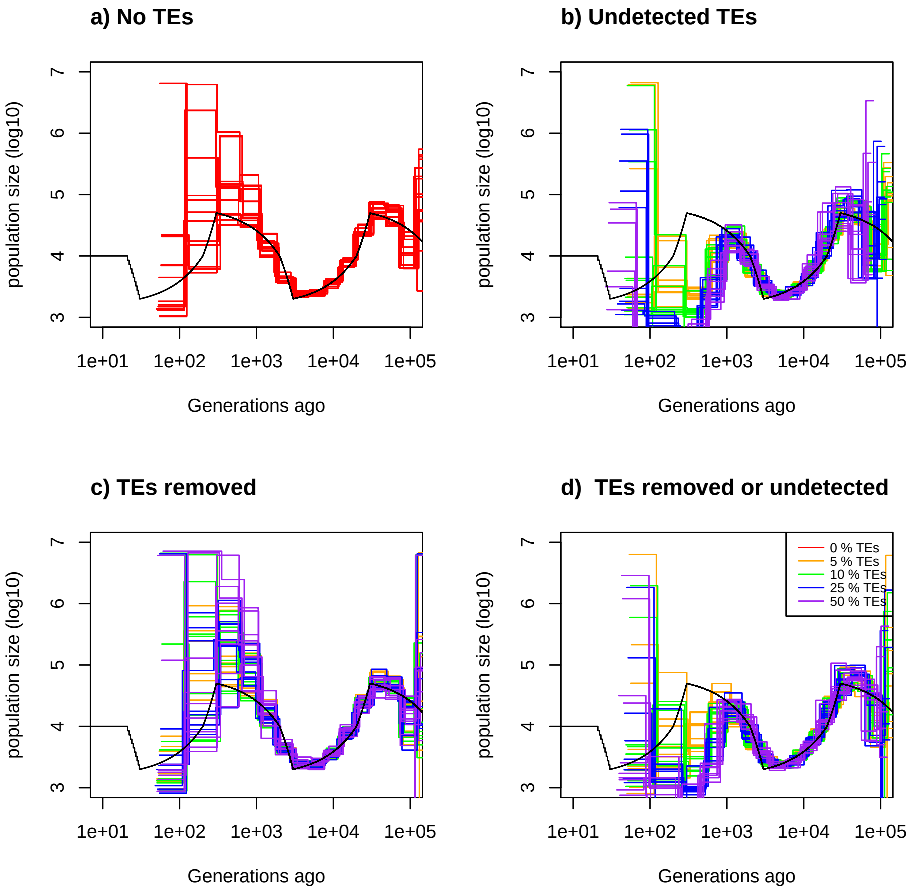


Figure 34: **Estimated demography of MSMC2 under a saw-tooth scenario with transposable elements.** Estimated demographic history by MSMC2 under a saw-tooth scenario with 10 replicates using simulated sequences. 4 sequences of 20 Mb. Recombination rate is set to $1.25 \times 10^{-8}$ per generation per bp and mutation rate to $1.25 \times 10^{-8}$ per generation per bp . The simulated demographic history is represented in black. Here tansposable element are of length 100 kbp. a) Demographic history simulated with no transposable elements. b) Demographic history simulated where transposable element are removed. c) Demographic history simulated where SNPs on transposable are removed. d) Demographic history simulated where half of transposable are removed and SNPs on the other half are removed. Proportion of transpobable element of the genome is set to 0% (red), 5% (orange), 10% (green), 25 % (blue) and 50 % (purple).

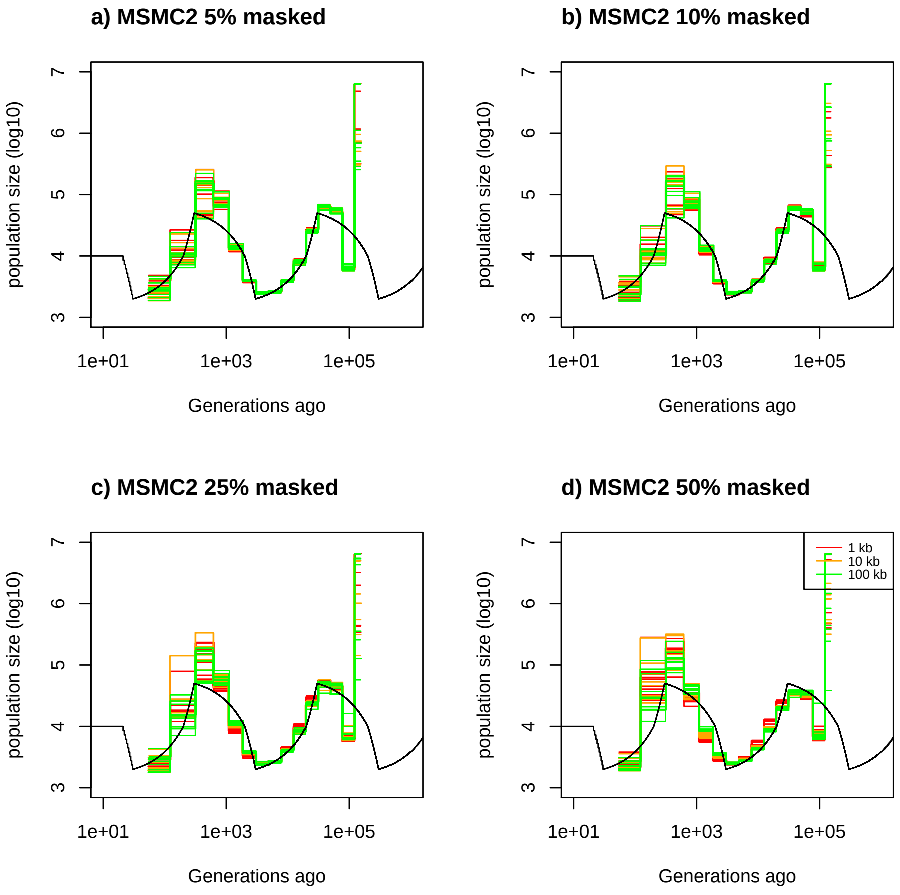


Figure 35: **Estimated demography of MSMC2 under a saw-tooth scenario with masked transposable elements.** Estimated demographic history by MSMC2 under a saw-tooth scenario with 10 replicates using simulated sequences. 4 sequences of 20 Mb. Recombination rate is set to $1.25 \times 10^{-8}$ per generation per bp and mutation rate to $1.25 \times 10^{-8}$ per generation per bp . The simulated demographic history is represented in black. Here tansposable element are of length 1kbp. a) Demographic history simulated with 5% transposable elements. b) DDemographic history simulated with 10% transposable elements. c) Demographic history simulated with 25% transposable elements. d) Demographic history simulated with 50% transposable elements. Size of transposable elements are set to 1kb (red), 10kb (orange), 100 kb (green).

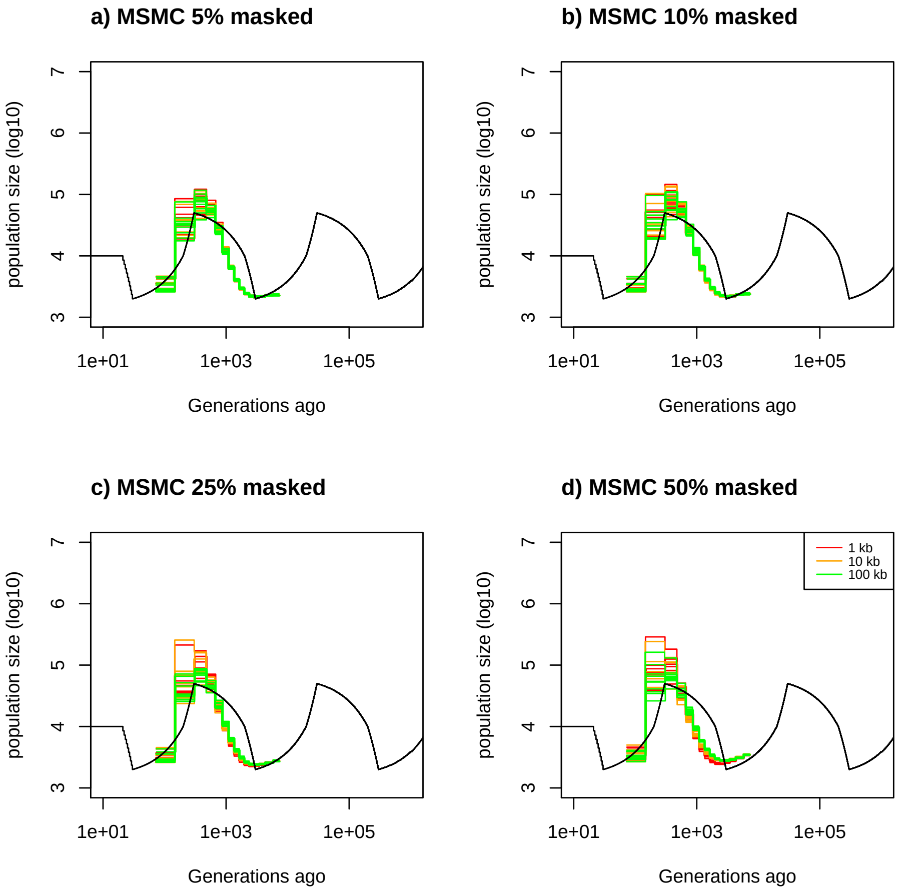


Figure 36: **Estimated demography of MSMC under a saw-tooth scenario with masked transposable elements.** Estimated demographic history by MSMC under a saw-tooth scenario with 10 replicates using simulated sequences. 4 sequences of 20 Mb. Recombination rate is set to $1.25 \times 10^{-8}$ per generation per bp and mutation rate to $1.25 \times 10^{-8}$ per generation per bp . The simulated demographic history is represented in black. Here tansposable element are of length 1kbp. a) Demographic history simulated with 5% transposable elements. b) DDemographic history simulated with 10% transposable elements. c) Demographic history simulated with 25% transposable elements. d) Demographic history simulated with 50% transposable elements. Size of transposable elements are set to 1kb (red), 10kb (orange), 100 kb (green).

### 2 Supplementary Tables

| Figure | eSMC 10 Mb | eSMC 100 Mb | eSMC 1 Gb | eSMC 10 Gb |
|---|---|---|---|---|
| 1 a) | 7.23 (0.11) | 6.34 (0.040) | 6.10 (0.01) | 5.99 (0.005) |
| 1 b) | 8.59 (0.099) | 7.99 (0.012) | 7.86 (0.010) | 7.86 (0.002) |
| 1 c) | 8.96 (0.038) | 8.74 (0.011) | 8.77 (0.005) | 8.76 (0.006) |
| 1 d) | 10.36 (0.002) | 10.37 (0.001) | 10.38 (0.001) | 10.38 (0.001) |

Table 1: Average mean square error of Figure 1 (in log10). The coefficient of variation is indicated in brackets.

| Figure | eSMC | MSMC | MSMC2 | SMC++ |
|---|---|---|---|---|
| 3 a) | 6.82 (0.07) | 7.60 (0.05) | 8.21 (0.17) | 7.57 (0.03) |
| 3 b) | 8.16 (0.01) | 8.69 (0.08) | 10.0 (0.16) | 7.84 (0.03) |
| 3 c) | 9.27 (0.03) | 9.53 (0.08) | 11.43 (0.08) | 8.86 (0.004) |
| 3 d) | 10.0 (0.008) | 10.90 (0.12) | 11.0 (0.04) | 10.46 (0.0001) |

Table 2: Average mean square error of Figure 3 (in log10). The coefficient of variation is indicated in brackets.

| Figure | $\frac{\rho}{\theta} = 0.1$ | $\frac{\rho}{\theta} = 1$ | $\frac{\rho}{\theta} = 10$ |
|---|---|---|---|
| 4 a) | 8.21 (0.12) | 7.45 (0.06) | 9.62 (0.04) |
| 4 b) | 15.7 (0.3) | 9.46 (0.33) | 10.43 (0.20) |
| 4 c) | 9.95 (0.24) | 8.48 (0.22) | 10.65 (0.13) |
| 4 d) | 7.78 (0.008) | 7.79 (0.003) | 7.79 (0.001) |

Table 3: Average mean square error of Figure 4 (in log10). The coefficient of variation is indicated in brackets.

| % of problematic SNPs | 5 a) | 5 b) | 5 c) | 5 d) |
|---|---|---|---|---|
| 0 | 9.10 (0.10) | | | |
| 5 | | 12.66 (0.002) | 13.0 (0.007) | 13.27 (0.001) |
| 10 | | 10.08 (0.11) | 12.15 (0.07) | 12.55 (0.01) |
| 25 | | 12.83 (0.012) | 13.03 (0.001) | 13.28 (0.006) |

Table 4: Average mean square error of Figure 5 (in log10). The coefficient of variation is indicated in brackets.

| Figure | Share | Not share |
|---|---|---|
| 6 a) | 6.58 (0.06) | 6.68 (0.08) |
| 6 b) | 7.99 (0.04) | 8.02 (0.098) |
| 6 c) | 7.08 (0.15) | 7.43 (0.093) |
| 6 d) | 7.99 (0.07) | 8.09 (0.09) |

Table 5: Average mean square error of Figure 6 (in log10). The coefficient of variation is indicated in brackets.

| % of problematic TEs | 7 a) | 5 b) | 5 c) | 5 d) |
|---|---|---|---|---|
| 0 | 8.04 (0.03) | | | |
| 5 | | 8.05 (0.029) | 8.05 (0.035) | 8.039 (0.03) |
| 10 | | 8.04 (0.07) | 8.06 (0.038) | 8.058 (0.035) |
| 25 | | 8.20 (0.018) | 8.08 (0.04) | 8.13 (0.022) |
| 50 | | 8.59 (0.006) | 8.33 (0.08) | 8.25 (0.016) |

Table 6: Average mean square error of Figure 7 (in log10). The coefficient of variation is indicated in brackets.
